# Analysis of ductal carcinoma in situ by self-reported race reveals molecular differences related to outcome

**DOI:** 10.1101/2023.09.28.560000

**Authors:** Siri H. Strand, Kathleen E. Houlahan, Vernal Branch, Thomas Lynch, Bryan Harmon, Fergus Couch, Kristalyn Gallagher, Mark Kilgore, Shi Wei, Angela DeMichele, Tari King, Priscilla McAuliffe, Christina Curtis, Kouros Owzar, Jeffrey R. Marks, Graham A. Colditz, E. Shelley Hwang, Robert B. West

## Abstract

Ductal carcinoma in situ (DCIS) is a non-obligate precursor to invasive breast cancer (IBC). Studies have indicated differences in DCIS outcome based on race or ethnicity, but molecular differences have not been investigated. We examined the molecular profile of DCIS by self-reported race (SRR) and outcome groups in Black (n=99) and White (n=191) women with DCIS in a large DCIS case-control cohort with longitudinal follow up. Gene expression and pathway analyses indicated that different genes and pathways are involved in ipsilateral breast outcome (DCIS or IBC) after DCIS treatment in White versus Black women. We identified differences in ER and HER2 expression, tumor microenvironment composition, and copy number variations by SRR and outcome groups. Our results suggest that different molecular mechanisms drive subsequent ipsilateral breast events in Black versus White women.

---

Ductal carcinoma in situ (DCIS) is a preinvasive neoplastic lesion of the breast that reflects increased risk for invasive breast cancer (IBC)^1^. DCIS encompasses a heterogenous group of lesions, the natural history of which is not well-understood. The incidence of DCIS in the U.S. has increased greatly over the past 40 years, mainly due to screening mammography^2^. Treatment for DCIS includes mastectomy or breast-conserving surgery with or without radiation or endocrine therapy. However, research indicates many women are over-diagnosed and over-treated, and trials on active surveillance cohorts are ongoing to better understand the factors that lead to DCIS disease progression^3–5^.

Several studies have demonstrated a significant variation in risk of developing IBC after DCIS by race or ethnicity. Overall, results indicate that Black women are more likely to have recurring invasive and noninvasive tumors in either breast, as well as overall higher IBC mortality rates after DCIS, compared to White women^6–12^. In these studies, racial or ethnic risk disparities could not be attributed to histologic features or treatment of DCIS^10,12^ though subsequent IBC after DCIS was biologically more aggressive among Black than White women^11^.

While race and ethnicity are social constructs, ancestry refers to a person’s genetic admixture reflecting ancestral lineage^13^. Studies have implicated ancestry-related molecular differences in invasive cancers, including IBC^14–17^, but molecular differences by race, ethnicity, or ancestry have not yet been investigated in DCIS or other precancers. We recently generated the Human Tumor Atlas Network (HTAN) DCIS Atlas^18,19^, where we identified molecular changes in DCIS at the DNA, RNA and protein levels. This work showed that many of the molecular alterations found in IBC are already present at the earlier DCIS stage, implying that any racial differences in IBC may arise at this precursor stage. In the present study, we leveraged the large amount of metadata available for two HTAN DCIS cohorts to analyze molecular features and DCIS outcome by self-reported race (SRR) and inferred global ancestry from DNA sequencing data to facilitate analysis by both ancestry and SRR.

## Results

### Outcome analysis by SRR

We examined the molecular profile of DCIS by SRR for 99 Black and 191 White women in the combined TBCRC and RAHBT cohorts (**Table 1**). While we previously analyzed these cohorts separately^18^, we here combined them to enhance statistical power and thus allow analysis by SRR. To investigate the correlation between SRR and genomic ancestry in our dataset, we inferred global ancestry using whole genome sequencing (WGS) data from the combined cohort and compared these to SRR for the 208 patients where both parameters were available. Similar to previous reports^20^, we found that African ancestry was highly concordant with Black SRR, and European ancestry with White SRR (**Extended Data fig. 1A**). Due to the high level of agreement between genomic ancestry and SRR, and because WGS data and thus ancestry was not available for the entire study population, the following analyses were carried out using SRR only.

**Figure 1:**
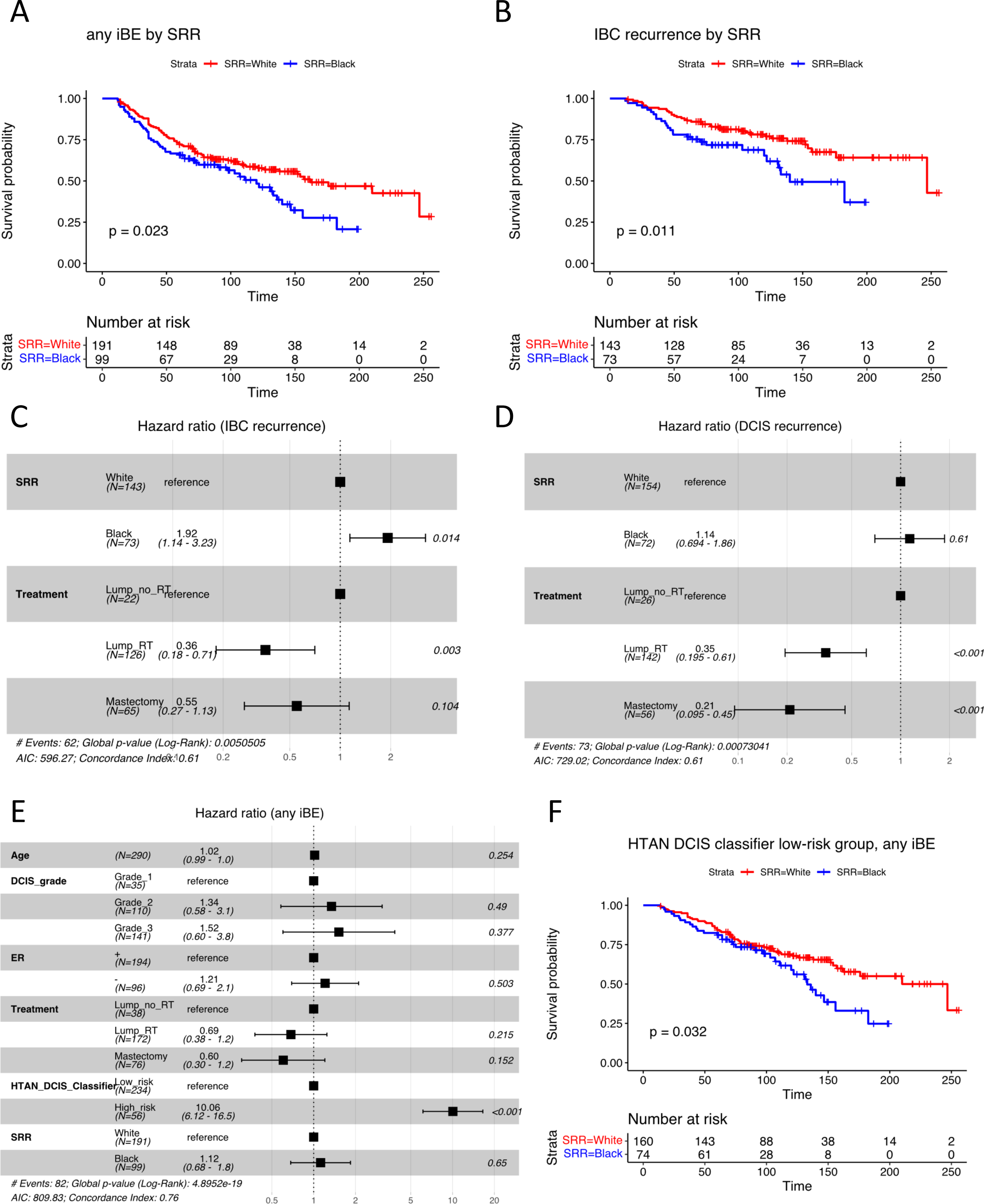
Outcome analysis by SRR. A) Kaplan-Meier plot of time to iBE (full follow-up) stratified by SRR. B) Kaplan-Meier plot of time to IBC recurrence only (full follow-up) stratified by SRR. C) Forest plot of multivariable Cox regression analysis including SRR and treatment type, for IBC recurrence only (full follow-up). D) Forest plot of multivariable Cox regression analysis including SRR and treatment type, for DCIS recurrence only (full follow-up). E) Forest plot of multivariable Cox regression analysis including the HTAN DCIS classifier, SRR, age at diagnosis, ER status, DCIS grade, and treatment, for any iBE (5-year outcome). F) Kaplan-Meier plot of time to iBE (full follow-up) in the HTAN DCIS classifier low-risk group stratified by SRR. A, B, F) P-values from log-rank tests.

**Table 1:**
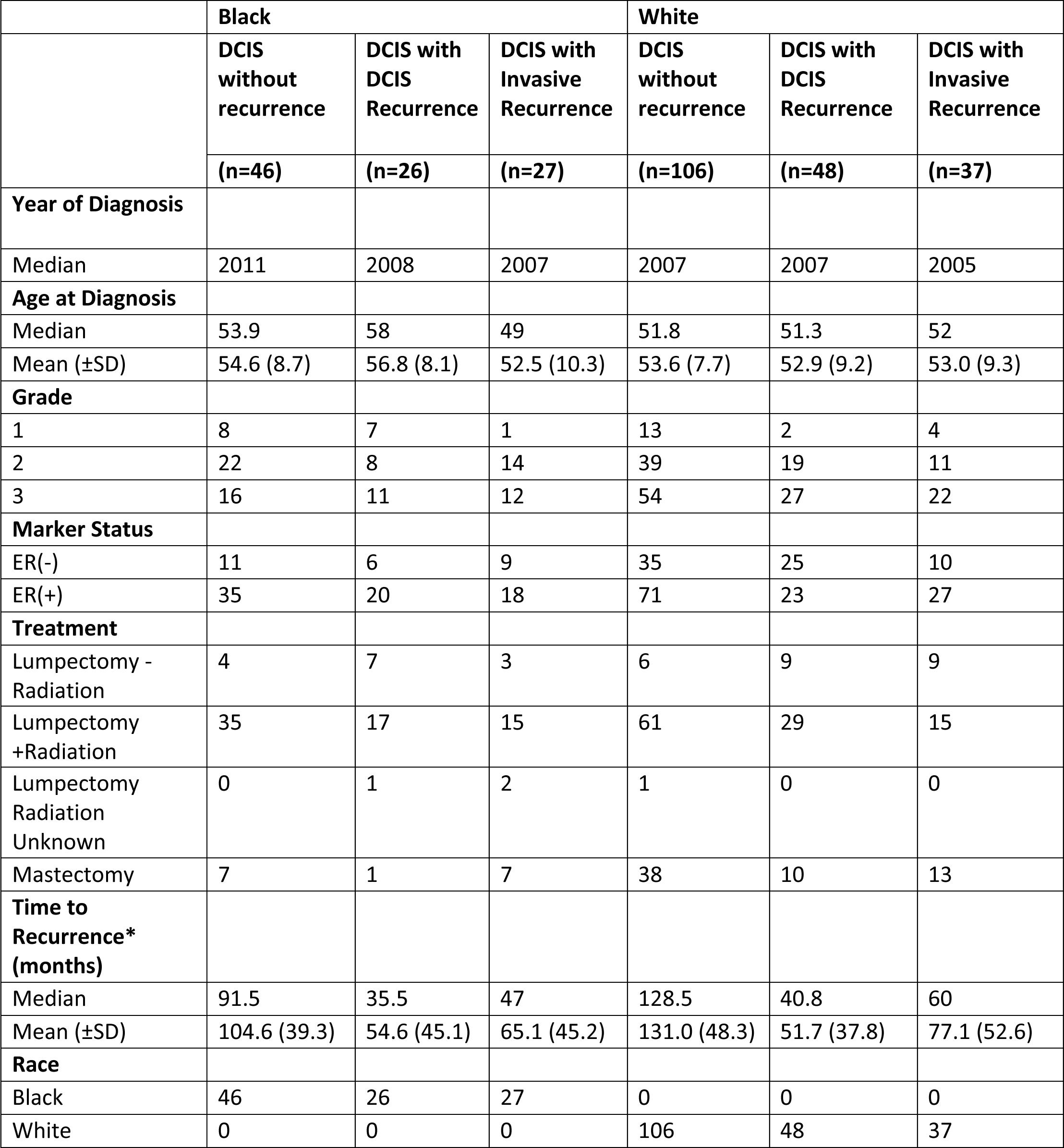
Combined Breast Pre-cancer Atlas Patient Cohorts stratified by SRR. * To end of follow-up for no recurrence.

To investigate outcome by SRR in the combined cohort, we first performed Kaplan-Meier analysis for time to any subsequent ipsilateral breast event (iBE), either DCIS or IBC recurrence. Here, Black women had significantly higher risk and shorter time to subsequent iBEs compared to White women (HR (95% CI): 1.5 (1.1 −2.1), P=0.023, **Figure 1A**). Moreover, Black women had significantly shorter time and almost twice the risk of IBC recurrence specifically compared to White women (HR (95% CI): 1.9 (1.2 – 3.2), P=0.011, **Figure 1B**). This is in concordance with previous reports^6,8,9,11,12^. Conversely, no significant difference was observed for time to DCIS recurrence by SRR (HR (95% CI): 1.3 (0.83 – 2.2), P=0.23, **Extended Data fig. 1B**).

Next, we investigated the impact of different treatment modalities on outcome. We found a higher proportion of patients with recurrence in the lumpectomy only group compared with the lumpectomy +RT or mastectomy groups (P=0.0007, **Extended Data fig. 1C**). To investigate if treatment differences could explain the observed outcome disparity by SRR, we performed multivariable regression analysis including SRR and treatment using full follow-up. While treatment type was highly significantly associated with outcome (Lump +RT HR (95% CI): 0.44 (0.28 – 0.68), P<0.001; Mastectomy HR (95% CI): 0.43 (0.26 – 0.73), P=0.002, **Extended Data fig. 1D**), Black women had significantly higher risk of iBE after adjusting for treatment (HR (95% CI): 1.44 (1.01 – 2.06), P=0.044, **Extended Data fig. 1D**). Moreover, Black women had 1.9 times higher risk of IBC recurrence compared to White women after adjusting for treatment (HR (95% CI): 1.92 (1.14 – 3.23), P=0.014, **Figure 1C**). Again, we observed no difference by SRR for DCIS recurrence only after adjusting for treatment (HR (95% CI): 1.14 (0.694 – 1.86), P=0.61, **Figure 1D**). We observed no significant difference in time to any iBE by treatment (**Extended Data fig. 1E-G**), probably due to small sample sets for each treatment type.

We recently presented the HTAN DCIS classifier consisting of 812 genes^18^, which was trained to predict iBEs within 5 years from treatment in the TBCRC cohort, and then validated in the RAHBT cohort. To see how the classifier performed by SRR, we performed multivariable analysis including the predicted risk groups from the HTAN DCIS classifier, clinical variables (Age, DCIS grade, RNA-based ER-status, and treatment) and SRR for 5-year follow-up (**Figure 1E**). We found that the HTAN DCIS classifier was highly predictive (Classifier high-risk group HR (95% CI): 10.06 (6.12 – 16.5), P<0.001), without substantial contribution by SRR (Black HR (95% CI): 1.12 (0.68 – 1.8), P=0.65). While the classifier performed well irrespective of SRR, we noted a trend towards enrichment of high-risk cases amongst Black women (P=0.065, **Extended Data fig. 1H**). To further investigate the classifier’s performance by SRR, we performed Kaplan-Meier analysis of the Classifier High-risk and Low-risk groups, respectively, stratified by SRR. While no significant difference by SRR was observed in the High-risk group (P=0.46, **Extended Data fig. 1I**, Black women classified as Low-risk had significantly shorter time to recurrence compared to White women (HR (95% CI) for Black women: 1.6 (1 – 2.5), P=0.032, **Figure 1F**). We noted that the biggest outcome difference was for late recurrences >10 years from treatment, whereas the classifier was trained to predict recurrence ≤5 years from treatment.

Taken together, these results indicate there are DCIS outcome differences by SRR, with Black women having significantly shorter time to subsequent iBE, specifically IBC recurrence. Moreover, the HTAN DCIS classifier was highly informative in predicting iBE within 5 years from treatment for both Black and White women.

### Gene expression differences by SRR

To investigate possible molecular differences underlying the observed DCIS outcome disparity by SRR, we performed differential gene expression analysis between primary DCIS from Black and White women. This analysis identified 384 differentially expressed (DE) genes (FDR<0.05, **Supplementary Table 1**). Next, we performed gene set enrichment analysis (GSEA) using the list of DE genes between DCIS from Black and White women with a variety of gene sets but found no significantly enriched pathways (**Extended Data fig. 2A** showing Hallmark pathways without significant enrichment). Since we identified DE genes between DCIS from Black and White women, we hypothesized there could be significant differences in gene expression between cases and controls by SRR. To investigate, we performed differential gene expression analysis in cases with iBE within 5 years from treatment versus the rest (as before^18^), but separately for Black and White women. The analysis of Black cases (n=33) vs controls (n=66) identified 266 DE genes (FDR<0.05, **Supplementary Table 2, Extended Data fig. 2B**), whereas analysis of White cases (n=51) vs controls (n=140) identified 812 DE genes (FDR<0.05, **Supplementary Table 3**, **Extended Data fig. 2C**).

**Figure 2:**
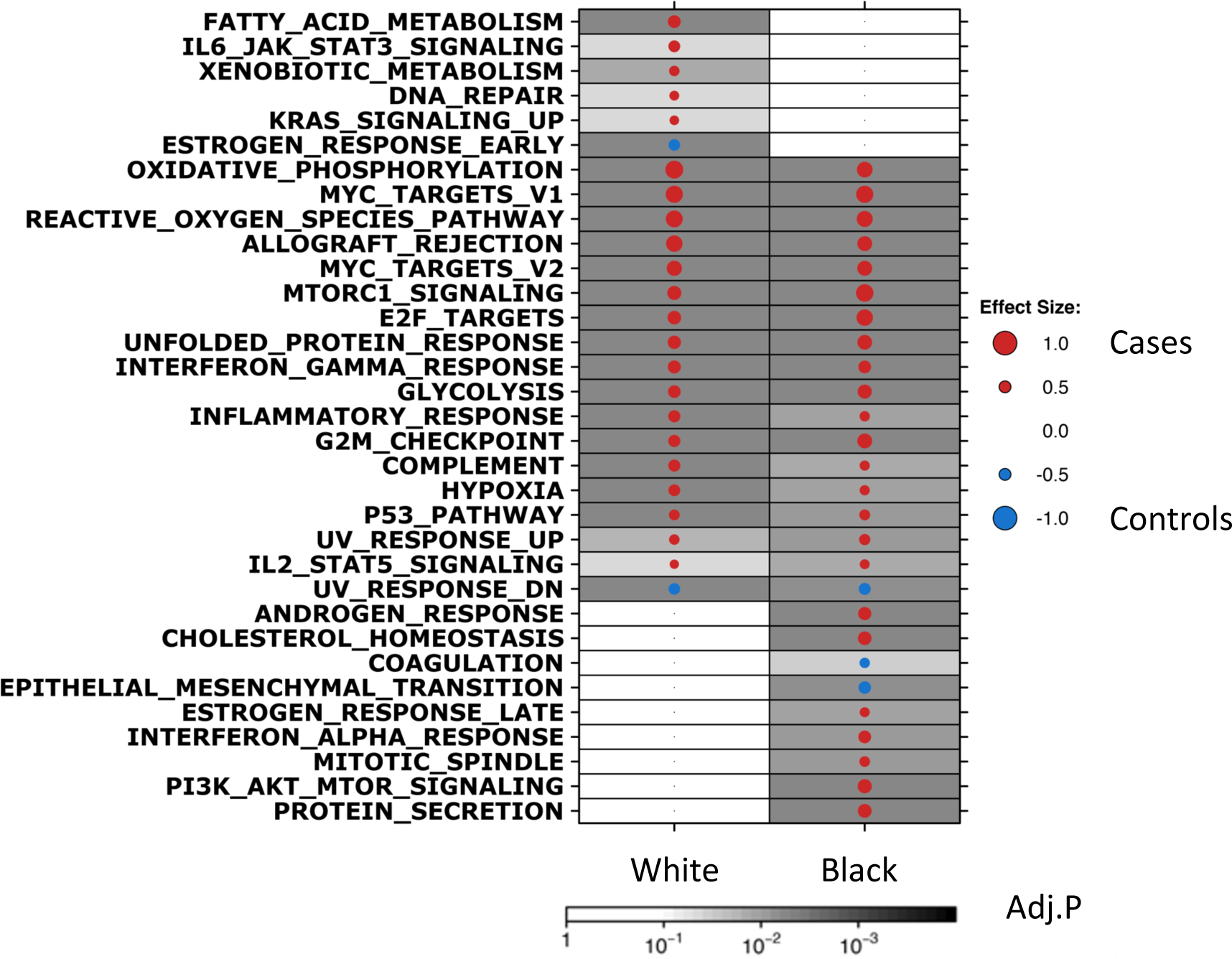
Gene Set Enrichment Analysis by SRR and outcome groups. GSEA Hallmark analysis of differentially expressed genes between DCIS from White cases vs controls (left column) and Black cases vs controls (right column), respectively. Dot size and color represent the magnitude and direction of pathway deregulation. Background shading indicates FDR. Effect size and FDR from GSEA algorithm.

We performed GSEA using the Hallmark gene sets, which identified many common pathways associated with iBEs in both Black and White women, including Allograft Rejection, cell-cycle associated pathways (E2F Targets, G2M Checkpoint), pathways involved in metabolism (Oxidative Phosphorylation, Glycolysis), and mTORc1 signaling (**Figure 2**). However, we also observed pathways that were differentially enriched by SRR, such as Fatty Acid Metabolism, IL6/JAK/STAT3 Signaling, and DNA Repair, which were associated with recurrence in White cases only. Conversely, Androgen Response and Interferon Alpha Signaling were amongst the pathways that were enriched in Black cases only. Finally, Estrogen Response pathways were significantly enriched in White controls and Black cases, further indicating fundamental pathways that may play different roles in iBEs depending on SRR.

To further investigate differential pathway enrichment in iBEs by SRR, we performed GSEA with GO terms, KEGG, and REACTOME gene sets (**Extended Data fig. 2D-F**). We noted that pathways involved in immune response, such as Allograft Rejection (Hallmark, KEGG), Antigen Processing and Presentation (KEGG), and Natural Killer Cell Mediated Cytotoxicity (KEGG) were up in cases regardless of SRR. However, some immune-associated pathways were specifically enriched in Black cases, such as Activation of Immune Response (GO), Immune Effector Processes (GO), Leukocyte Cell-Cell Adhesion (GO), and T-cell Receptor Signaling (KEGG). Conversely, Positive Regulation of T-cell Mediated Cytotoxicity (GO) and Regulatory T-cell Differentiation (GO) were up in White cases only, in addition to IL-6/JAK/STAT3 signaling (Hallmark), which is known to suppress the antitumor immune response^21^. These analyses further highlighted pathways that were differentially associated with 5-year outcome between DCIS from Black versus White women.

Together these results suggest different pathways are involved in iBEs by SRR. Of the 266 DE genes identified in Black case-vs-control analysis, 99 (37%) were represented in the previously identified HTAN DCIS classifier, whereas 357 (44%) of the 812 DE genes identified in White case-vs-control analysis were included in the classifier. Correlation analysis of the effect size (ES) of the 812 genes included in the HTAN DCIS classifier in the Black case-vs-control analysis and White case-vs-control analysis showed significant correlation (R=0.72, P<2.2e-16, **Extended Data fig. 2G**), however the ES from analysis in Black patients were attenuated compared to White patients. This analysis, combined with the worse outcome amongst Black women in the HTAN DCIS classifier Low-risk group (**Figure 1F**) suggests genes associated with iBEs in Black women may be underrepresented in the classifier.

### Established biomarkers, cell type distribution, and genomic aberrations by SRR

To further investigate molecular differences in DCIS from Black versus White women, we analyzed established IBC biomarkers by SRR and outcome, including gene expression of *ESR1* (ER) and *ERBB2* (HER2) and the PAM50 intrinsic subtypes. ESR1 expression was significantly elevated in Black compared to White women (P=0.034, **Figure 3A**), consistent with previous studies that found more ER+ DCIS tumors in Black women compared to other racial/ethnic groups^11,12^. Moreover, as indicated by GSEA (**Figure 2**), we found a significant difference in ESR1 expression by SRR and outcome groups, with White cases having significantly lower ESR1 expression compared to White controls (P=0.03, **Figure 3B**). No significant difference in ESR1 was observed between Black cases and Black controls. Furthermore, White cases had significantly lower ESR1 expression compared to all other groups combined (Black cases and all controls P=0.008, **Figure 3C**). For ERBB2 expression we observed the opposite trend, with significantly lower expression in Black women compared to White (P=0.021, **Figure 3D**). Again, a difference by outcome was observed for White women only, with increased ERBB2 expression in White cases versus controls, although it did not reach statistical significance (P=0.064, **Figure 3E**). Moreover, White cases had significantly higher ERBB2 expression compared to all other groups combined (P=0.014, **Figure 3F**). These results are in contrast to results obtained by analysis of ER and HER2 status in IBCs from TCGA, which reported significantly more ER+ tumors in women of European compared to women of African ancestry, with no difference in HER2 expression by ancestry^22^.

**Figure 3:**
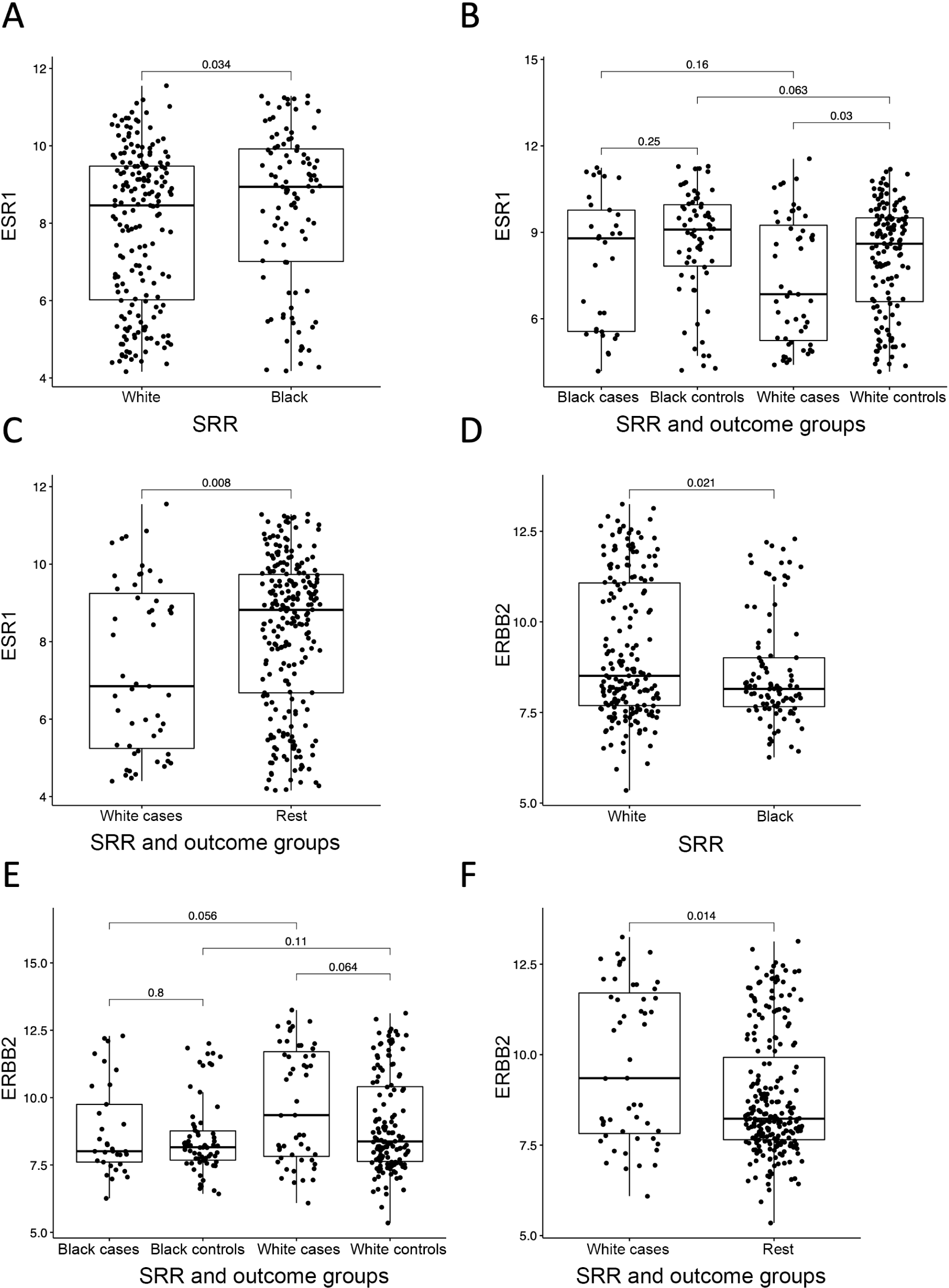
ER and HER2 expression by SRR and outcome groups. A) ER (*ESR1*) gene expression by SRR. B) ER (*ESR1*) gene expression by SRR and outcome groups. C) ER (*ESR1*) gene expression in White cases vs Black cases and all controls combined. D) HER2 (*ERBB2*) gene expression by SRR. E) HER2 (*ERBB2*) gene expression by SRR and outcome groups. F) HER2 (*ERBB2*) gene expression in White cases vs Black cases and all controls combined. A-F) Boxplots represent median, 0.25 and 0.75 quantiles with whiskers at 1.5x interquartile range. P-values from Wilcoxon rank-sum test.

Next, we analyzed the association between PAM50 subtypes and SRR. We found no significant association between PAM50 and SRR only (P=0.085, **Extended Data fig. 3A**), but identified a significant association for Black controls who were enriched for the Normal-like subtype (P=0.016, **Extended Data fig. 3B**) in the analysis including SRR and outcome groups. We note that we observed no enrichment for Basal-like DCIS in Black women, despite these being reported as more common in IBC from Black/African women^11,22–26^. This could be a reflection of the PAM50 Basal-like subtype not being fully applicable to early-stage breast tumors like DCIS^18,27^.

GSEA indicated an increased immune component in cases versus controls for both Black and White women (**Figure 2 and Extended Data fig. 2D-F**). To investigate differences in (immune) cell type composition, we leveraged available CibersortX data from the TBCRC and RAHBT cohorts^18^. We observed no significant difference in cell type distribution based on SRR (**Extended Data fig. 4**). However, cell type distribution by SRR and outcome groups revealed significant differences (**Figure 4A, Extended Data fig. 5A-O**), with myeloid dendritic cells (mDCs) and CD4 T-cells significantly higher in both Black and White cases versus controls (ES_mDC, Black_=1.4, P_mDC, Black_=0.0056; ES_mDC, White_=1.18, P_mDC, White_=0.0052; ES_CD4, Black_=2.29, P_CD4, Black_=0.0013; ES_CD4, White=_1.63, P_CD4, White_=0.01). Moreover, macrophages, monocytes, plasmacytoid dendritic cells (pDCs), NKT-cells, and overall immune cell population were significantly higher in White cases versus controls (ES_macro_=1.39, P_macro_=0.01; ES_mono_=2.39, P_mono_=0.0084; ES_pDC_=1.46, P_pDC_=0.0035; ES_NKT_=1.27, P_NKT_=0.0014, ES_immune_=1.24, P_immune_=0.023). For most of these, increased immune cell populations could be observed in Black cases vs. controls, although they did not reach statistical significance. However, fibroblasts and endothelial cells showed significantly lower proportions in Black cases versus Black controls (ES_fibro_=0.83, P_fibro_=0.0032; ES_endo_=0.91, P_endo_=0.024), with samples from Black cases depleted of fibroblasts compared to all other SRR-outcome groups. Moreover, high levels of fibroblasts were associated with better 5-year outcome in Black (HR (95% CI): 0.004 (0.0001 – 0.16), P=0.003), but not White women (HR (95% CI): 0.57 (0.032 – 10), P=0.70).

**Figure 4:**
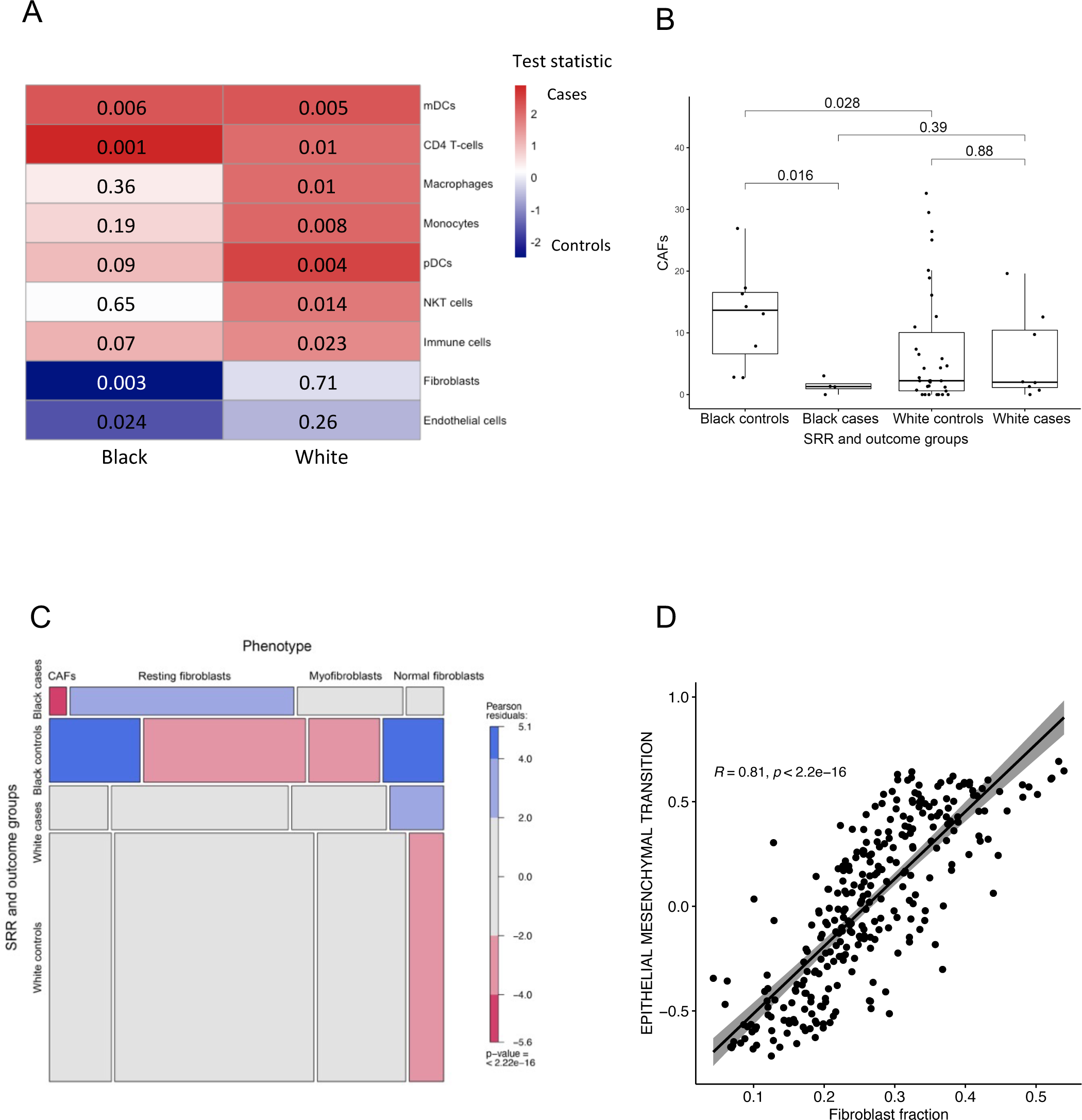
Cell type distribution by SRR and outcome groups. A) Heatmap of P-values (Wilcoxon rank-sum test) for inferred cell types in Black cases vs controls (left column) and White cases vs controls (right column). mDC = Myeloid dendritic cells. pDC = Plasmacytoid dendritic cells. B) Cancer associated fibroblasts (CAFs) distribution by SRR and outcome groups in MIBI sample-level data (n=54). Boxplot represents median, 0.25 and 0.75 quantiles with whiskers at 1.5x interquartile range. P-values from Wilcoxon rank-sum test. C) Mosaic plot showing distribution of fibroblast phenotypes by SRR and outcome groups in MIBI single-cell data (n=4926). P-value from Chi^2^ test. D) Correlation between EMT and fibroblast fraction by RNA-seq analysis in combined TBCRC and RAHBT cohorts, all samples regardless of SRR (n=290). Correlation coefficient and P-value from Pearson correlation analysis.

To further investigate the differences in fibroblast population in Black cases versus other SRR-outcome groups, we used Multiplex Ion Beam Imaging (MIBI) protein expression data from the RAHBT cohort^18,19^. Previous MIBI analysis identified four different fibroblast phenotypes, namely normal fibroblasts, myofibroblasts, resting fibroblasts, and cancer associated fibroblasts (CAFs)^19^. Using available sample-level MIBI data from 54 patients, we found that CAFs were significantly depleted in Black cases vs. controls (P=0.016, **Figure 4B**). None of the remaining fibroblast phenotypes or overall fibroblast population showed significant difference between SRR-outcome groups (**Extended Data fig. 5P-S**). However, using single-cell level MIBI data, we found that the overall distribution of fibroblasts differed between SRR-outcome groups, with Black cases depleted for CAFs but enriched for resting fibroblasts, and Black controls depleted for resting- and myofibroblasts while enriched for normal fibroblasts and CAFs. White cases were enriched, and White controls depleted, of normal fibroblasts (**Figure 4C**). Finally, we performed single sample gene set variation analysis of the 34 Hallmark pathways in **Figure 2A**. We correlated the enrichment score for each pathway in each sample to the respective sample’s fibroblast fraction. This analysis showed that the Epithelial Mesenchymal Transition (EMT) pathway and fibroblast fraction were highly correlated across all samples (R=0.81, P<2.2e-16, **Figure 4D**). According to Hallmark GSEA (**Figure 2A**), the EMT pathway was enriched in Black controls compared to cases, correlating with the observed enrichment of fibroblasts in these samples. These results indicate increased mesenchymal gene expression in DCIS samples from Black cases compared to Black controls. Taken together, our results further indicate that different cell abundances contribute to the observed outcome differences based on SRR.

Finally, we investigated differences by SRR on the genomic level by analyzing copy number variations (CNVs) by SRR and outcome groups (**Table 2, Extended Data fig. 6**). Amplification of 17q21.31 and 17q12, and deletion of 8p11.22, were significantly enriched in DCIS from White versus Black women (P_17q21.31_=0.0004, P_17q12_=0.025, P_8p11.22_=0.041, **Table 2A, Extended Data fig. 6A-C**). The amplification of 17q12 in DCIS from White women correlated with the observed increased ERBB2 expression in these lesions (**Figure 3D**). Analysis of CNVs by SRR and outcome groups found that deletion of 1p21.3 (P=0.033) and 3p14.1 (P=0.009) were enriched in White controls compared to White cases (**Table 2B, Extended Data fig. 6D, E**). We found no significant differential CNVs between Black cases and controls. None of the CNVs identified here were significantly enriched in IBC by ancestry in TCGA data^22^. Finally, we investigated Percentage of the Genome Altered (PGA), overall amplifications and deletions by SRR only and SRR-outcome, with no significant difference between groups (**Extended Data fig. 6F-K**). Taken together, our results indicate that there are larger differences in DCIS by SRR on the transcriptomic than the genomic level.

**Table 2:** Significant CNVs by SRR (A) and SRR and outcome groups (B).

| **CNVs** | **P=** |
| --- | --- |
| Amp 17q21.31 | **0.0004** |
| Amp 17q12 | **0.0245** |
| Del 8p11.22 | **0.0412** |

|  | **White cases vs controls** | **Black cases vs controls** |
| --- | --- | --- |
| **CNVs** | **P=** | **P=** |
| Del 1p21.3 | **0.033** | 0.614 |
| Del 3p14.1 | **0.009** | 0.913 |

## Discussion

We recently presented the HTAN DCIS Atlas which included molecular analyses of DCIS epithelium and microenvironment on the genomic, transcriptomic and proteomic level for 774 DCIS samples from 542 patients. In that study we identified epithelial and stromal subtypes specific to DCIS, and generated the HTAN DCIS classifier, but our analyses did not take SRR or ancestry into account. Recently there has been an increased focus on the need to investigate molecular differences between tumors based on ancestry, ethnicity, or race and the potential implication for screening and treatment options^28^. While race is a social construct, it is often associated with poor outcomes in part due to access to health care, social determinants of health, and racial inequity, and thus continues to offer incremental, useful information, including elucidation of health disparities^28–30^. Emerging evidence suggests there are biologic variations by race, supporting the drive to understand the pathways that could account for such differences. In the TAILORx trial of over 10,000 women with hormone receptor-positive, node-negative breast cancer, locoregional recurrence was significantly higher in Black women after adjustment for treatment, patient and tumor characteristics (HR=1.78 (1.15 – 2.77)) compared to White women^31^. Moreover, study cohorts have traditionally included few Black and Asian patients, or indeed have not collected information on ancestry, ethnicity, or race^32^. Thus, there is an unmet need to address disparities in IBC diagnosis and outcomes among Black women and other minorities, which can be in part addressed by better understanding key differences in tumor biology between White and Black women.

Here, we set out to investigate the molecular differences in DCIS based on SRR and ancestry by combining data from the two large DCIS cohorts from the HTAN DCIS Atlas. We generated global genetic ancestry calls from WGS data and compared these to SRR, which showed exceptionally high concordance between the two parameters, as expected. Thus, using SRR in the analyses we observed a significant outcome difference by race, with Black women having significantly shorter time to recurrence, and IBC recurrence specifically, compared to White women (**Figure 1A, B**). While treatment modality was strongly associated with outcome, it could not fully explain the observed outcome disparity between Black and White women (**Figure 1C, D, Extended Data fig. 1D**).

While we observed few clear genomic differences between DCIS tumors from Black and White women, we observed significant differences relating to gene expression and pathway enrichment between cases and controls within each SRR group (**Figure 2**). Several pathways were involved in recurrence in Black but not White DCIS, including Androgen Response, Interferon Alpha Response, and PI3K-AKT-MTOR Signaling. Conversely, DNA Repair, Fatty Acid- and Xenobiotic Metabolism, and KRAS Signaling were associated with recurrence in White women. Notably, Estrogen Response Early and -Late was enriched in White controls compared to cases, whereas Estrogen Response Late was enriched in Black cases versus controls (**Figure 2**). In line with this, we found that the ER- and HER2+ phenotypes were associated with iBEs in White women only, contrary to what has been shown in IBC. The finding that ER expression was lower in DCIS from White compared to Black women (**Figure 3A**) contrasts with results from analysis of IBC tumors from the TCGA^22^, but agrees with analysis of SEER data where Black women had significantly more ER+ DCIS compared to other racial/ethnic groups^11,12^. We observed a significant enrichment of the 17q12 amplification in DCIS from White women, which correlated with the observed increased ERBB2 RNA expression in these samples (**Figure 3D**). The 17q12 amplification was not differentially enriched between White cases and controls, suggesting that the observed trend of increased ERBB2 expression in White cases versus controls (**Figure 3E**) was caused by mechanisms other than amplification of the 17q12 locus, in line with previous reports on IBC^33^.

We found no difference in the distribution of PAM50 subtypes by SRR only but found that Black controls were enriched for the Normal-like subtype. This is somewhat surprising, since Basal-like/triple negative (TNBC) IBC is more prevalent in Black women compared to White^11,22–26^. Moreover, a study including more than 160.000 women reported that the risk of TNBC in Black women was almost twice that of White women after DCIS^11^. We and others have previously reported that the Basal-like subtype does not seem to apply fully to tumors at the DCIS stage^18,27^, which may in part explain this discrepancy.

Because DCIS is confined to the intraepithelial compartment, the DCIS microenvironment is very different from that of IBC. In our cohort, inferred immune cells showed an overall trend of enrichment in cases versus controls regardless of SRR. In line with this, pathway analyses showed that several immune-related pathways were enriched in cases compared controls for both Black and White patients (**Figure 2A, Extended Data fig. 2D-F**). Intriguingly, we found that low inferred fibroblast abundance was associated with recurrence within 5 years for Black women only (**Figure 4A, Extended Data fig. 4H**). This contrasts with IBC studies, where fibroblasts, and CAFs in particular, are associated with poor prognosis^34–38^. Recent studies suggested fibroblast abundance in DCIS contribute to invasive progression^39,40^. Importantly, none of these studies included race as a factor in the analyses. Intriguingly, by MIBI we here observed significantly reduced CAF levels in Black cases versus Black controls (**Figure 4B**), albeit in a very small data set. Moreover, using RNA-seq data we found that fibroblast abundance was highly correlated with EMT enrichment (**Figure 4D**).

According to American Cancer Society’s Breast Cancer Statistics 2022, Black women have slightly lower incidence of IBC than White women, but 40% higher IBC mortality rates^41^. This reflects in part that Black women have higher early incidence of TNBC and the poorest 5-year survival across race/ethnic groups within TNBC^41^. Several studies have sought for tumor biological features that can explain this discrepancy. One study found that IBC tumors from Black women had greater genetic heterogeneity and more basal gene expression, suggesting more aggressive tumor biology^26^. Another reported higher microvessel density and macrophage infiltration in IBC from Black women compared to White^42^. Huo et al. analyzed genomic, transcriptomic, proteomic and methylome data from IBCs in TCGA by ancestry. While they found many molecular differences between IBCs from women with African versus European ancestry, including CNVs, gene expression, and DNA methylation, most of the molecular differences were eliminated after adjusting for intrinsic subtype^22^.

While there is compelling epidemiologic and clinical evidence about differences in the biology and behavior of IBC by race or ancestry, it is not clear to what extent the difference in outcome is due to biologic or socioeconomic factors related to racial inequity. Martini and colleagues investigated differential gene expression in women with TNBC based on both SRR and ancestry^43^. They found that genes uniquely associated with SRR were involved in pathways associated with lifestyle diseases, and in clustering analysis separated African American women from Ghanaians and Ethiopians. The authors hypothesized that these genes represent distinct environmental influences unique to African American patients, supporting the premise of social determinants linked to racial constructs.

The HTAN DCIS classifier was trained to predict iBE within 5 years from treatment. While we here showed that the classifier performs well in predicting 5-year iBEs regardless of SRR, we found significantly poorer outcome in the HTAN DCIS classifier low-risk group for Black compared to White women when including the full follow-up time (**Figure 1F**). The observed difference was greatest >10 years after initial treatment, which the classifier was not trained to detect. Liu et al found that Black women had significantly higher risk of developing ipsilateral IBC away from the original DCIS lesion, as well as contralateral IBC, indicating there is an underlying genetic susceptibility to IBC, early exposures, and/or interactions between these fundamental to breast tumors in Black women^11^. While our recurrence dataset included only ipsilateral events, one can speculate that the patients with recurrence >10 years from treatment could be de-novo lesions not related to the original DCIS, as shown in clonality studies^44^.

Our study has several limitations. The combined cohort analyzed here consisted of 34% Black women, which is a large fraction compared to previous studies analyzing DCIS by race or ethnicity^6–12^. Nevertheless, our results are based on a relatively small sample set, and we were not well powered for several analyses that stratified the cohort by SRR. In addition, the original cohorts included only 5 women identifying as Asian and a single Pacific islander, thus these groups were excluded from our analyses. Future studies investigating molecular differences in DCIS by SRR and ancestry should focus on including sizeable representations of other racial groups. Our cohort also did not include information on ethnicity, socioeconomic, or environmental factors, thus we were unable to evaluate their contribution to outcome disparity. Furthermore, race is often used as a surrogate for global genetic ancestry. Importantly, we noticed a broad range in genetic ancestry in both self-reported White and Black patients, which provides an important future opportunity to evaluate the role of local and global genetic ancestry in determining breast cancer biology.

Taken together, our results indicate there are fundamental biologic differences related to iBEs in Black and White women. Our study highlights the need for larger molecular and epidemiological studies to identify biological factors that contribute to the racial differences in DCIS and IBC outcome, as well as opportunities to tailor prevention strategies according to relevant and variable pathways. Given the distinct nature of the disease and risk for recurrence based on SRR, evaluating risk predictors either based on established clinical elements such as receptor status or complex molecular parameters (such as the HTAN classifier), should take race into account and be aware of the composition of the population to be evaluated.

## Acknowledgements

This publication is part of the HTAN (Human Tumor Atlas Network) Consortium paper package. A list of HTAN members is available at humantumoratlas.org/htan-authors/. This work was supported by the following grants: U2C CA-17-035 Pre-Cancer Atlas (PCA) Research Centers grant, R01CA193694 (RBW, GAC), BCRF PPI-18-006 (RBW), DOD BC132057 (ESH), and 1R01-CA185138-01 (ESH). We are grateful for the funding support to the TBCRC from The Breast Cancer Research Foundation and Susan G. Komen.

## Author Contributions

Conceptualization: ESH, RBW, GAC, JRM. Investigation: SHS, KEH, and RBW. Resources: BH, FC, KG, MK, SW, AD, TK, PM. Writing – Original Draft: SHS and RBW. Writing – Review & Editing: All co-authors. Funding Acquisition: ESH, RBW, GAC. Supervision: CC, GAC, ESH, and RBW.

## Additional information

Supplementary Information is available for this paper: Supplementary Tables 1–3. Correspondence and requests for materials should be addressed to Dr. Robert B. West.

## Data and code availability

We analyzed data that is publicly available on the HTAN data portal (https://www.ncbi.nlm.nih.gov/projects/gap/cgi-bin/study.cgi?study_id=phs002371.v3.p1).

No original code was generated for this analysis. All scripts needed to reproduce the analysis will be made available online by the time of publication.

## Methods

### Patient samples

We used data from two previously described case-control cohorts of patients diagnosed with pure DCIS with or without a subsequent ipsilateral breast event (iBE, either DCIS or IBC) after surgical treatment^1^. The cohorts were combined to improve statistical power to allow comparison by SRR. Identical eligibility criteria were used for outcome analysis in both cohorts. Briefly, The Resource of Archival Breast Tissue (RAHBT) cohort includes women ≥18 years of age with documented cases of premalignant breast disease including DCIS. The study was approved by the Washington University in St. Louis Institutional Review Board (IRB ID #: 201707090). TBCRC 038 is a retrospective cohort with patients from 12 participating TBCRC (Translational Breast Cancer Research Consortium) sites, which includes women treated for DCIS at one of the enrolling institutions between 01/01/1998 and 02/29/2016. The study was approved by the Duke Health Institutional Review Board (Protocol ID: Pro00068646) as well as the IRB at each participating institution.

For the combined cohort studied here, samples from a total of 313 patients (n_TBCRC_ = 216, n_RAHBT_ = 97) were previously analyzed by RNA-seq. We excluded patients with SRR listed as Asian (n=5) and Pacific Islander (n = 1) due to low numbers in these groups, in addition to 17 patients with missing SRR. The final cohort consisted of 290 patients (**Table 1**) and was composed of 34.1% Black women and 65.9% White women. For White women, the median age at diagnosis was 51.6 years, and median year of diagnosis was 2007. Median time to recurrence with ipsilateral IBC was 60 months, and time to diagnosis of ipsilateral DCIS was 40.8 months. For White women with no iBEs, median follow-up extended to 128.5 months. Treatment of initial DCIS ranged from lumpectomy with radiation (55.0%), and no radiation (12.6%) to mastectomy (31.9%). For Black women, the median age at diagnosis was 54 years, and median year of diagnosis was 2009. Median time to recurrence with ipsilateral IBC was 47 months, and time to diagnosis of ipsilateral DCIS was 35.5 months. For Black women with no iBEs, median follow-up was 91.5 months. Treatment of initial DCIS ranged from lumpectomy with radiation (67.7%), and no radiation (14.1%) to mastectomy (15.2%). Tumor low-pass WGS data was available for 208 patients (71.7%), 33.7% of whom were Black.

### Sequencing and Multiplex Ion Beam Imaging (MIBI) data

We analyzed data that is publicly available on the HTAN portal (https://www.ncbi.nlm.nih.gov/projects/gap/cgi-bin/study.cgi?study_id=phs002371.v3.p1) including RNA and DNA sequencing, metadata, and MIBI data. We used copy number variation, Cibersort X, and PAM50 subtype calls from the HTAN DCIS atlas. For details see^1^.

### Inference of global genetic ancestry

Given the low DNA sequence coverage of the WGS data, we leveraged QUILT (v0.1.9)^2^ to genotype the TBCRC and RAHBT cohorts. QUILT is designed for genotyping low sequence coverage samples using a Gibbs sampling method to facilitate rapid imputation. Genotyping was conducted in batches considering 5 Mbp windows and 100 samples at a time. We used the 1000 Genomes Project as a reference panel (n=3,202)^3^. We used default parameters apart from “– buffer 10000 –nGen 100”. The resulting variants were annotated with dbSNP (b155) and converted to EIGENSTRAT format using a combination of plink^4^ (v1.9) and CONVERTF (v3.0). Finally, ancestry weights were inferred using SNPWEIGHTS^5^ (v2.1) and the snpwt.NA reference panel.

### Differential abundance analyses

Differential abundance analysis was performed using the R package DESeq2 v1.30.1^6^ with default options. P-values were adjusted for multiple testing using the Benjamini-Hochberg method. False-discovery rate (FDR) <0.05 was considered significant for all DESeq2 analyses.

### Gene Set Enrichment Analyses

Gene set enrichment analyses were performed using fgsea R package (v1.16.0) based on the MsigDB Hallmark, Kegg, Reactome, and Gene Ontology gene sets v2023.1^7^. All genes from differential abundance analyses were included and were ranked by their signed adjusted P-values. Pathways with false-discovery rate (FDR) <0.05 were considered significantly enriched.

Single sample gene set variation analysis was performed using the GSVA R package^8^ (v1.38.2) using default parameters.

### Outcome analysis

Associations with time to event were quantified using Cox Proportional Hazard model. Kaplan-Meier plots as implemented in the R packages survival (v3.3-1) and survminer (v0.4.9) were used to visualize outcome differences.

We previously generated the HTAN DCIS prognostic classifier^1^ which was trained to predict iBEs within 5-years from treatment, regardless of treatment type. Time to recurrence analyses in the present paper were made either using 5-year follow-up, or full follow-up, as specified in the text.

### Statistical analyses

Wilcoxon rank-sum test was used to compare continuous distributions between two groups. Reported effect size was calculated as median(case)/median(control). Gene expression data were quantified as VST normalized reads generated using the DESeq2 R package (v1.30.1). All statistical analyses were implemented in the R statistical environment (v4.0.5). P-values were corrected for multiple hypothesis testing using the Benjamini-Hochberg method.

Association between categorical variables were analyzed and visualized using mosaic plots from the vcd (1.4-11) package in R with default Pearson’s chi-squared residuals. Boxplots, heatmaps, scatterplots and barplots were generated using the BoutrosLab.plotting.general R package v7.0.3^9^, or the R packages ggplot2 (v3.3.6, boxplots), or corrplot (v0.92, scatterplots).

---

**Extended Data Figure 1:**
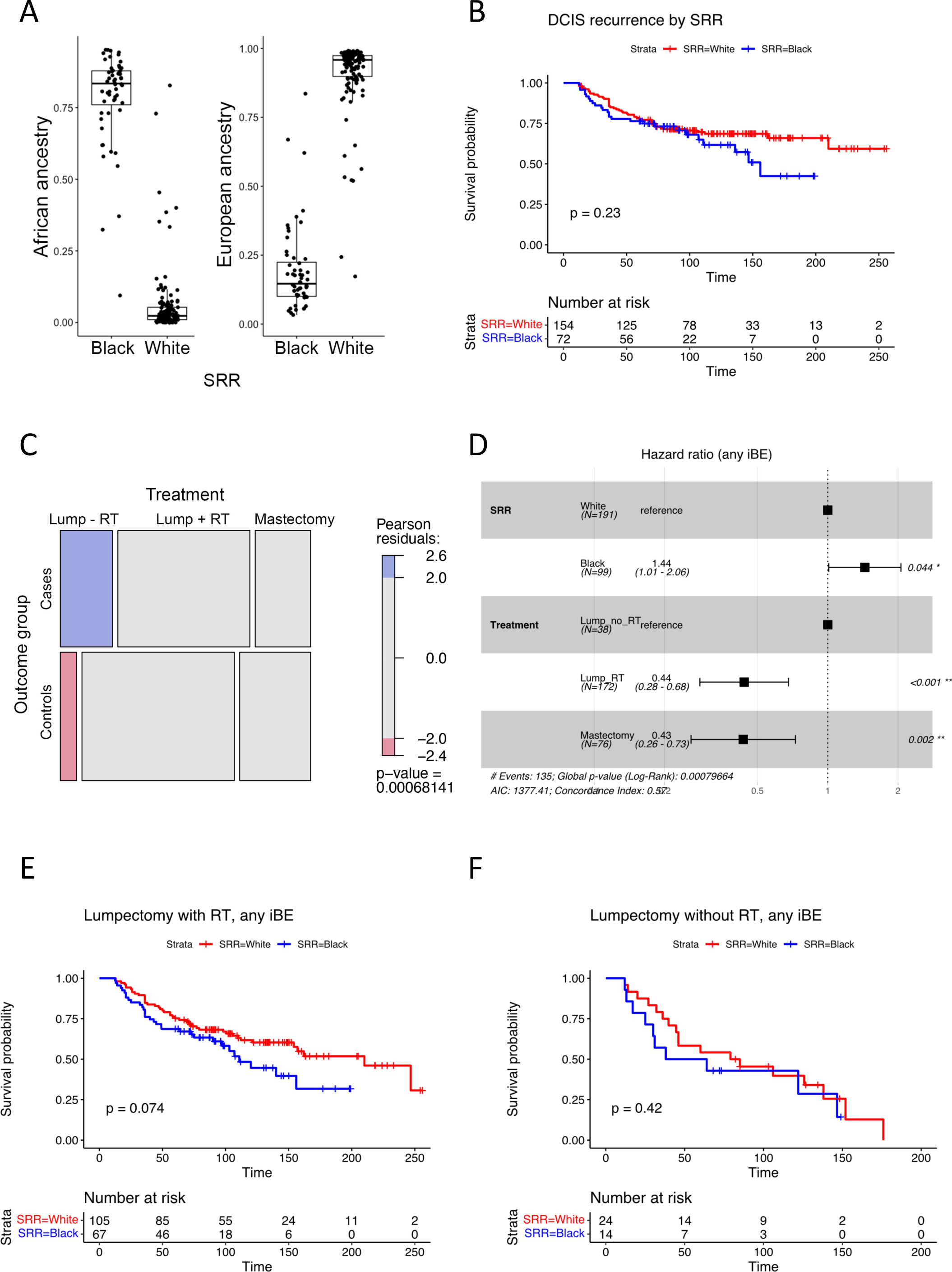
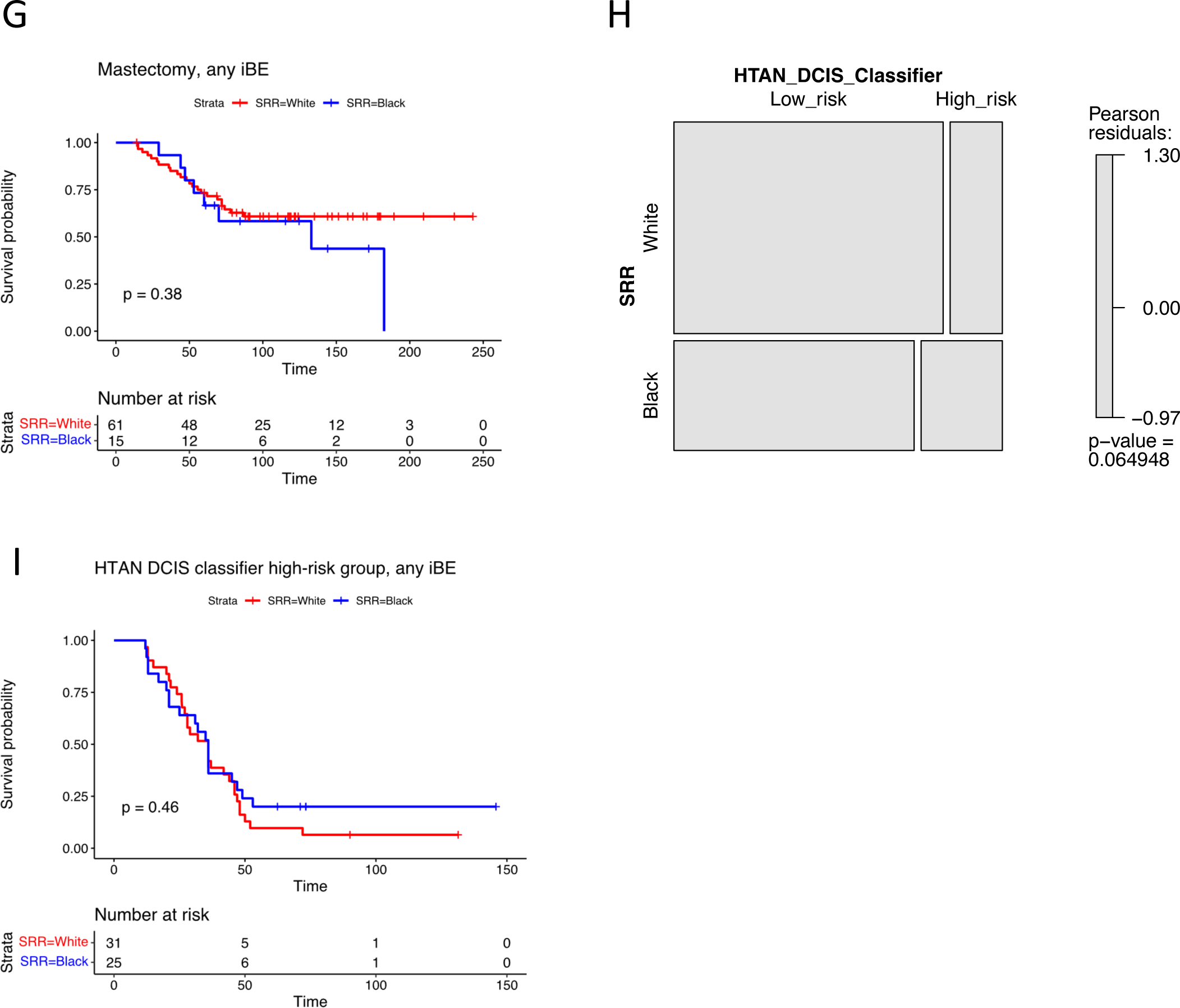
Outcome analysis by SRR. A) Boxplot of SRR vs African ancestry (left) and European ancestry (right), respectively. Boxplots represent median, 0.25 and 0.75 quantiles with whiskers at 1.5x interquartile range. B) Kaplan-Meier plot of time to DCIS recurrence only (full follow-up) stratified by SRR. C) Mosaic plot showing distribution of treatment type by outcome groups. P-value from Chi^2^ test. D) Forest plot of multivariable Cox regression analysis including SRR and treatment type, for any iBE (full follow-up). E) Kaplan-Meier plot of time to any iBE (full follow-up) for patients treated by lumpectomy and radiation treatment stratified by SRR. F) Kaplan-Meier plot of time to any iBE (full follow-up) for patients treated by lumpectomy without radiation treatment stratified by SRR. G) Kaplan-Meier plot of time to any iBE (full follow-up) for patients treated by mastectomy stratified by SRR. H) Mosaic plot showing distribution of HTAN DCIS classifier risk by SRR. P-value from Chi^2^ test. I) Kaplan-Meier plot of time to iBE (full follow-up) in the HTAN DCIS classifier high-risk group stratified by SRR. B, E, F, G, I) P-values from log-rank tests.

**Extended Data Figure 2:**
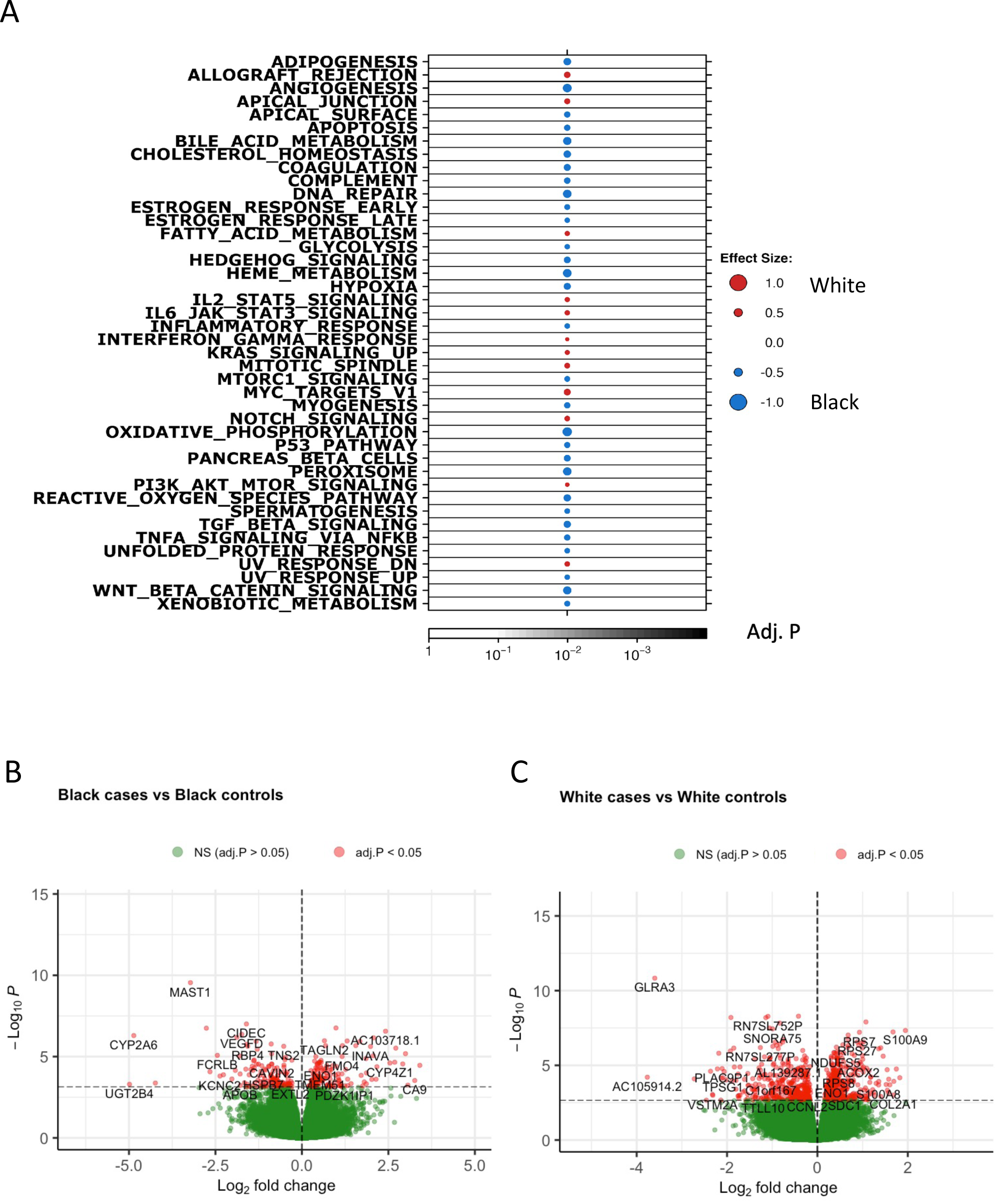
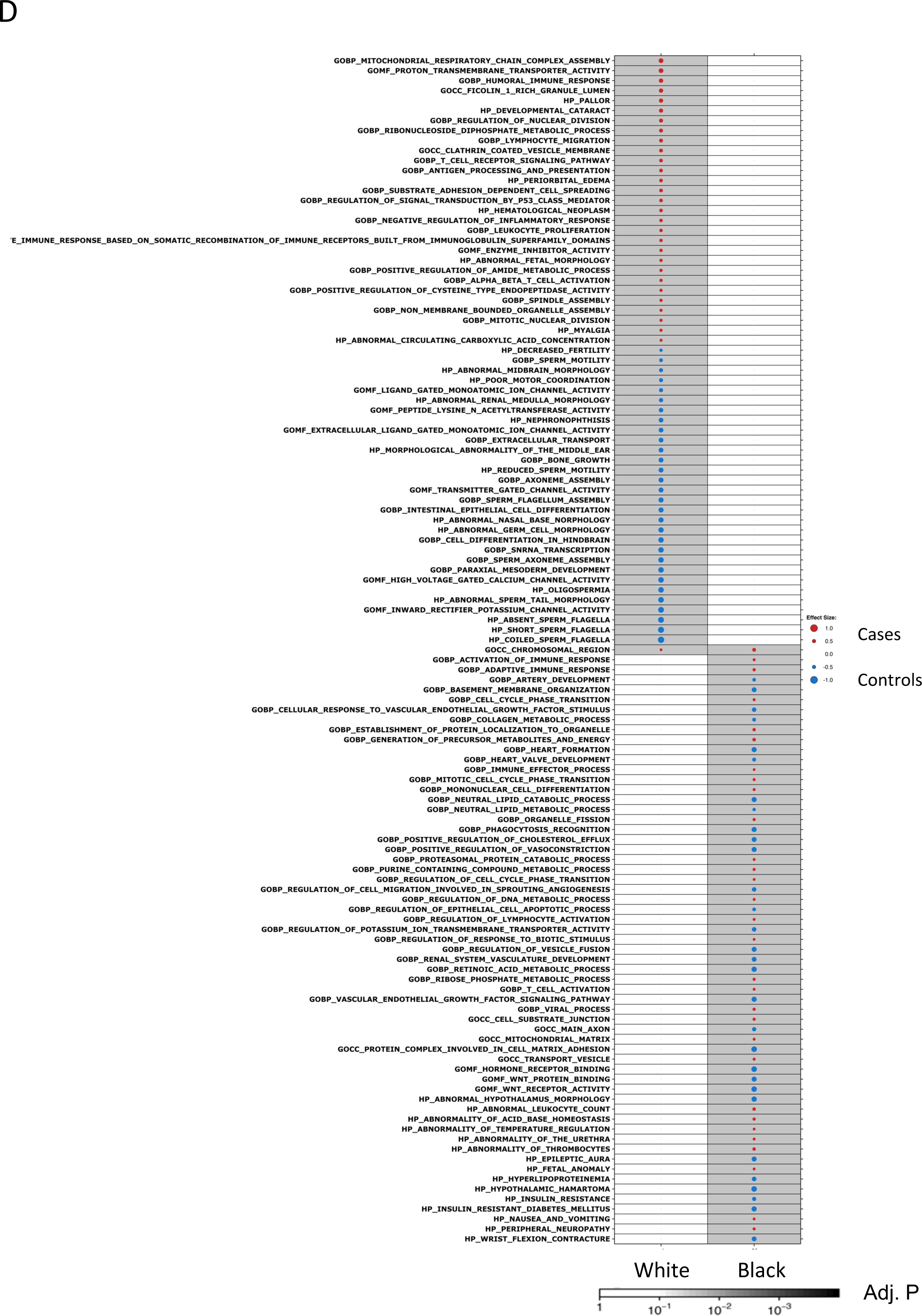
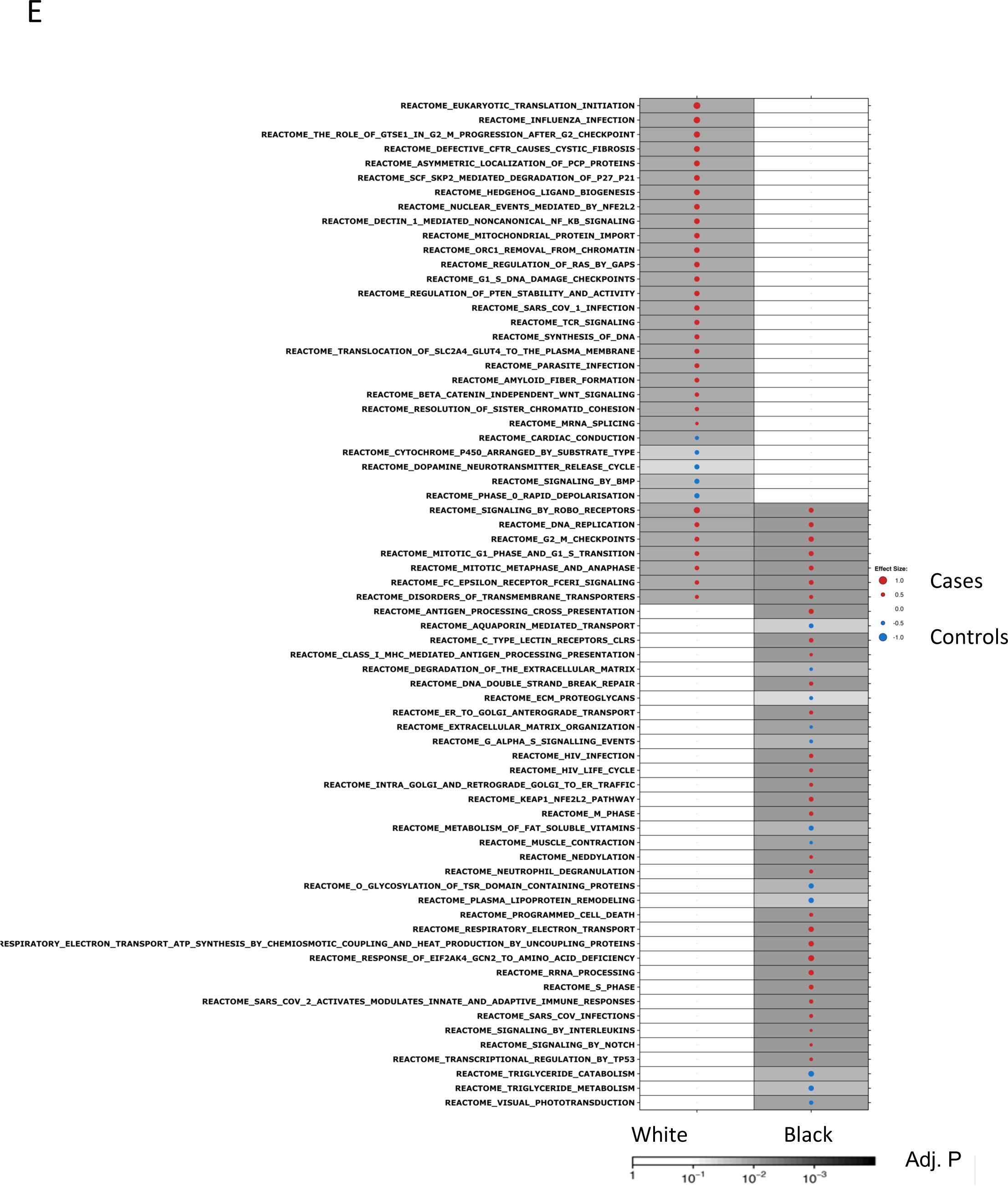
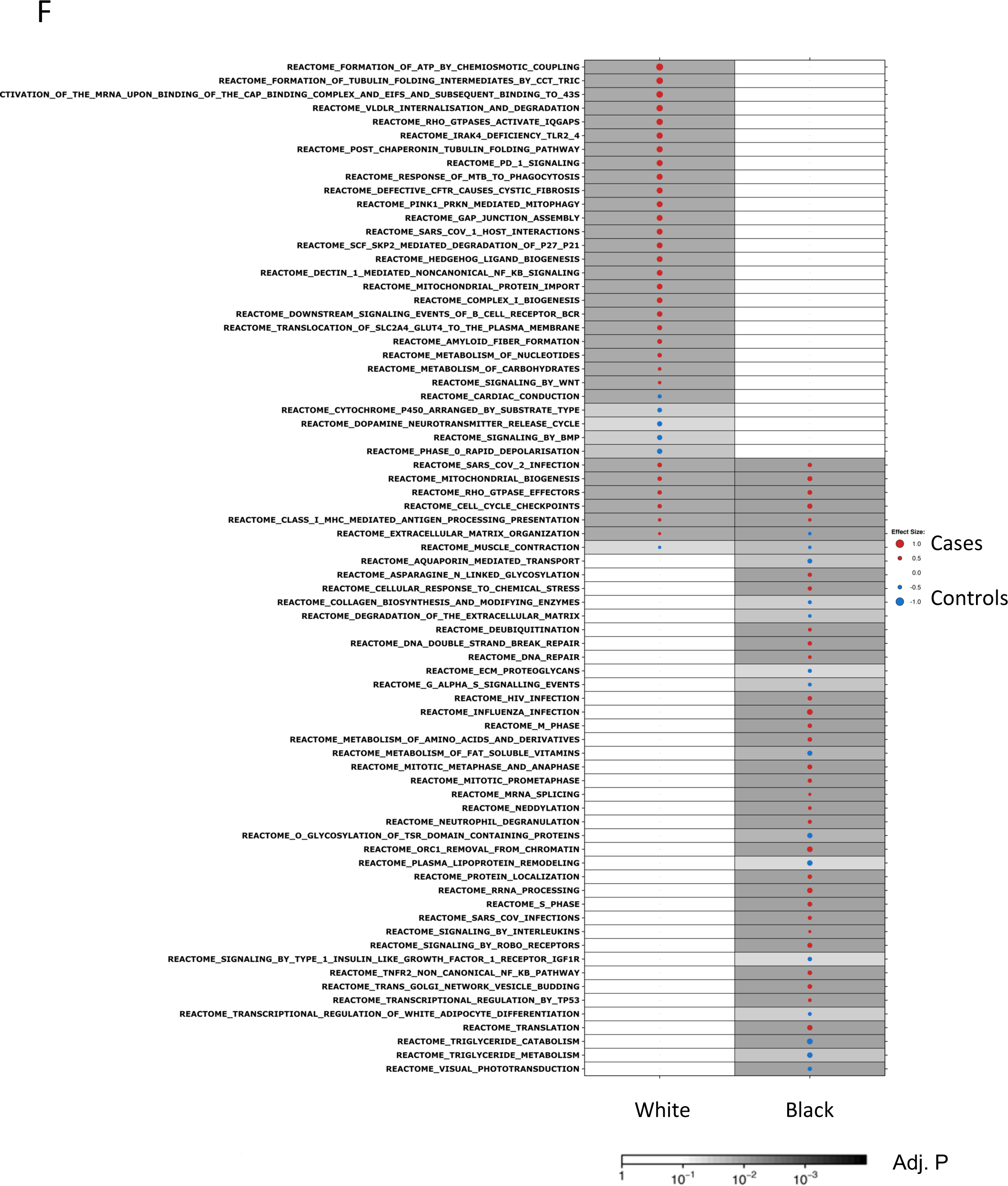
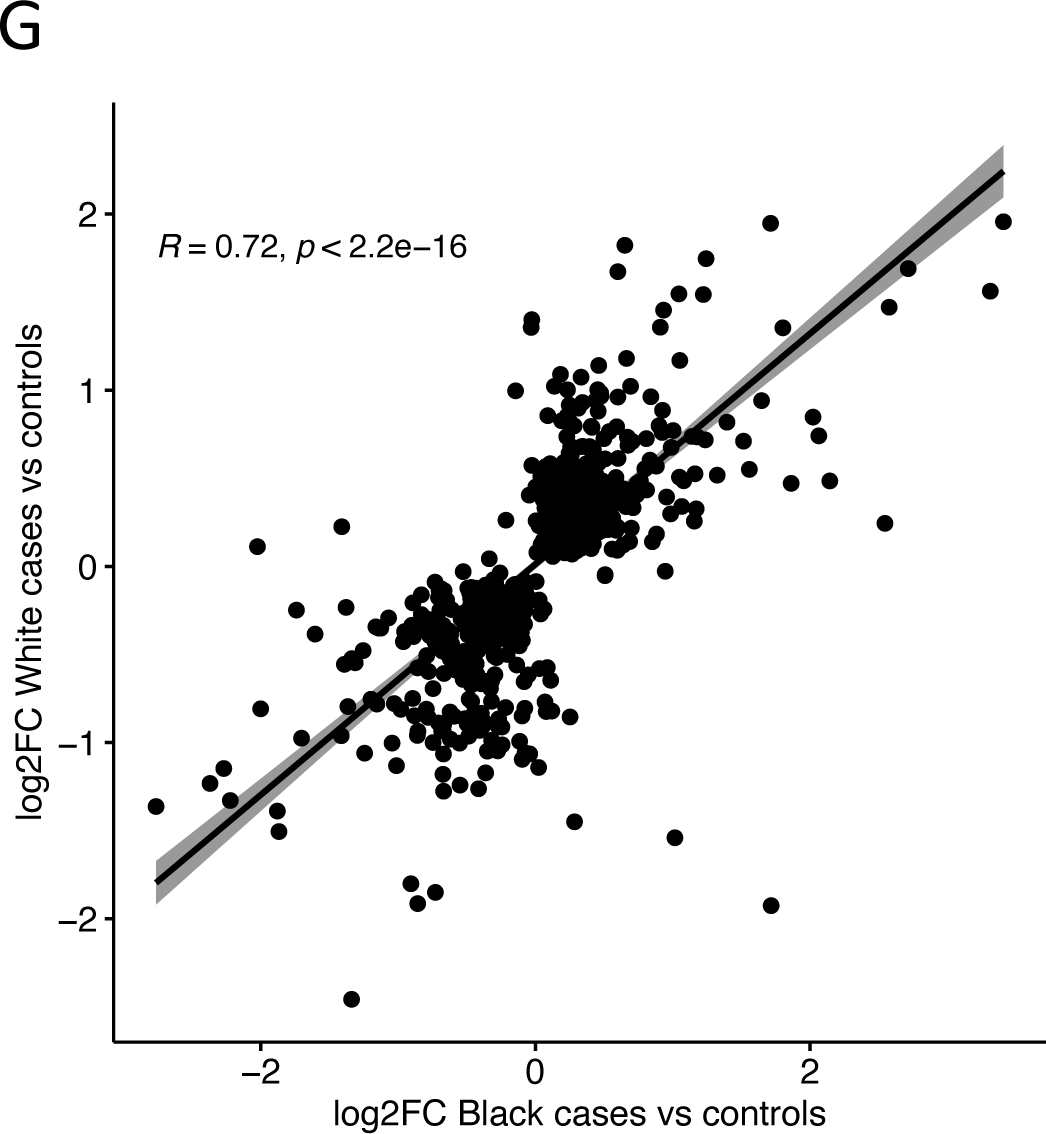
Differential gene expression by SRR and outcome groups. A) GSEA Hallmark analysis of differentially expressed genes between DCIS from White and Black women. B) Volcano plot of differentially expressed genes from Black case-vs-controls analysis. Genes with adj. P<0.05 in red (n=266). C) Volcano plot of differentially expressed genes from White case-vs-controls analysis. Genes with adj. P<0.05 in red (n=812). D – F) GSEA of differentially expressed genes between DCIS from White cases vs controls (left column) and Black cases vs controls (right column), respectively, using GO terms (D), KEGG (E), and Reactome (F) gene sets. A, D, E, F) Dot size and color represent the magnitude and direction of pathway deregulation. Background shading indicates FDR. Effect size and FDR from GSEA algorithm. G) Scatter plot of log2FC values of the 812 genes included in the HTAN DCIS classifier from Black case-vs-control analysis (x-axis) versus White case-vs-control analysis (y-axis). Correlation coefficient and P-value from Pearson correlation test.

**Extended Data Figure 3:**
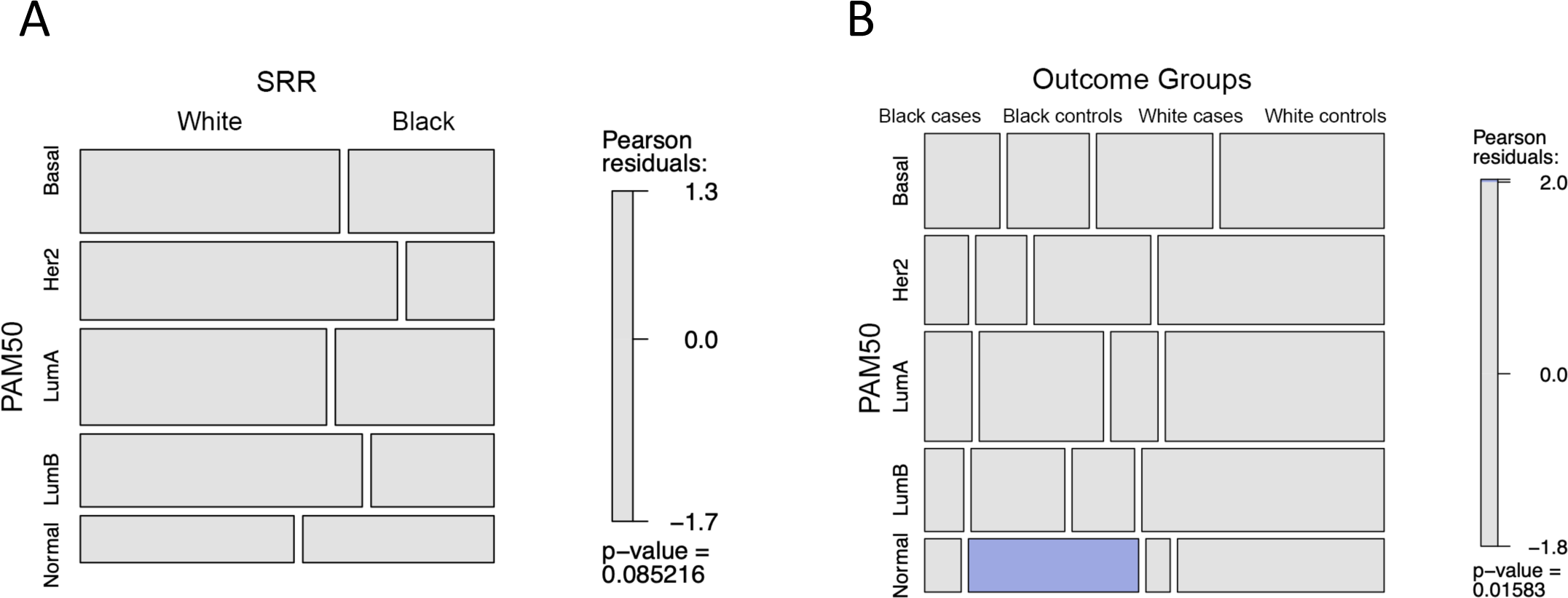
PAM50 by SRR and outcome groups. Mosaic plots showing distribution of PAM50 by SRR (A) and SRR and outcome groups (B). P-values from Chi^2^ test.

**Extended Data Figure 4:**
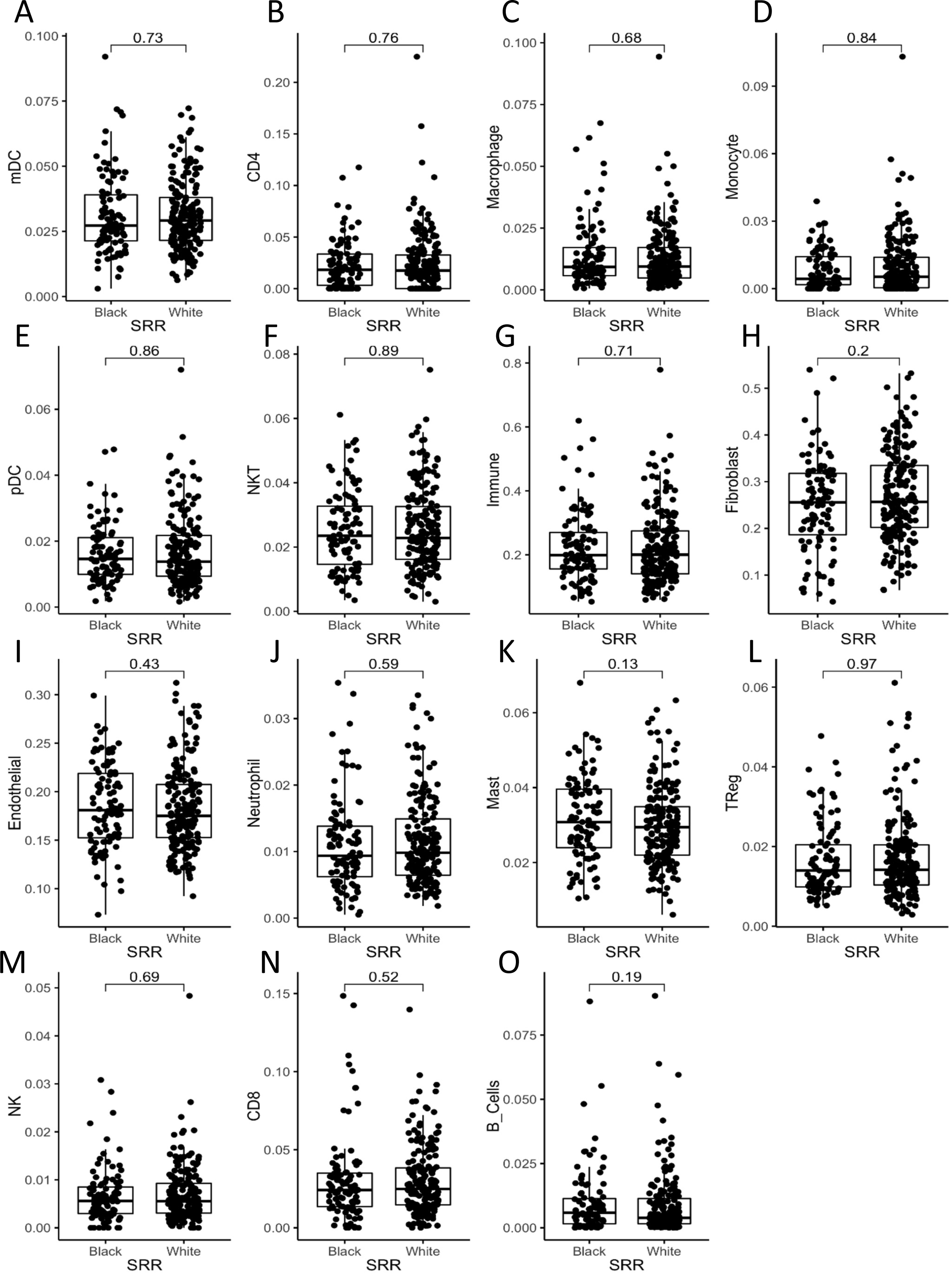
Cell type distribution by SRR. A – O) Inferred cell type distribution from RNA-seq data using CibersortX. Boxplot represents median, 0.25 and 0.75 quantiles with whiskers at 1.5x interquartile range. P-values from Wilcoxon rank-sum test. mDC = Myeloid dendritic cells. pDC = Plasmacytoid dendritic cells.

**Extended Data Figure 5:**
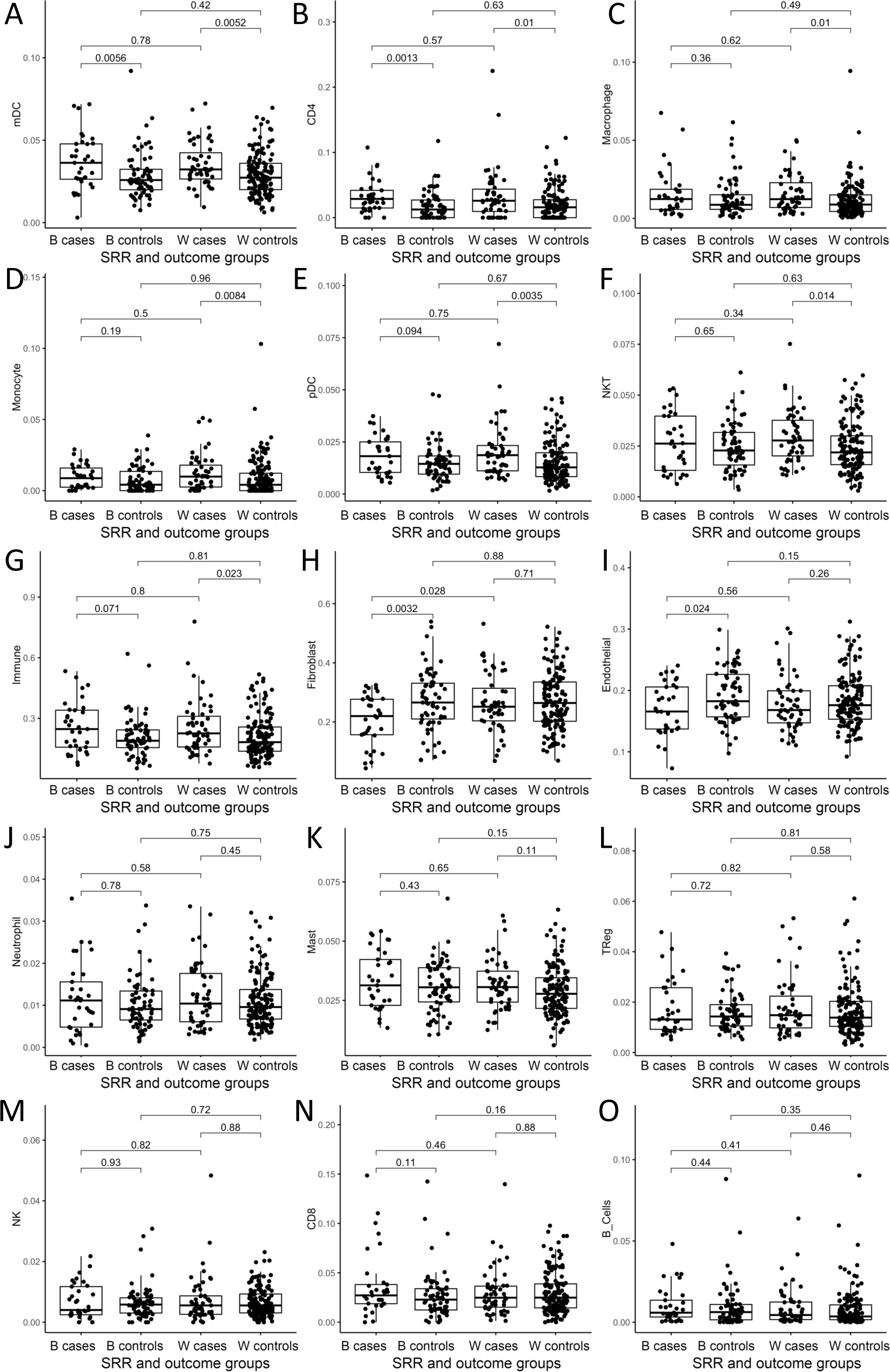
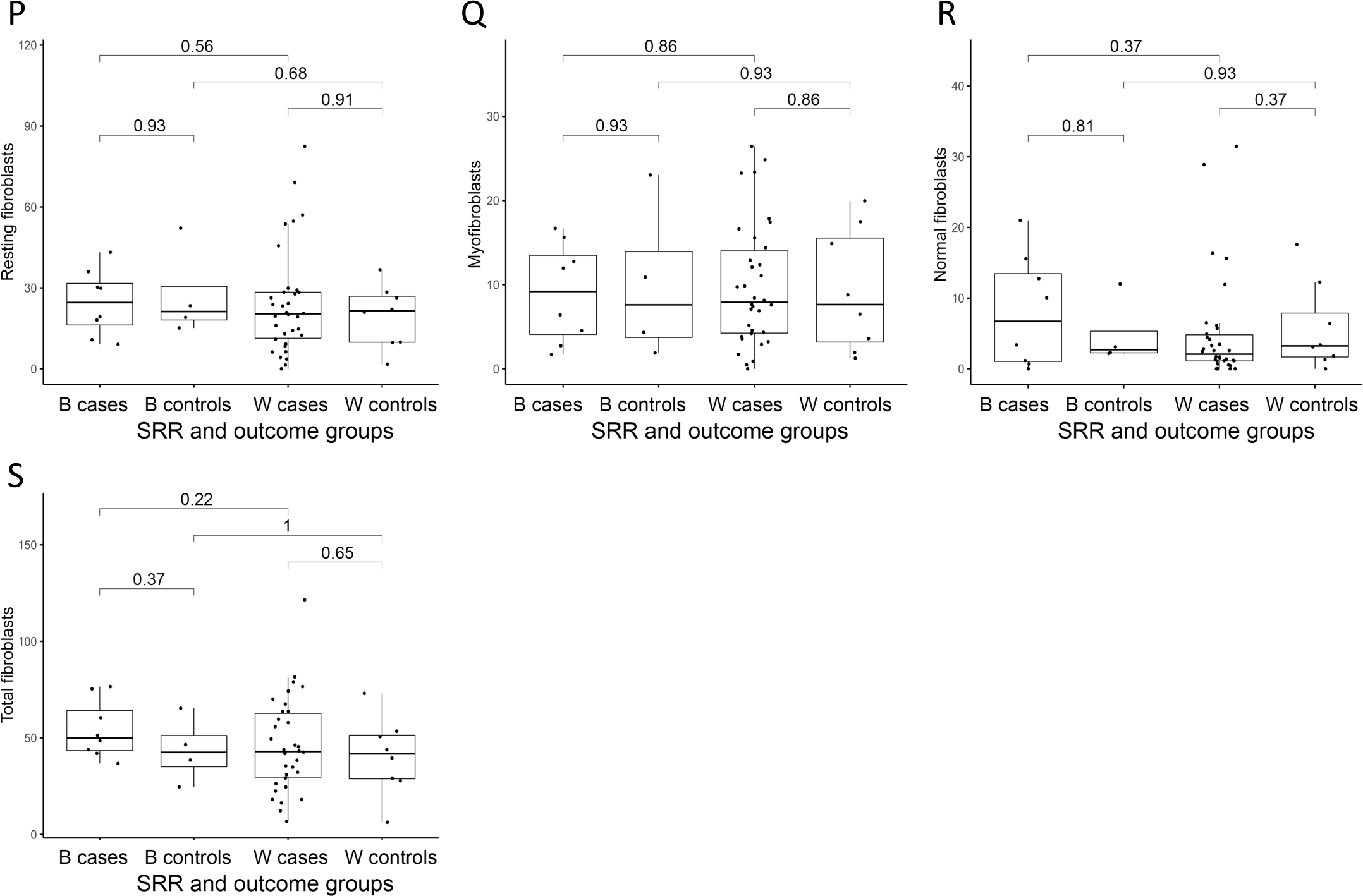
Cell type distribution by SRR and outcome groups. A – O) Inferred cell type distribution from RNA-seq data using CibersortX. mDC = Myeloid dendritic cells. pDC = Plasmacytoid dendritic cells. P – S) Fibroblast phenotypes by SRR and outcome groups in MIBI sample-level data (n=54). P) Resting fibroblasts. Q) Myofibroblasts. R) Normal fibroblasts. S) Total fibroblasts. A – S) Boxplot represents median, 0.25 and 0.75 quantiles with whiskers at 1.5x interquartile range. P-values from Wilcoxon rank-sum test. B: Black. W: White.

**Extended Data Figure 6:**
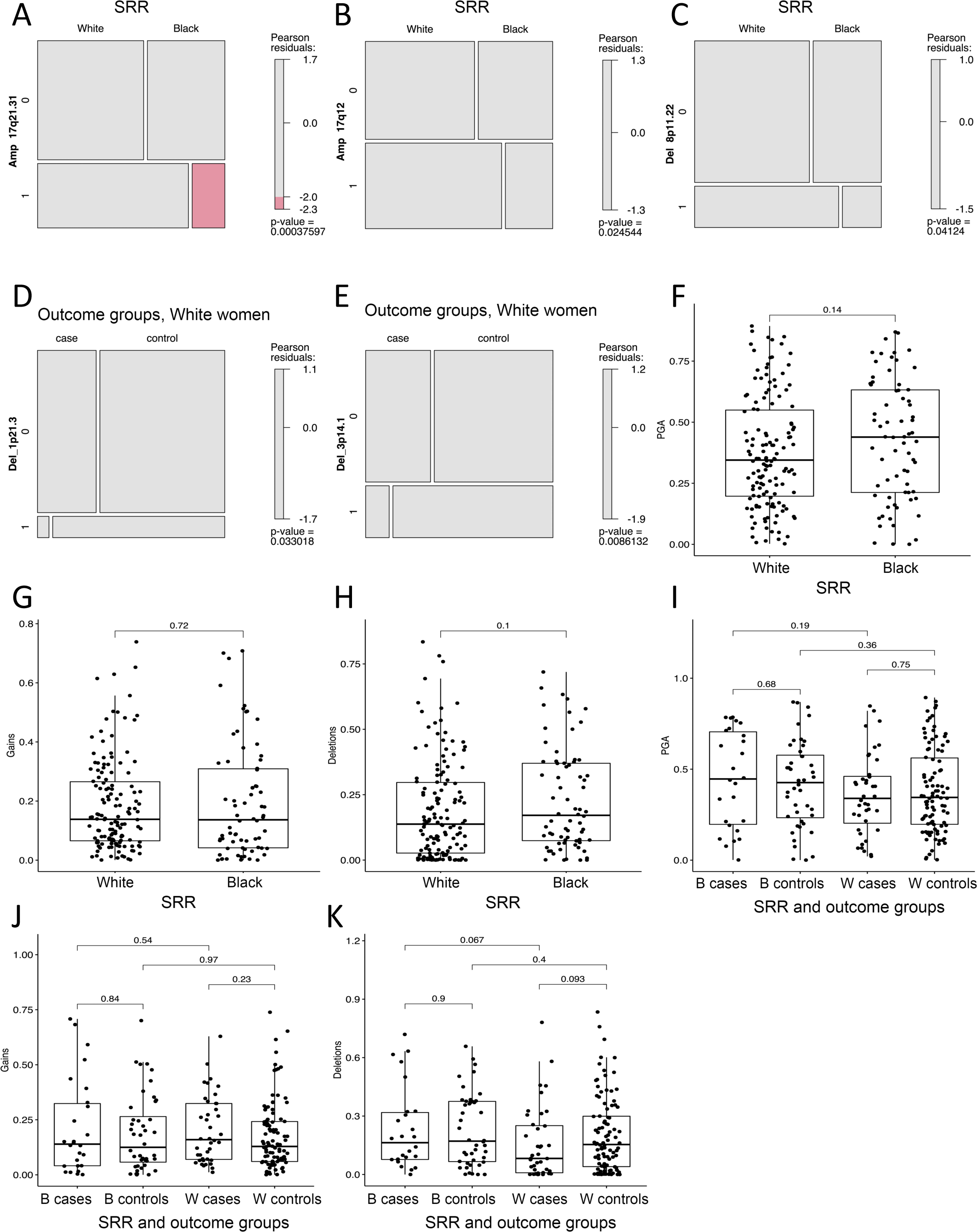
Genomic alterations by SRR and outcome groups. A – C) Mosaic plots showing distribution of significant CNVs by SRR. D – E) Mosaic plots showing distribution of significant CNVs by 5-year outcome groups in White women only. A – E) P-values from Chi^2^ test. F – H) Boxplots showing Proportion of the Genome copy number Altered (PGA, F), Gains (G), and Deletions (H) by SRR. I - K) Boxplots showing PGA (I), Gains (J), and Deletions (K) by SRR and outcome groups. F – K) Boxplots represent median, 0.25 and 0.75 quantiles with whiskers at 1.5x interquartile range. P-values from Wilcoxon rank-sum test.

