## Supplementary Tables for "Analysis of ductal carcinoma in situ by self-reported race reveals molecular differences related to outcome"

### Table S1

| | baseMean | log2FoldChang | lfcSE | stat | pvalue | padj |
|---|---|---|---|---|---|---|
| MTND1P23 | 11.78583146 | -5.192897556 | 0.319898488 | -16.23295438 | 2.95E-59 | 3.26E-55 |
| MTCO1P12 | 11.28771166 | -1.836682442 | 0.158558669 | -11.58361413 | 4.99E-31 | 1.57E-27 |
| MTCO3P12 | 0.923229765 | -1.09946826 | 0.276953669 | -3.969863486 | 7.19E-05 | 0.009400286 |
| AL645608.7 | 11.45556467 | 0.724717282 | 0.170692612 | 4.245744878 | 2.18E-05 | 0.003912952 |
| ACAP3 | 75.60338123 | 0.293130891 | 0.074859089 | 3.915768878 | 9.01E-05 | 0.011247258 |
| ANKRD65 | 2.923120175 | 0.67822389 | 0.199038929 | 3.407493663 | 0.000655624 | 0.042545656 |
| VWA1 | 115.8326265 | -0.262868502 | 0.077206922 | -3.404727121 | 0.000662302 | 0.042545656 |
| AL031282.1 | 34.03903826 | 0.340552609 | 0.09711131 | 3.506827465 | 0.000453483 | 0.033844239 |
| SLC35E2A | 46.00178746 | 0.512048775 | 0.081703547 | 6.267154773 | 3.68E-10 | 3.25E-07 |
| MEGF6 | 67.48393391 | 0.386940852 | 0.08481775 | 4.562026865 | 5.07E-06 | 0.001286411 |
| FBXO2 | 30.7072927 | -0.511963932 | 0.147753922 | -3.464976929 | 0.000530277 | 0.037545989 |
| KAZN-AS1 | 1.253540809 | 2.443408359 | 0.589220455 | 4.146849175 | 3.37E-05 | 0.005557073 |
| CROCCP2 | 56.50333691 | -0.370548707 | 0.075779368 | -4.889836339 | 1.01E-06 | 0.000314003 |
| PAX7 | 3.25668013 | 2.023898078 | 0.499420419 | 4.052493652 | 5.07E-05 | 0.007280958 |
| PNRC2 | 76.05467256 | -0.257255663 | 0.058126603 | -4.42578186 | 9.61E-06 | 0.002060974 |
| MFSD2A | 11.45230635 | 0.648779434 | 0.183903881 | 3.527818057 | 0.000419 | 0.032139341 |
| CITED4 | 99.26773208 | -0.475558324 | 0.140310916 | -3.389318065 | 0.000700667 | 0.044098096 |
| RNU5F-1 | 24.24684438 | 1.023565864 | 0.22900308 | 4.469659808 | 7.83E-06 | 0.00184117 |
| PRPF38A | 65.26253681 | -0.17226787 | 0.045532625 | -3.783394168 | 0.000154704 | 0.016752788 |
| FGGY | 12.50928086 | -0.417013149 | 0.103978841 | -4.010557785 | 6.06E-05 | 0.008110138 |
| LRRIQ3 | 4.972300527 | -0.7061773 | 0.16985293 | -4.157580915 | 3.22E-05 | 0.00545864 |
| PRKACB | 128.127692 | 0.467973504 | 0.127341353 | 3.674953124 | 0.000237893 | 0.022174246 |
| GBP7 | 0.663775808 | 1.305021072 | 0.380132306 | 3.433070677 | 0.000596787 | 0.039709671 |
| PLPPR4 | 9.289930191 | 0.522088708 | 0.14035931 | 3.719658549 | 0.000199492 | 0.019807366 |
| NBPF26 | 4.2229726 | 0.617279359 | 0.174227739 | 3.542945358 | 0.000395685 | 0.031068743 |
| NOTCH2NLA | 3.047302502 | -0.810711428 | 0.183933557 | -4.40763199 | 1.05E-05 | 0.002198726 |
| S100A1 | 54.14090175 | -0.703807155 | 0.205977629 | -3.416910661 | 0.000633361 | 0.041641575 |
| AL590560.3 | 3.474862117 | 0.868373539 | 0.189931174 | 4.572043236 | 4.83E-06 | 0.001255266 |
| LINC01133 | 4.491308877 | 0.781475667 | 0.216844324 | 3.603855758 | 0.000313531 | 0.026940519 |
| RGS5 | 581.7978868 | 0.411566116 | 0.122564381 | 3.357958594 | 0.000785204 | 0.047423682 |
| FMO4 | 11.39971732 | -0.589208428 | 0.160110563 | -3.680009731 | 0.000233225 | 0.02201784 |
| AC119673.3 | 8.763670674 | 3.056988763 | 0.310976339 | 9.830293753 | 8.34E-23 | 1.67E-19 |

| | | | | | | |
|---|---|---|---|---|---|---|
| C4BPB | 1.662893145 | -1.426510993 | 0.345227681 | -4.132087526 | 3.59E-05 | 0.005796605 |
| C4BPA | 1.69463658 | -2.564066475 | 0.396688161 | -6.463682874 | 1.02E-10 | 1.07E-07 |
| FLVCR1-DT | 7.584273677 | 0.520036695 | 0.12957383 | 4.013439241 | 5.98E-05 | 0.008060576 |
| LINC01139 | 1.589958909 | -1.816597476 | 0.35275897 | -5.14968471 | 2.61E-07 | 0.00010863 |
| EFCAB2 | 24.26354293 | -0.339279665 | 0.098976154 | -3.42789301 | 0.000608285 | 0.040112319 |
| RN7SL832P | 8.24934362 | 0.606695717 | 0.121460242 | 4.995014895 | 5.88E-07 | 0.000206292 |
| DHX57 | 48.54677354 | 0.187293492 | 0.055549182 | 3.371669655 | 0.00074714 | 0.045975128 |
| NRXN1 | 7.568565731 | -0.949446337 | 0.206868424 | -4.589614592 | 4.44E-06 | 0.001167839 |
| DQX1 | 1.984329849 | 1.271066643 | 0.361214223 | 3.518872075 | 0.000433386 | 0.032900075 |
| CAPG | 171.701926 | -0.282846206 | 0.082901763 | -3.411823796 | 0.000645298 | 0.042175386 |
| POLR1A | 57.31496174 | 0.336418769 | 0.052012394 | 6.468050095 | 9.93E-11 | 1.07E-07 |
| ANKRD36B | 36.56496467 | -0.640408771 | 0.099629623 | -6.427895143 | 1.29E-10 | 1.30E-07 |
| IL18R1 | 9.502237264 | 0.451767222 | 0.132955905 | 3.397872575 | 0.00067912 | 0.043110488 |
| LINC01123 | 1.074251121 | -1.524893191 | 0.293537956 | -5.194875683 | 2.05E-07 | 8.87E-05 |
| MTLN | 18.46511477 | -0.344977299 | 0.099279795 | -3.474798663 | 0.000511237 | 0.036549302 |
| MIR4435-2HG | 54.29614488 | 0.313072202 | 0.077602268 | 4.034317695 | 5.48E-05 | 0.007705283 |
| MTND3P10 | 32.02060232 | -2.35363016 | 0.384831782 | -6.115997357 | 9.60E-10 | 6.84E-07 |
| HS6ST1 | 87.55870746 | -0.306736854 | 0.070732882 | -4.336552485 | 1.45E-05 | 0.002829503 |
| SCN1A | 1.807426955 | -1.765231945 | 0.362485942 | -4.869794224 | 1.12E-06 | 0.000342762 |
| SCN7A | 16.29136102 | 0.569383911 | 0.149624561 | 3.805417426 | 0.000141565 | 0.015558794 |
| MTND3P17 | 1.156244364 | 1.083235365 | 0.235247652 | 4.604659616 | 4.13E-06 | 0.001099602 |
| SNORC | 13.39877975 | -0.996842282 | 0.194306104 | -5.130267449 | 2.89E-07 | 0.000113983 |
| D2HGDH | 78.26617699 | 0.281759705 | 0.073394635 | 3.838968676 | 0.000123552 | 0.014365218 |
| LRRN1 | 17.20349868 | -0.595104957 | 0.150790647 | -3.94656412 | 7.93E-05 | 0.010065459 |
| HMGB1P5 | 7.950451582 | -2.604212133 | 0.222467425 | -11.70603804 | 1.19E-31 | 4.37E-28 |
| AC112220.2 | 22.98826618 | -0.370067481 | 0.09120604 | -4.057488731 | 4.96E-05 | 0.007209109 |
| CTNNB1 | 386.4825452 | -0.156608974 | 0.045289691 | -3.457938717 | 0.000544325 | 0.038295184 |
| CLEC3B | 4.565862417 | -0.810401678 | 0.198640497 | -4.079740495 | 4.51E-05 | 0.00686893 |
| NBEAL2 | 87.21305963 | 0.269089042 | 0.064196883 | 4.191621605 | 2.77E-05 | 0.004743018 |
| SNORD13P3 | 9.6519365 | -1.364090186 | 0.282223955 | -4.833360749 | 1.34E-06 | 0.000390218 |
| COL7A1 | 45.52652881 | 0.39814529 | 0.11086307 | 3.591324758 | 0.000329001 | 0.027334187 |
| ROBO2 | 7.515064681 | 1.40044342 | 0.266890507 | 5.247258279 | 1.54E-07 | 6.96E-05 |
| AC074043.1 | 0.699115479 | -1.430786827 | 0.374919325 | -3.816252541 | 0.000135494 | 0.014984215 |
| IFT57 | 46.26401605 | -0.180310881 | 0.051829362 | -3.478933033 | 0.000503414 | 0.036106905 |
| ROPN1 | 1.720452771 | -0.99512177 | 0.292645122 | -3.400438602 | 0.000672778 | 0.042953793 |

| | | | | | | |
|---|---|---|---|---|---|---|
| AGTR1 | 14.9438782 | -1.138730058 | 0.29706103 | -3.833320235 | 0.000126425 | 0.014546141 |
| ARHGEF26 | 51.89983691 | 0.364017956 | 0.106591619 | 3.415071098 | 0.000637654 | 0.041799427 |
| SPTSSB | 41.38940747 | 0.885642079 | 0.243348783 | 3.639393911 | 0.00027328 | 0.024342901 |
| XXYLT1 | 31.24554656 | 0.263930861 | 0.071147067 | 3.709652041 | 0.000207544 | 0.020286995 |
| MELTF | 20.48115341 | -0.488301165 | 0.109915245 | -4.442524479 | 8.89E-06 | 0.002034125 |
| DLG1 | 167.7046056 | 0.170305573 | 0.045403457 | 3.750938429 | 0.000176174 | 0.018444829 |
| PCGF3 | 60.58645855 | 0.222188839 | 0.059742915 | 3.719082669 | 0.000199948 | 0.019807366 |
| DGKQ | 26.16415038 | 0.314036003 | 0.070862716 | 4.431611153 | 9.35E-06 | 0.002052228 |
| UVSSA | 66.88106332 | 0.253476954 | 0.072149019 | 3.513241869 | 0.000442674 | 0.03337583 |
| MXD4 | 194.0516952 | 0.163760643 | 0.047443359 | 3.451708459 | 0.000557049 | 0.038427089 |
| AC093323.3 | 1.187795595 | 0.857124013 | 0.247588852 | 3.461884509 | 0.000536407 | 0.037858701 |
| SH3TC1 | 26.87906851 | 0.321691681 | 0.092204196 | 3.488904996 | 0.000485003 | 0.035076667 |
| AC108519.1 | 5.229494474 | -0.991281636 | 0.183604084 | -5.399017371 | 6.70E-08 | 3.27E-05 |
| AC093809.1 | 1.90266388 | -1.012281994 | 0.229805721 | -4.40494689 | 1.06E-05 | 0.002205136 |
| STIM2-AS1 | 1.767637531 | -1.049277314 | 0.223919014 | -4.685967905 | 2.79E-06 | 0.000769429 |
| HOPX | 23.42628697 | 0.616791382 | 0.124909352 | 4.937911961 | 7.90E-07 | 0.000260356 |
| SMR3B | 1.104990692 | 2.352819931 | 0.607505581 | 3.87291904 | 0.00010754 | 0.013064547 |
| ALB | 9.238176499 | -1.163944029 | 0.313608712 | -3.711453112 | 0.000206073 | 0.02023269 |
| AC114801.3 | 5.06473098 | -2.314261992 | 0.252523856 | -9.164528157 | 4.98E-20 | 8.34E-17 |
| FAM198B-AS1 | 25.53183782 | 0.526989472 | 0.128288193 | 4.107856366 | 3.99E-05 | 0.006300527 |
| AC098679.5 | 0.953526107 | 1.013700367 | 0.297270651 | 3.410025054 | 0.000649569 | 0.042329296 |
| NAF1 | 36.42292263 | -0.332675358 | 0.09504626 | -3.5001415 | 0.000465011 | 0.034333838 |
| NPY1R | 254.477085 | -1.028512198 | 0.306620102 | -3.354353453 | 0.000795507 | 0.047423682 |
| SCRG1 | 6.620486288 | -0.646961406 | 0.173304361 | -3.73309362 | 0.000189142 | 0.019281193 |
| CENPU | 42.77278985 | 0.442817029 | 0.128826859 | 3.437303618 | 0.000587537 | 0.039450672 |
| PDCD6 | 12.02026752 | 0.550533919 | 0.102346287 | 5.379129361 | 7.48E-08 | 3.52E-05 |
| IRX2 | 161.9591829 | -0.498423865 | 0.148552071 | -3.35521318 | 0.000793038 | 0.047423682 |
| C5orf38 | 45.81805628 | -0.588536128 | 0.166521434 | -3.534296536 | 0.000408862 | 0.031581022 |
| AC008957.3 | 5.086878315 | 0.756686576 | 0.206241384 | 3.668936669 | 0.000243561 | 0.022607205 |
| NDUFAF2 | 30.3668691 | -0.262915613 | 0.068582637 | -3.833559429 | 0.000126302 | 0.014546141 |
| AC091429.1 | 4.159531085 | -2.482246865 | 0.389717457 | -6.369349947 | 1.90E-10 | 1.82E-07 |
| NR2F1-AS1 | 27.87859083 | 0.371105949 | 0.106955741 | 3.469715087 | 0.000521011 | 0.037127896 |
| LINC01554 | 1.482793285 | -1.243060162 | 0.286569298 | -4.337729722 | 1.44E-05 | 0.002829503 |
| MTND4P35 | 4.336802248 | -0.542891376 | 0.144907742 | -3.746462195 | 0.000179346 | 0.018688363 |
| MTND3P19 | 403.1539213 | -2.137938972 | 0.33824693 | -6.320645602 | 2.60E-10 | 2.40E-07 |

| | | | | | | |
|---|---|---|---|---|---|---|
| MTCO3P22 | 0.99287504 | -0.81221704 | 0.232276182 | -3.496772825 | 0.000470923 | 0.034333838 |
| ALDH7A1 | 118.1342883 | -0.256495269 | 0.068714795 | -3.732751737 | 0.000189399 | 0.019281193 |
| ISOC1 | 36.32061138 | -0.498933039 | 0.122589014 | -4.069965347 | 4.70E-05 | 0.006971287 |
| ADAMTS19 | 4.941761694 | -0.942258791 | 0.255809738 | -3.683435975 | 0.000230111 | 0.021911139 |
| LEAP2 | 2.800342106 | -0.60461306 | 0.177478686 | -3.406679837 | 0.000657582 | 0.042545656 |
| MTND4P12 | 108.3788133 | -0.546843933 | 0.093618687 | -5.841183514 | 5.18E-09 | 3.47E-06 |
| MTND4LP30 | 0.688022688 | -1.921757915 | 0.432678203 | -4.441540849 | 8.93E-06 | 0.002034125 |
| MTND3P25 | 11.60389289 | -2.130217192 | 0.310597321 | -6.858453206 | 6.96E-12 | 8.09E-09 |
| SLC25A48 | 3.419885838 | -1.04697512 | 0.25212151 | -4.152660829 | 3.29E-05 | 0.00549985 |
| AC114296.1 | 1.853296418 | -1.016198391 | 0.250072574 | -4.063613912 | 4.83E-05 | 0.007068933 |
| SPINK1 | 2.710493418 | -1.800910133 | 0.495963345 | -3.631135546 | 0.000282177 | 0.024934282 |
| RBM22 | 73.72484654 | -0.166357928 | 0.042754218 | -3.8910296 | 9.98E-05 | 0.012319096 |
| ADRA1B | 1.913679027 | -0.794466215 | 0.230387729 | -3.448387721 | 0.000563944 | 0.038427089 |
| RPL10P9 | 9.855507402 | -2.420695796 | 0.266585664 | -9.080367487 | 1.08E-19 | 1.59E-16 |
| HEIH | 47.42193355 | 0.244867169 | 0.060992416 | 4.01471504 | 5.95E-05 | 0.008060576 |
| TRIM52-AS1 | 20.07107292 | 0.398901206 | 0.078601578 | 5.074977102 | 3.88E-07 | 0.000142687 |
| FOXC1 | 36.40529966 | -0.600186309 | 0.123347993 | -4.865797132 | 1.14E-06 | 0.000344972 |
| NQO2 | 56.41083519 | -0.239070733 | 0.063425697 | -3.769304013 | 0.000163703 | 0.017470395 |
| STK19 | 9.497038905 | 0.387558338 | 0.105936881 | 3.658389171 | 0.000253805 | 0.023132016 |
| CYP21A2 | 3.852769983 | 0.880957614 | 0.262709858 | 3.353348143 | 0.000798402 | 0.047423682 |
| FO393411.1 | 9.113596332 | 0.579520115 | 0.160865897 | 3.602504485 | 0.000315166 | 0.026940519 |
| KCNQ5 | 5.571263111 | -0.903667232 | 0.258289677 | -3.49865795 | 0.000467606 | 0.034333838 |
| MRAP2 | 1.502934524 | -0.847688759 | 0.239365482 | -3.54139934 | 0.000398011 | 0.031068743 |
| HSPD1P10 | 3.429467348 | -2.990781619 | 0.29441893 | -10.15825177 | 3.04E-24 | 7.47E-21 |
| Z99129.4 | 7.133461032 | -0.408408799 | 0.120935809 | -3.37707087 | 0.000732622 | 0.045207671 |
| HEY2 | 12.33042714 | -0.597929755 | 0.14700716 | -4.067351232 | 4.76E-05 | 0.007002927 |
| IL20RA | 44.41674285 | 0.900076218 | 0.176318049 | 5.104844465 | 3.31E-07 | 0.000126096 |
| AIG1 | 85.64474716 | -0.325913118 | 0.079599058 | -4.094434382 | 4.23E-05 | 0.006516463 |
| SNHG26 | 6.111353934 | -0.570835305 | 0.162413888 | -3.514695141 | 0.000440259 | 0.033307409 |
| CHN2 | 5.708354167 | 0.701077817 | 0.195452811 | 3.586941587 | 0.000334579 | 0.027579067 |
| LINC01176 | 8.34079239 | 0.599840467 | 0.139805165 | 4.290545813 | 1.78E-05 | 0.003281149 |
| AMPH | 13.03590141 | 0.924430035 | 0.179626677 | 5.146396126 | 2.66E-07 | 0.00010863 |
| AC099654.5 | 3.60919621 | 3.644299023 | 0.304073494 | 11.9849283 | 4.26E-33 | 1.88E-29 |
| CCDC146 | 36.8323672 | 0.296689163 | 0.086042035 | 3.448188579 | 0.00056436 | 0.038427089 |
| TFPI2 | 59.95856344 | 1.009868743 | 0.288776316 | 3.497062214 | 0.000470412 | 0.034333838 |

| | | | | | | |
|---|---|---|---|---|---|---|
| TAC1 | 10.00045721 | 0.863756494 | 0.246990921 | 3.497118401 | 0.000470313 | 0.034333838 |
| PILRB | 36.32155079 | 0.604933795 | 0.123142711 | 4.912461263 | 8.99E-07 | 0.000292186 |
| PRKRIP1 | 66.32873605 | 0.162298119 | 0.045337179 | 3.579801876 | 0.000343855 | 0.028170075 |
| POLR2J3 | 71.62011271 | -0.387984662 | 0.101617538 | -3.81808759 | 0.00013449 | 0.014984215 |
| NAPEPLD | 42.87101347 | 0.254821329 | 0.066709829 | 3.819846814 | 0.000133535 | 0.014984215 |
| LSM8 | 85.04406738 | 0.18589199 | 0.046231524 | 4.020892517 | 5.80E-05 | 0.008042381 |
| TMEM213 | 3.664288532 | -0.89950736 | 0.268816104 | -3.346181074 | 0.000819329 | 0.048137741 |
| LINC00996 | 3.02116925 | 0.879305547 | 0.263031936 | 3.342961166 | 0.000828895 | 0.048442114 |
| GIMAP5 | 4.144128355 | 0.651255081 | 0.180809427 | 3.601886755 | 0.000315916 | 0.026940519 |
| UBE3C | 110.9847504 | 0.149442388 | 0.04412939 | 3.386459418 | 0.000708007 | 0.044236016 |
| CCAR2 | 20.51172965 | -0.304054673 | 0.076712134 | -3.963579917 | 7.38E-05 | 0.009482964 |
| SLC25A37 | 125.9684665 | -0.305623148 | 0.077917434 | -3.922397478 | 8.77E-05 | 0.011004356 |
| CLU | 1012.389904 | -0.383629516 | 0.111270415 | -3.447722533 | 0.000565334 | 0.038427089 |
| NRG1 | 22.0565979 | -1.091242091 | 0.175268357 | -6.226121536 | 4.78E-10 | 3.91E-07 |
| SDCBPP2 | 11.89110956 | 0.978813501 | 0.25616064 | 3.821092508 | 0.000132862 | 0.014984215 |
| MSC-AS1 | 6.351382599 | 0.537238082 | 0.145963105 | 3.680643001 | 0.000232647 | 0.02201784 |
| TMEM64 | 22.11285187 | -0.503614204 | 0.146160794 | -3.445617593 | 0.000569756 | 0.038490765 |
| CPQ | 64.59028503 | -0.272468265 | 0.073166392 | -3.723953824 | 0.000196127 | 0.019693807 |
| AC103718.1 | 2.703774283 | 1.290436498 | 0.318460008 | 4.052114758 | 5.08E-05 | 0.007280958 |
| NDRG1 | 206.3361647 | -0.503631124 | 0.103991765 | -4.842990428 | 1.28E-06 | 0.000376724 |
| WDR97 | 9.79550066 | 0.62121002 | 0.142522689 | 4.358674557 | 1.31E-05 | 0.002651985 |
| AC084125.2 | 4.942480815 | 0.714816286 | 0.18910896 | 3.779917601 | 0.00015688 | 0.016905571 |
| FREM1 | 9.356810863 | 0.756552401 | 0.15833483 | 4.778180519 | 1.77E-06 | 0.00050098 |
| SNAPC3 | 70.01898648 | 0.167353113 | 0.049554241 | 3.377170319 | 0.000732357 | 0.045207671 |
| MIR31HG | 1.293083316 | -0.940846313 | 0.264398589 | -3.558439231 | 0.000373065 | 0.029752277 |
| TMEM215 | 1.180987417 | 1.728487152 | 0.452273598 | 3.821773281 | 0.000132495 | 0.014984215 |
| LERFS | 2.161008471 | -1.418396809 | 0.398941754 | -3.555398233 | 0.000377407 | 0.029990281 |
| AL691447.2 | 1.135542827 | -1.16390347 | 0.293337638 | -3.967794513 | 7.25E-05 | 0.009426469 |
| AL354797.1 | 4.102616004 | 0.500408446 | 0.149667332 | 3.343471405 | 0.000827372 | 0.048442114 |
| CUTALP | 54.95489314 | -0.448155061 | 0.07459624 | -6.007743284 | 1.88E-09 | 1.30E-06 |
| DBH-AS1 | 1.988110208 | 0.814063199 | 0.239111262 | 3.404537251 | 0.000662763 | 0.042545656 |
| OLFM1 | 41.03188909 | 0.381153768 | 0.104213379 | 3.657436048 | 0.000254751 | 0.023132016 |
| OBP2A | 3.79362762 | 1.461139672 | 0.407154903 | 3.588657929 | 0.000332385 | 0.027500781 |
| PTGDS | 213.9490111 | 0.527548288 | 0.153063048 | 3.446607764 | 0.000567672 | 0.038467622 |
| CLIC3 | 12.99242257 | 0.602668265 | 0.149925169 | 4.019793797 | 5.82E-05 | 0.008042381 |

| | | | | | | |
|---|---|---|---|---|---|---|
| ZMYND19 | 42.00513883 | 0.243787545 | 0.068358655 | 3.566301061 | 0.000362055 | 0.028978863 |
| USP6NL | 33.50661834 | 0.247161854 | 0.073059208 | 3.383034948 | 0.000716895 | 0.044485757 |
| C1DP1 | 0.74674315 | -1.472869206 | 0.332145971 | -4.434403346 | 9.23E-06 | 0.002052228 |
| MSMB | 13.12524311 | 1.18452597 | 0.289362323 | 4.093573617 | 4.25E-05 | 0.006516463 |
| MICU1 | 48.59965854 | -0.194214673 | 0.052322902 | -3.711848272 | 0.000205751 | 0.02023269 |
| MYOZ1 | 1.72087569 | -0.717572766 | 0.208048595 | -3.449063263 | 0.000562535 | 0.038427089 |
| LDB3 | 7.790919005 | 0.631013919 | 0.139920811 | 4.509793194 | 6.49E-06 | 0.001575279 |
| LIPA | 80.38158423 | -0.352858607 | 0.1049913 | -3.360836611 | 0.000777068 | 0.047264822 |
| PPP1R3C | 20.61224808 | -0.704320573 | 0.199987035 | -3.521831169 | 0.000428577 | 0.032647222 |
| MYOF | 212.9900448 | -0.240718273 | 0.071105814 | -3.385352886 | 0.000710868 | 0.044236016 |
| GSTO1 | 95.4856524 | -0.217524068 | 0.063892251 | -3.404545378 | 0.000662743 | 0.042545656 |
| CASP7 | 22.81009255 | -0.358904151 | 0.078740641 | -4.558054735 | 5.16E-06 | 0.001296079 |
| ADRB1 | 7.8956798 | 1.195116771 | 0.243857231 | 4.900887158 | 9.54E-07 | 0.000305448 |
| ATE1-AS1 | 2.606392986 | -0.741569323 | 0.206495102 | -3.591219923 | 0.000329134 | 0.027334187 |
| INSYN2A | 18.2700497 | -1.24494054 | 0.305779083 | -4.071372473 | 4.67E-05 | 0.006971287 |
| PGGHG | 213.0632609 | 0.651604721 | 0.182429194 | 3.57182262 | 0.000354505 | 0.028686377 |
| MUC5B | 30.45615522 | 1.133630546 | 0.337367679 | 3.360222734 | 0.000778797 | 0.047264822 |
| HBB | 298.9030569 | -1.082012181 | 0.196234593 | -5.513870751 | 3.51E-08 | 1.99E-05 |
| MTRNR2L8 | 19.75818978 | 1.061752496 | 0.183943231 | 5.772174881 | 7.83E-09 | 5.00E-06 |
| TPH1 | 7.974583859 | 0.670121242 | 0.185086128 | 3.620591392 | 0.00029393 | 0.025766732 |
| SVIP | 147.1179613 | -0.260017368 | 0.07100534 | -3.661941003 | 0.000250312 | 0.022968458 |
| ACCS | 29.21245904 | 0.327422371 | 0.085630818 | 3.823651104 | 0.00013149 | 0.014984215 |
| CREB3L1 | 66.41275578 | 0.53805999 | 0.143811396 | 3.741428041 | 0.000182978 | 0.018888587 |
| MS4A7 | 158.4658946 | -0.450293407 | 0.127112357 | -3.542483329 | 0.000396378 | 0.031068743 |
| FEN1 | 17.74757827 | -0.412277133 | 0.112114975 | -3.677270898 | 0.000235743 | 0.022067728 |
| ASRGL1 | 16.43354739 | -0.511602367 | 0.149209153 | -3.428759953 | 0.000606346 | 0.040104131 |
| NEAT1 | 5615.230921 | 0.421823995 | 0.102727432 | 4.106244916 | 4.02E-05 | 0.006300527 |
| AP002387.2 | 7.343111977 | 0.674436515 | 0.156783463 | 4.301706954 | 1.69E-05 | 0.003173005 |
| XRRA1 | 58.95189715 | 0.346188639 | 0.083296118 | 4.156119727 | 3.24E-05 | 0.00545864 |
| GUCY1A2 | 72.6423217 | 0.46386451 | 0.131172725 | 3.536287826 | 0.000405792 | 0.031453888 |
| NCAM1 | 10.24885974 | -0.908617168 | 0.166531916 | -5.45611428 | 4.87E-08 | 2.56E-05 |
| RPL5P30 | 1.964133671 | -0.721902055 | 0.214402574 | -3.367040052 | 0.000759797 | 0.046366478 |
| APLP2 | 679.0852108 | -0.258916132 | 0.0737166 | -3.512317883 | 0.000444216 | 0.033378173 |
| TPI1 | 316.030266 | 0.343628194 | 0.060687677 | 5.662240047 | 1.49E-08 | 9.17E-06 |
| CLSTN3 | 124.4360129 | 0.2638045 | 0.07240754 | 3.643329139 | 0.000269134 | 0.024070633 |

| | | | | | | |
|---|---|---|---|---|---|---|
| HTR7P1 | 8.076924078 | 0.548579745 | 0.152542764 | 3.596235771 | 0.000322855 | 0.027015887 |
| PRR13 | 65.56167079 | 0.505400064 | 0.104013579 | 4.858981592 | 1.18E-06 | 0.000352235 |
| LACRT | 2.519820235 | 2.870705257 | 0.778269063 | 3.688576863 | 0.000225512 | 0.021754513 |
| NTS | 4.820653762 | 2.890423198 | 0.582737764 | 4.960075316 | 7.05E-07 | 0.000239974 |
| MYBPC1 | 23.53311334 | 1.092200607 | 0.325693696 | 3.353459467 | 0.000798081 | 0.047423682 |
| AC026362.2 | 3.871335202 | 0.695115797 | 0.18635235 | 3.730115542 | 0.000191392 | 0.01939468 |
| SCARB1 | 43.48683449 | -0.280283471 | 0.084033084 | -3.335394333 | 0.000851785 | 0.049258604 |
| EP400P1 | 46.60236621 | 0.263054542 | 0.07728159 | 3.403844834 | 0.000664445 | 0.042545656 |
| ANKRD20A9P | 3.536047438 | -2.133021551 | 0.41602178 | -5.12718721 | 2.94E-07 | 0.000113983 |
| DCLK1 | 110.3407004 | 0.350789878 | 0.103284832 | 3.396334892 | 0.000682947 | 0.043140827 |
| CAB39L | 59.13296601 | -0.317509181 | 0.095240713 | -3.333754773 | 0.000856822 | 0.049313709 |
| LINC00383 | 1.805787111 | 1.881292849 | 0.492673456 | 3.818539085 | 0.000134244 | 0.014984215 |
| CHMP4A | 50.10405793 | 0.331574861 | 0.07510078 | 4.415065474 | 1.01E-05 | 0.002144939 |
| FUT8-AS1 | 3.710341684 | -0.704176606 | 0.211234113 | -3.333631092 | 0.000857203 | 0.049313709 |
| ACOT1 | 3.018435037 | -1.399137729 | 0.333595264 | -4.194117479 | 2.74E-05 | 0.00472775 |
| PRIMA1 | 2.904580513 | -0.682766543 | 0.20359041 | -3.353628218 | 0.000797595 | 0.047423682 |
| INF2 | 134.2177706 | 0.237191721 | 0.06569304 | 3.61060656 | 0.000305482 | 0.026568494 |
| IGHA2 | 132.080585 | -4.327193976 | 0.252434926 | -17.14181965 | 7.24E-66 | 1.60E-61 |
| IGHG4 | 732.9995505 | -1.279477927 | 0.282777818 | -4.524675722 | 6.05E-06 | 0.001484716 |
| IGHG2 | 14.50392729 | -3.074098436 | 0.313689852 | -9.799802002 | 1.13E-22 | 2.08E-19 |
| IGHGP | 13.31773171 | 1.392385404 | 0.32824819 | 4.241867725 | 2.22E-05 | 0.003949075 |
| IGHG1 | 425.4111773 | -2.034119433 | 0.235603164 | -8.633667739 | 5.94E-18 | 8.20E-15 |
| IGHM | 548.5662579 | 0.842544708 | 0.192579217 | 4.375055205 | 1.21E-05 | 0.002483228 |
| AC087283.1 | 1.673221075 | -1.12343603 | 0.218687295 | -5.137180141 | 2.79E-07 | 0.000112018 |
| SRP14 | 206.1567047 | -0.253548303 | 0.07168799 | -3.536830968 | 0.000404959 | 0.031453888 |
| ADAL | 32.29115451 | 0.319996074 | 0.078047573 | 4.100013152 | 4.13E-05 | 0.006427029 |
| ALDH1A2 | 8.439667988 | -0.615986903 | 0.168225489 | -3.66167403 | 0.000250573 | 0.022968458 |
| MTND3P12 | 2.719414131 | 3.445120916 | 0.286954605 | 12.00580459 | 3.31E-33 | 1.83E-29 |
| HMGB1P6 | 2.186081152 | -1.41709643 | 0.229540117 | -6.173632954 | 6.67E-10 | 5.08E-07 |
| ETFA | 63.19752834 | 0.309739849 | 0.085841808 | 3.608263347 | 0.000308254 | 0.026704425 |
| UBE2Q2P2 | 12.30123257 | 0.460070783 | 0.119673177 | 3.844393498 | 0.000120851 | 0.01420063 |
| GOLGA6L9 | 5.768033505 | 0.541765854 | 0.144937208 | 3.737934926 | 0.000185538 | 0.019063809 |
| FURIN | 76.41554756 | -0.276065891 | 0.082701469 | -3.338101432 | 0.00084353 | 0.048909213 |
| LINC00923 | 2.379320765 | -0.862102347 | 0.235760641 | -3.656684778 | 0.000255498 | 0.023132016 |
| SYNM | 180.8005556 | -0.496239195 | 0.14849885 | -3.341703959 | 0.000832658 | 0.048513363 |

| | | | | | | |
|---|---|---|---|---|---|---|
| AC015712.6 | 4.92846003 | -1.400261681 | 0.182286205 | -7.681665651 | 1.57E-14 | 2.04E-11 |
| HBA1 | 148.5387923 | -0.821407685 | 0.180515117 | -4.550354011 | 5.36E-06 | 0.001329326 |
| TSC2 | 131.0993943 | 0.189011344 | 0.055655217 | 3.396111897 | 0.000683504 | 0.043140827 |
| PKD1 | 78.0286797 | 0.185705122 | 0.055351712 | 3.355002322 | 0.000793643 | 0.047423682 |
| ACSM2A | 0.818309089 | -1.045566422 | 0.301457529 | -3.468370571 | 0.000523625 | 0.03719419 |
| AC008870.1 | 1.328161193 | 2.504463066 | 0.575959307 | 4.348333356 | 1.37E-05 | 0.002730049 |
| HSD3B7 | 40.21514153 | -0.272930733 | 0.068652918 | -3.975515386 | 7.02E-05 | 0.009234433 |
| CBLN1 | 1.634282767 | 0.990965897 | 0.287187514 | 3.45058838 | 0.000559366 | 0.038427089 |
| LRRC36 | 1.915417258 | -0.819126414 | 0.229301912 | -3.572261603 | 0.000353912 | 0.028686377 |
| TPPP3 | 40.11641304 | -0.440654258 | 0.128505497 | -3.429069329 | 0.000605655 | 0.040104131 |
| SLC7A6 | 44.47229398 | 0.312596985 | 0.088180369 | 3.544972506 | 0.000392654 | 0.031068743 |
| COG4 | 62.62184353 | 0.235320354 | 0.058243657 | 4.040274335 | 5.34E-05 | 0.007560321 |
| GABARAPL2 | 66.05333514 | -0.234191878 | 0.060721139 | -3.856842662 | 0.000114861 | 0.013568959 |
| SRR | 9.636019752 | -0.464997305 | 0.101906047 | -4.563000133 | 5.04E-06 | 0.001286411 |
| VMO1 | 4.127872191 | -0.677742884 | 0.154286399 | -4.392758455 | 1.12E-05 | 0.002310707 |
| CHRNE | 4.85099446 | 0.579359151 | 0.167988028 | 3.448812147 | 0.000563058 | 0.038427089 |
| DVL2 | 24.01431859 | 0.246068068 | 0.068057516 | 3.615589877 | 0.000299664 | 0.026165566 |
| CHRNB1 | 12.81155596 | -0.336205981 | 0.093443756 | -3.597950219 | 0.000320735 | 0.026940519 |
| TP53 | 41.89511346 | -0.281553921 | 0.077453416 | -3.635138843 | 0.000277831 | 0.024648851 |
| MTRNR2L1 | 56.03199093 | 0.7781404 | 0.146430875 | 5.314045974 | 1.07E-07 | 4.93E-05 |
| MTND1P15 | 0.997340502 | -1.362417722 | 0.361822721 | -3.765428881 | 0.000166263 | 0.017658297 |
| CORO6 | 11.43652153 | 0.874289835 | 0.217700101 | 4.016028621 | 5.92E-05 | 0.008060576 |
| AC104984.4 | 49.57609717 | 0.657927853 | 0.10526673 | 6.250102526 | 4.10E-10 | 3.49E-07 |
| AC130324.3 | 1.514114275 | 1.063126166 | 0.275177172 | 3.863424266 | 0.000111809 | 0.013279371 |
| CCT6B | 11.76768522 | -0.433751586 | 0.124099642 | -3.495188043 | 0.000473728 | 0.034424745 |
| RN7SL301P | 0.626627418 | -1.519504042 | 0.37516298 | -4.050250484 | 5.12E-05 | 0.00729186 |
| AC243829.1 | 1.43896332 | -1.035659725 | 0.279533969 | -3.704951243 | 0.000211431 | 0.020575908 |
| SYNRG | 137.6615081 | 0.351987384 | 0.090889291 | 3.872704674 | 0.000107634 | 0.013064547 |
| DDX52 | 187.3072484 | 0.38223091 | 0.110574048 | 3.456786791 | 0.000546657 | 0.038337154 |
| SRCIN1 | 21.78789601 | 0.742390714 | 0.193358924 | 3.839443767 | 0.000123313 | 0.014365218 |
| MLLT6 | 337.1025599 | 0.546601263 | 0.125629776 | 4.350889413 | 1.36E-05 | 0.002722945 |
| PCGF2 | 118.4382279 | 0.501488894 | 0.136058811 | 3.685824475 | 0.000227963 | 0.021800607 |
| PSMB3 | 200.4163675 | 0.5280379 | 0.143017503 | 3.692120824 | 0.000222392 | 0.021547616 |
| PIP4K2B | 88.69228707 | 0.505830281 | 0.127619235 | 3.963589678 | 7.38E-05 | 0.009482964 |
| CWC25 | 88.88510319 | 0.501860391 | 0.117520473 | 4.270408203 | 1.95E-05 | 0.00353303 |

| | | | | | | |
|---|---|---|---|---|---|---|
| RPL23 | 1600.87173 | 0.474123164 | 0.130664796 | 3.628545552 | 0.000285022 | 0.025085383 |
| LASP1 | 464.0543692 | 0.568637926 | 0.132089087 | 4.304957643 | 1.67E-05 | 0.003153494 |
| LINC00672 | 28.43343417 | 1.092187436 | 0.199251049 | 5.481463924 | 4.22E-08 | 2.27E-05 |
| RPL19 | 1528.955416 | 0.593275765 | 0.137296853 | 4.321117006 | 1.55E-05 | 0.003008277 |
| FBXL20 | 121.4867152 | 0.656735097 | 0.130903567 | 5.016938151 | 5.25E-07 | 0.000187066 |
| MED1 | 350.4403703 | 0.545522645 | 0.147948466 | 3.687247729 | 0.000226693 | 0.021773336 |
| CDK12 | 411.8393456 | 0.493897243 | 0.139993286 | 3.52800665 | 0.000418702 | 0.032139341 |
| AC009283.1 | 22.4576005 | 0.713619574 | 0.213588949 | 3.341088471 | 0.000834506 | 0.048513363 |
| STARD3 | 256.1865217 | 0.739158174 | 0.172899334 | 4.275078208 | 1.91E-05 | 0.003488374 |
| TCAP | 10.29148712 | 0.944958903 | 0.262114115 | 3.605143141 | 0.000311981 | 0.02692176 |
| ERBB2 | 1176.336369 | 0.898699119 | 0.213407811 | 4.21118193 | 2.54E-05 | 0.004489564 |
| MIEN1 | 467.2101911 | 0.7376278 | 0.189937806 | 3.883522798 | 0.000102954 | 0.0126353 |
| GRB7 | 246.7335408 | 0.804171214 | 0.223445927 | 3.598952218 | 0.000319502 | 0.026940519 |
| GSDMB | 88.81418502 | 1.116610248 | 0.182572243 | 6.115991277 | 9.60E-10 | 6.84E-07 |
| ORMDL3 | 330.9291448 | 0.878102367 | 0.169543802 | 5.179206535 | 2.23E-07 | 9.47E-05 |
| PSMD3 | 211.8079658 | 0.673201142 | 0.173987531 | 3.869249352 | 0.000109171 | 0.013178663 |
| KRT15 | 273.0397365 | -0.908162004 | 0.210604943 | -4.312159015 | 1.62E-05 | 0.003105574 |
| NBR2 | 6.292742011 | -0.498448456 | 0.144274531 | -3.454861047 | 0.000550576 | 0.038427089 |
| AC003102.1 | 19.49946276 | 0.414249165 | 0.115784516 | 3.5777596 | 0.000346552 | 0.028249739 |
| KANSL1-AS1 | 7.213772737 | 0.678256525 | 0.161327225 | 4.204228539 | 2.62E-05 | 0.004593072 |
| ARL17B | 15.1074149 | 0.77852191 | 0.199511544 | 3.902139665 | 9.53E-05 | 0.011833095 |
| MRPL10 | 30.77218106 | -0.302748825 | 0.078350566 | -3.864028563 | 0.000111532 | 0.013279371 |
| HOXB8 | 2.640605271 | 0.877152245 | 0.259069892 | 3.385774545 | 0.000709777 | 0.044236016 |
| MYCBPAP | 4.506671715 | 0.908955165 | 0.183268897 | 4.959680452 | 7.06E-07 | 0.000239974 |
| SNHG25 | 667.5864485 | 0.601011928 | 0.163440087 | 3.677261436 | 0.000235751 | 0.022067728 |
| ROCR | 6.350652188 | -1.072690996 | 0.241864578 | -4.435089285 | 9.20E-06 | 0.002052228 |
| TRIM65 | 34.12609284 | 0.305550396 | 0.086277467 | 3.541485454 | 0.000397881 | 0.031068743 |
| ENGASE | 39.22464853 | 0.328088705 | 0.095508709 | 3.435170556 | 0.000592181 | 0.039535951 |
| RAB40B | 29.17212042 | -0.284234634 | 0.080697743 | -3.522212944 | 0.00042796 | 0.032647222 |
| ADCYAP1 | 6.06644423 | 1.010595335 | 0.210444992 | 4.802182866 | 1.57E-06 | 0.00045027 |
| GNAL | 16.86137805 | -0.461760487 | 0.10742334 | -4.298511706 | 1.72E-05 | 0.003192035 |
| TPGS2 | 74.48317369 | -0.252023613 | 0.05851632 | -4.306894417 | 1.66E-05 | 0.003152968 |
| SLC14A1 | 7.546991704 | -0.863783031 | 0.19284832 | -4.479079893 | 7.50E-06 | 0.001800068 |
| TNFRSF11A | 12.50440686 | -0.475119705 | 0.133106923 | -3.569459007 | 0.000357719 | 0.028840783 |
| SERPINB5 | 16.13083566 | -0.663073834 | 0.148858822 | -4.454380507 | 8.41E-06 | 0.001956467 |

| | | | | | | |
|---|---|---|---|---|---|---|
| TCF3 | 106.2033465 | 0.211612202 | 0.059311681 | 3.567799761 | 0.000359991 | 0.028918439 |
| CLPP | 71.86817105 | -0.224192269 | 0.059897103 | -3.742956806 | 0.000181868 | 0.018862136 |
| KEAP1 | 54.71151818 | -0.199783763 | 0.057906905 | -3.450085316 | 0.000560409 | 0.038427089 |
| SSBP4 | 119.3941271 | 0.20171353 | 0.058565094 | 3.444262032 | 0.00057262 | 0.038566337 |
| SLC7A10 | 3.160996202 | 0.924007968 | 0.265487565 | 3.480419011 | 0.00050063 | 0.036024172 |
| HAMP | 4.77571127 | -1.055193834 | 0.194290388 | -5.431014088 | 5.60E-08 | 2.85E-05 |
| IGFLR1 | 6.026886245 | 0.860232715 | 0.168855971 | 5.094476154 | 3.50E-07 | 0.000130939 |
| PROSER3 | 50.50568883 | 0.285745041 | 0.068121699 | 4.194625891 | 2.73E-05 | 0.00472775 |
| AC008982.2 | 19.7662161 | 0.350105431 | 0.0928111 | 3.772236648 | 0.000161791 | 0.017350089 |
| FCGBP | 37.33246508 | -0.838003175 | 0.160405133 | -5.224291502 | 1.75E-07 | 7.72E-05 |
| MIA | 21.10871818 | -0.7539393 | 0.170235393 | -4.428804653 | 9.48E-06 | 0.002052228 |
| ARHGEF1 | 92.45163249 | 0.225027747 | 0.067248957 | 3.346189414 | 0.000819304 | 0.048137741 |
| PLA2G4C | 18.67074155 | -0.408016162 | 0.105564395 | -3.865092592 | 0.000111047 | 0.013279371 |
| FAM71E1 | 4.307416594 | -0.616831263 | 0.161645456 | -3.81595177 | 0.000135659 | 0.014984215 |
| ZNF761 | 12.33796747 | 0.338084656 | 0.094451193 | 3.579464128 | 0.0003443 | 0.028170075 |
| LAIR2 | 1.47643804 | -2.273775735 | 0.476741497 | -4.769410149 | 1.85E-06 | 0.000516667 |
| COX6B2 | 2.985727975 | 1.203926087 | 0.323592206 | 3.720503974 | 0.000198826 | 0.019807366 |
| AIMP1P1 | 0.958120639 | -2.62753971 | 0.42489776 | -6.183934012 | 6.25E-10 | 4.93E-07 |
| CST2 | 3.425020714 | 1.160176009 | 0.261884704 | 4.43010223 | 9.42E-06 | 0.002052228 |
| FRG1CP | 18.57613993 | -0.62668578 | 0.114162129 | -5.489436699 | 4.03E-08 | 2.23E-05 |
| CHMP4B | 177.1919255 | -0.167833198 | 0.048111451 | -3.488425171 | 0.000485875 | 0.035076667 |
| MMP24OS | 116.4816365 | 0.330657461 | 0.079988166 | 4.133829766 | 3.57E-05 | 0.005796605 |
| AL391095.2 | 2.199598989 | -0.770652019 | 0.191440116 | -4.025551354 | 5.68E-05 | 0.007947443 |
| CCN5 | 71.52571107 | -0.633622634 | 0.15330649 | -4.133045091 | 3.58E-05 | 0.005796605 |
| SMIM25 | 7.209223178 | -0.71653337 | 0.190770839 | -3.755990024 | 0.000172658 | 0.018162754 |
| BMP7 | 14.80742343 | 0.821860025 | 0.208639615 | 3.939136983 | 8.18E-05 | 0.010322838 |
| CTCFL | 1.946715258 | 0.898472482 | 0.266636763 | 3.369649679 | 0.000752638 | 0.046152876 |
| C20orf204 | 10.86504346 | 0.67571788 | 0.177859503 | 3.799166591 | 0.000145183 | 0.015858127 |
| FP565260.1 | 3.275119528 | 0.808935653 | 0.220612181 | 3.66677691 | 0.000245627 | 0.022703531 |
| GRIK1 | 4.268946377 | -1.384140831 | 0.36814638 | -3.759756734 | 0.000170079 | 0.01797707 |
| GRIK1-AS1 | 5.081746603 | -1.360811703 | 0.329922256 | -4.124643537 | 3.71E-05 | 0.00594391 |
| TMEM50B | 90.54334096 | 0.224931444 | 0.050305396 | 4.471318463 | 7.77E-06 | 0.00184117 |
| S100B | 15.32793767 | -0.520173855 | 0.148656143 | -3.499174984 | 0.0004667 | 0.034333838 |
| ATP6V1E1 | 105.3335187 | -0.193842585 | 0.057535835 | -3.369075719 | 0.000754207 | 0.046152876 |
| UBE2L3 | 73.2047855 | -0.214955542 | 0.05973426 | -3.59853024 | 0.000320021 | 0.026940519 |

| | | | | | | |
|---|---|---|---|---|---|---|
| ZDHHC8P1 | 4.905373662 | 0.615877195 | 0.178549355 | 3.449338678 | 0.000561961 | 0.038427089 |
| AP000347.1 | 18.20340071 | 0.552303097 | 0.111700001 | 4.944521842 | 7.63E-07 | 0.000255489 |
| MMP11 | 84.89573814 | 0.640128868 | 0.15943067 | 4.01509238 | 5.94E-05 | 0.008060576 |
| CRYBB3 | 1.139966601 | 0.963084445 | 0.287270028 | 3.352540642 | 0.000800735 | 0.047423682 |
| CRYBB2 | 13.19141579 | -2.765436633 | 0.301969319 | -9.158005318 | 5.29E-20 | 8.34E-17 |
| CRYBB2P1 | 8.81725463 | 0.641099924 | 0.118101221 | 5.428393723 | 5.69E-08 | 2.85E-05 |
| AL008721.2 | 2.119948687 | 1.177017362 | 0.233716694 | 5.036085959 | 4.75E-07 | 0.000172073 |
| NEFH | 4.556036757 | 0.691921806 | 0.192239415 | 3.599271284 | 0.00031911 | 0.026940519 |
| SOX10 | 20.19942381 | -0.725379849 | 0.176417109 | -4.11173188 | 3.93E-05 | 0.00624114 |
| POLDIP3 | 33.84963066 | -0.249135894 | 0.066854882 | -3.726517579 | 0.000194144 | 0.019583675 |
| FAM118A | 58.56985378 | 0.696514808 | 0.123625229 | 5.634083052 | 1.76E-08 | 1.05E-05 |
| TTC38 | 24.7426193 | -0.369367755 | 0.075535407 | -4.889994881 | 1.01E-06 | 0.000314003 |
| BX890604.2 | 17.82549355 | 0.372973694 | 0.093690944 | 3.980893743 | 6.87E-05 | 0.009136709 |
| CA5BP1 | 5.667936122 | 1.796931271 | 0.240831251 | 7.461370844 | 8.56E-14 | 1.05E-10 |
| ADGRG2 | 1.744152798 | -1.021272641 | 0.279446108 | -3.654631832 | 0.000257551 | 0.023222717 |
| BCLAF3 | 50.08091718 | 0.191994045 | 0.052686542 | 3.644081358 | 0.000268349 | 0.024070633 |
| MAP7D2 | 2.984207982 | 1.025323276 | 0.292212495 | 3.508827633 | 0.000450086 | 0.033704613 |
| EIF1AX | 211.0529002 | 0.600690741 | 0.060760512 | 9.886202737 | 4.78E-23 | 1.06E-19 |
| PDK3 | 105.2968762 | -0.287335047 | 0.062197886 | -4.619691509 | 3.84E-06 | 0.001035343 |
| MTRNR2L10 | 10.66037084 | -0.821911134 | 0.152313058 | -5.396196133 | 6.81E-08 | 3.27E-05 |
| CITED1 | 5.324646212 | -0.944803178 | 0.281780841 | -3.352971671 | 0.000799489 | 0.047423682 |
| RAB9B | 7.355128142 | -0.497368669 | 0.125873062 | -3.951351145 | 7.77E-05 | 0.009923226 |
| RNF128 | 4.323739459 | -0.824566803 | 0.242544212 | -3.399655663 | 0.000674708 | 0.042953793 |
| CLDN2 | 1.115277631 | 1.645514564 | 0.413572178 | 3.978784481 | 6.93E-05 | 0.009162936 |
| RHOXF1-AS1 | 2.936661083 | -0.760117824 | 0.221281176 | -3.435076751 | 0.000592386 | 0.039535951 |
| HSPA8P1 | 0.912040496 | -1.154535452 | 0.303965536 | -3.798244584 | 0.000145724 | 0.015858127 |
| MTND4P24 | 2.538329535 | -3.909410924 | 0.242491426 | -16.12185214 | 1.79E-58 | 1.32E-54 |
| IGSF1 | 22.05282749 | 1.151717587 | 0.329156777 | 3.498993999 | 0.000467017 | 0.034333838 |
| MIR503HG | 15.08551431 | 0.515841075 | 0.152349343 | 3.385909415 | 0.000709428 | 0.044236016 |
| IDS | 33.03174748 | 0.2615709 | 0.078073994 | 3.350294865 | 0.000807256 | 0.047682051 |
| GABRE | 33.33632129 | 0.723930775 | 0.177651581 | 4.075003276 | 4.60E-05 | 0.006962251 |
| RPL10 | 84.09836356 | 2.266092578 | 0.211184176 | 10.73040899 | 7.33E-27 | 2.02E-23 |
| AC245140.2 | 3.095276149 | -0.637277714 | 0.137875421 | -4.622127049 | 3.80E-06 | 0.001035343 |
| FAM3A | 45.27826968 | -0.227090538 | 0.055779141 | -4.071244795 | 4.68E-05 | 0.006971287 |
| MT-TD | 21.34480412 | 1.162622741 | 0.210104085 | 5.533556101 | 3.14E-08 | 1.82E-05 |

| | | | | | | |
|---|---|---|---|---|---|---|
| MT-ND3 | 5984.612172 | 0.505785573 | 0.087655685 | 5.77013998 | 7.92E-09 | 5.00E-06 |
| MT-TL2 | 47.43955827 | 0.450346363 | 0.108493617 | 4.150901901 | 3.31E-05 | 0.005500621 |

**Table S1**: DE genes in DCIS from Black vs White women (n=384, P.adj < 0.05). Log2FC>0 = up in White vs Black DCIS.

#### Table S2

| | baseMean | log2FoldChang | lfcSE | stat | pvalue | padj |
|---|---|---|---|---|---|---|
| MAST1 | 36.68910536 | -3.222863418 | 0.511003661 | -6.306928233 | 2.85E-10 | 5.22E-06 |
| CIDEC | 17.17059804 | -1.60459299 | 0.301085349 | -5.329362568 | 9.86E-08 | 0.000822587 |
| NDRG1 | 265.3723272 | 0.984345708 | 0.188387161 | 5.225120997 | 1.74E-07 | 0.000822587 |
| RBM20 | 33.35338135 | -2.762469506 | 0.529257514 | -5.219518724 | 1.79E-07 | 0.000822587 |
| VEGFD | 8.056106499 | -1.740277292 | 0.34404649 | -5.058262023 | 4.23E-07 | 0.001552081 |
| CYP2A6 | 13.33461257 | -4.861471142 | 0.968043732 | -5.021954049 | 5.11E-07 | 0.001563607 |
| HOXA9 | 12.68169926 | 2.073388411 | 0.417896866 | 4.961483518 | 7.00E-07 | 0.001603935 |
| UBE3AP2 | 13.84893083 | -1.879316551 | 0.377821184 | -4.974089936 | 6.56E-07 | 0.001603935 |
| TAGLN2 | 644.9952062 | 0.651613272 | 0.133554895 | 4.878992046 | 1.07E-06 | 0.001774223 |
| HOXA10 | 21.69877265 | 1.308234966 | 0.268397602 | 4.874242372 | 1.09E-06 | 0.001774223 |
| PLIN1 | 61.93277069 | -1.324679847 | 0.272404197 | -4.862920112 | 1.16E-06 | 0.001774223 |
| SLPI | 104.8853045 | 1.678010436 | 0.345111944 | 4.862220702 | 1.16E-06 | 0.001774223 |
| INAVA | 2.248428386 | 1.983357729 | 0.420557639 | 4.716018792 | 2.41E-06 | 0.002205664 |
| MTHFD2 | 142.0041146 | 0.685698272 | 0.144657897 | 4.740137153 | 2.14E-06 | 0.002205664 |
| PON3 | 22.2117482 | 1.556522768 | 0.328169746 | 4.743041632 | 2.11E-06 | 0.002205664 |
| RBP4 | 31.03502263 | -1.566172715 | 0.331526361 | -4.724127241 | 2.31E-06 | 0.002205664 |
| GPAM | 69.44982032 | -0.888145729 | 0.185468517 | -4.788660327 | 1.68E-06 | 0.002205664 |
| TNS2 | 103.7216235 | -0.525989298 | 0.111376772 | -4.722612147 | 2.33E-06 | 0.002205664 |
| TRARG1 | 9.505776048 | -1.594735968 | 0.333391253 | -4.783376748 | 1.72E-06 | 0.002205664 |
| TK1 | 46.28673316 | 1.170590064 | 0.24681082 | 4.742863651 | 2.11E-06 | 0.002205664 |
| GLYATL2 | 44.57547019 | 2.714307568 | 0.581218515 | 4.670029423 | 3.01E-06 | 0.002630388 |
| PLIN4 | 79.03415536 | -1.290392731 | 0.278282323 | -4.63699138 | 3.54E-06 | 0.002947369 |
| CRISP3 | 58.45245141 | 2.994249622 | 0.664493157 | 4.506065396 | 6.60E-06 | 0.00507794 |
| CLDN5 | 57.03975796 | -0.874084942 | 0.194035225 | -4.504774547 | 6.64E-06 | 0.00507794 |
| FCRLB | 6.484691232 | -2.444946966 | 0.548929217 | -4.454029569 | 8.43E-06 | 0.006182977 |
| NCCRP1 | 14.5363019 | 2.063450802 | 0.464168251 | 4.445480271 | 8.77E-06 | 0.006186596 |
| PTX3 | 14.42638472 | -1.801760392 | 0.406094882 | -4.436796598 | 9.13E-06 | 0.006202823 |
| FMO4 | 12.94450045 | 1.161870313 | 0.262959325 | 4.418441193 | 9.94E-06 | 0.006310776 |
| ARPC4 | 115.6378157 | 0.402857633 | 0.091192657 | 4.417654307 | 9.98E-06 | 0.006310776 |
| UQCC1 | 32.75105566 | -0.476777646 | 0.108648714 | -4.38824932 | 1.14E-05 | 0.006986269 |
| NXPH4 | 12.60329992 | 1.853653338 | 0.425655973 | 4.354815755 | 1.33E-05 | 0.007879894 |
| TMEM175 | 49.11425492 | -0.493516965 | 0.114241087 | -4.319960348 | 1.56E-05 | 0.008875285 |

| | | | | | | |
|---|---|---|---|---|---|---|
| PELO | 21.92451398 | 0.553166202 | 0.128587168 | 4.301877162 | 1.69E-05 | 0.008875285 |
| HDC | 11.94613436 | -1.378985463 | 0.320220163 | -4.306366751 | 1.66E-05 | 0.008875285 |
| TNRC6A | 324.4448463 | -0.290575683 | 0.067491325 | -4.305378272 | 1.67E-05 | 0.008875285 |
| AQP7 | 6.99982718 | -1.397894524 | 0.32596667 | -4.288458463 | 1.80E-05 | 0.009166793 |
| RPN1 | 236.4679031 | 0.366785767 | 0.085790784 | 4.275351625 | 1.91E-05 | 0.009460301 |
| F11R | 128.6476752 | 0.496136995 | 0.117131439 | 4.235728682 | 2.28E-05 | 0.010213513 |
| GAPDH | 1368.456565 | 0.513841379 | 0.121028137 | 4.245635691 | 2.18E-05 | 0.010213513 |
| A2ML1 | 2.480194945 | 2.682723993 | 0.63257106 | 4.240984391 | 2.23E-05 | 0.010213513 |
| C18orf21 | 17.94651202 | 0.558357413 | 0.13183594 | 4.235244296 | 2.28E-05 | 0.010213513 |
| CYP4Z1 | 65.27517989 | 2.545057372 | 0.603625818 | 4.216283159 | 2.48E-05 | 0.010846318 |
| UCHL5 | 86.80512646 | 0.443283383 | 0.105672127 | 4.194894106 | 2.73E-05 | 0.011007651 |
| CAVIN2 | 74.0749235 | -0.868882898 | 0.207836012 | -4.180617646 | 2.91E-05 | 0.011007651 |
| ADIPOQ | 26.50022468 | -1.312164992 | 0.313983086 | -4.179094519 | 2.93E-05 | 0.011007651 |
| AP001528.2 | 13.73764565 | -0.875902721 | 0.209254777 | -4.185819474 | 2.84E-05 | 0.011007651 |
| GPD1 | 37.43076998 | -1.254549396 | 0.300274096 | -4.178014062 | 2.94E-05 | 0.011007651 |
| DCD | 30.94665922 | 2.915465933 | 0.696315875 | 4.1869876 | 2.83E-05 | 0.011007651 |
| ZBED4 | 39.79049361 | 0.471495772 | 0.112334681 | 4.197241396 | 2.70E-05 | 0.011007651 |
| CD46 | 451.9261611 | 0.439474616 | 0.10581126 | 4.153382301 | 3.28E-05 | 0.011959372 |
| LIFR | 109.9110737 | -0.520323547 | 0.125380322 | -4.149961793 | 3.33E-05 | 0.011959372 |
| S100A7 | 352.6858432 | 3.407731912 | 0.822845581 | 4.141399053 | 3.45E-05 | 0.012176047 |
| DBI | 487.5442697 | 0.721621155 | 0.17456192 | 4.133897895 | 3.57E-05 | 0.012288862 |
| FABP4 | 208.6127357 | -1.161720247 | 0.281407109 | -4.128254799 | 3.66E-05 | 0.012288862 |
| DHCR7 | 52.25987131 | 0.881533767 | 0.213632829 | 4.12639655 | 3.68E-05 | 0.012288862 |
| GPIHBP1 | 9.508654495 | -0.960908047 | 0.23341689 | -4.11670315 | 3.84E-05 | 0.012588199 |
| COL6A6 | 20.39736701 | -1.434780949 | 0.348963254 | -4.111553098 | 3.93E-05 | 0.012607984 |
| ASS1 | 53.32655051 | 0.988531719 | 0.240621592 | 4.108241962 | 3.99E-05 | 0.012607984 |
| ENO1 | 614.6314279 | 0.544584059 | 0.133019374 | 4.094020602 | 4.24E-05 | 0.012960354 |
| LPL | 106.6390328 | -0.953318366 | 0.232745799 | -4.095963802 | 4.20E-05 | 0.012960354 |
| GNG11 | 98.41198128 | -0.522251857 | 0.127894043 | -4.083472891 | 4.44E-05 | 0.01334083 |
| S100A11 | 601.4910788 | 0.552798018 | 0.136428576 | 4.051922503 | 5.08E-05 | 0.014134818 |
| NDUFS2 | 119.0242407 | 0.400905268 | 0.098814336 | 4.057156929 | 4.97E-05 | 0.014134818 |
| LINC02511 | 4.90272182 | -1.378241919 | 0.339486655 | -4.059782313 | 4.91E-05 | 0.014134818 |
| GGCT | 123.3287869 | 0.698307662 | 0.172352132 | 4.05163345 | 5.09E-05 | 0.014134818 |
| AOC3 | 27.92433325 | -0.718543804 | 0.176909213 | -4.061652822 | 4.87E-05 | 0.014134818 |
| PDGFD | 48.38175524 | -0.796832529 | 0.197017784 | -4.044470066 | 5.24E-05 | 0.014356477 |

| | | | | | | |
|---|---|---|---|---|---|---|
| COP1 | 36.34936352 | 0.436047205 | 0.108224274 | 4.029107233 | 5.60E-05 | 0.01510223 |
| PERP | 301.7329588 | 0.676812979 | 0.168543147 | 4.015665977 | 5.93E-05 | 0.015313804 |
| LRRN4CL | 13.37597685 | -1.069753487 | 0.266247153 | -4.01789644 | 5.87E-05 | 0.015313804 |
| NNAT | 4.157163517 | -1.621729024 | 0.403819593 | -4.015974096 | 5.92E-05 | 0.015313804 |
| AC010136.1 | 2.207104973 | -1.474238302 | 0.368345501 | -4.00232471 | 6.27E-05 | 0.015675257 |
| ABRACL | 68.439464 | 0.85465051 | 0.21364239 | 4.00037891 | 6.32E-05 | 0.015675257 |
| TOX | 8.181384314 | 1.267048771 | 0.316371253 | 4.004942796 | 6.20E-05 | 0.015675257 |
| SRD5A3 | 53.47585895 | 0.809280522 | 0.202673295 | 3.993029874 | 6.52E-05 | 0.015953712 |
| NOA1 | 25.2105018 | 0.445276831 | 0.111793688 | 3.983023016 | 6.80E-05 | 0.016240728 |
| TNNT3 | 9.450005359 | -1.051699105 | 0.264076593 | -3.9825533 | 6.82E-05 | 0.016240728 |
| G3BP2 | 202.0571899 | 0.261325819 | 0.065872714 | 3.96713301 | 7.27E-05 | 0.016486568 |
| GPM6A | 3.492516212 | -1.414048325 | 0.355969648 | -3.972384532 | 7.12E-05 | 0.016486568 |
| WSCD2 | 1.658464888 | -1.925674006 | 0.484513112 | -3.974451796 | 7.05E-05 | 0.016486568 |
| CHCHD10 | 155.040826 | 0.562194216 | 0.141720445 | 3.966923862 | 7.28E-05 | 0.016486568 |
| HDGF | 526.4731661 | 0.36259981 | 0.091633019 | 3.957086792 | 7.59E-05 | 0.016970683 |
| RUSC1-AS1 | 3.437490405 | -1.183385788 | 0.300307981 | -3.940573885 | 8.13E-05 | 0.017749577 |
| ABCA6 | 74.7826075 | -0.830765453 | 0.21067219 | -3.943403507 | 8.03E-05 | 0.017749577 |
| CSTB | 191.5321009 | 0.640520294 | 0.162936484 | 3.93110419 | 8.46E-05 | 0.018246323 |
| CAV1 | 222.33926 | -0.536911328 | 0.137195142 | -3.913486451 | 9.10E-05 | 0.01917962 |
| SNX21 | 35.40183461 | -0.42542403 | 0.108670887 | -3.914793009 | 9.05E-05 | 0.01917962 |
| AKR1C3 | 32.55799499 | -0.663698789 | 0.170021157 | -3.903624714 | 9.48E-05 | 0.019751559 |
| PFKFB2 | 24.34886829 | 0.710898794 | 0.182954588 | 3.885657096 | 0.000102053 | 0.020798498 |
| GLB1L3 | 1.957861603 | 1.865495585 | 0.479901592 | 3.887246089 | 0.000101388 | 0.020798498 |
| TIPRL | 90.94896154 | 0.333214396 | 0.086360936 | 3.858392582 | 0.000114135 | 0.022204615 |
| C1orf116 | 20.65549459 | 1.048622366 | 0.271720622 | 3.859193169 | 0.000113762 | 0.022204615 |
| LUC7L3 | 595.1516853 | -0.462310173 | 0.119607037 | -3.865242257 | 0.000110979 | 0.022204615 |
| FIRRE | 13.71263192 | 1.156740866 | 0.299943094 | 3.85653442 | 0.000115006 | 0.022204615 |
| NAA10 | 64.01574736 | 0.362594531 | 0.094020539 | 3.856545972 | 0.000115 | 0.022204615 |
| AL079343.1 | 5.81604385 | -1.786199965 | 0.463790466 | -3.851308071 | 0.000117489 | 0.022447664 |
| CD302 | 64.44185417 | -0.458741722 | 0.119352951 | -3.843572502 | 0.000121256 | 0.022928664 |
| ABCA9 | 69.37261283 | -0.892149139 | 0.232440239 | -3.838187153 | 0.000123946 | 0.023198138 |
| SFT2D2 | 118.9752083 | 0.49486668 | 0.129417404 | 3.823803187 | 0.000131409 | 0.023857644 |
| SAPCD2 | 29.48665472 | 0.823299213 | 0.21503745 | 3.828631778 | 0.000128858 | 0.023857644 |
| CIDEA | 5.738381978 | -1.525944954 | 0.399311085 | -3.821444007 | 0.000132673 | 0.023857644 |
| TDRD12 | 4.334269336 | -2.269382091 | 0.593522395 | -3.823582919 | 0.000131526 | 0.023857644 |

| | | | | | | |
|---|---|---|---|---|---|---|
| SYT12 | 57.5940028 | 0.970869577 | 0.254485417 | 3.8150303 | 0.000136166 | 0.024248164 |
| S100A9 | 472.517984 | 1.712187761 | 0.450249809 | 3.802750668 | 0.000143098 | 0.024761416 |
| ABCA10 | 56.87835414 | -0.900788724 | 0.236621111 | -3.806882327 | 0.00014073 | 0.024761416 |
| MAGT1 | 139.4869432 | 0.337235805 | 0.088657888 | 3.803787914 | 0.0001425 | 0.024761416 |
| HSPB7 | 10.67970414 | -1.107839589 | 0.291859871 | -3.795792776 | 0.000147172 | 0.024977937 |
| AC069120.1 | 9.481746617 | -1.232005029 | 0.324828671 | -3.792784134 | 0.000148968 | 0.024977937 |
| CCDC6 | 159.3367135 | 0.393583945 | 0.103726831 | 3.794427542 | 0.000147984 | 0.024977937 |
| PABPC1L | 46.20043803 | -0.709306493 | 0.187082703 | -3.79140606 | 0.000149797 | 0.024977937 |
| TMEM51 | 37.63621674 | 0.525467584 | 0.138932413 | 3.78218137 | 0.00015546 | 0.02538167 |
| GMDS | 22.18092403 | 0.72587957 | 0.192106052 | 3.778535665 | 0.000157753 | 0.02538167 |
| KCNC2 | 71.98687304 | -2.369691895 | 0.626930688 | -3.779830751 | 0.000156935 | 0.02538167 |
| CD300LG | 13.42695729 | -0.905780102 | 0.239486225 | -3.782180379 | 0.000155461 | 0.02538167 |
| TES | 132.4537146 | 0.425329307 | 0.112771017 | 3.771618956 | 0.000162192 | 0.025789834 |
| DPAGT1 | 48.98209704 | 0.426903241 | 0.113230258 | 3.770222283 | 0.000163102 | 0.025789834 |
| PDXK | 109.2267217 | 0.361567861 | 0.096107995 | 3.762099706 | 0.000168493 | 0.026414494 |
| LSR | 153.6603196 | 0.49740875 | 0.132693134 | 3.748564349 | 0.00017785 | 0.027645079 |
| PFDN2 | 86.41592429 | 0.463472608 | 0.12373419 | 3.745711728 | 0.000179883 | 0.027726177 |
| MEOX2 | 14.90430328 | -0.85935578 | 0.229598567 | -3.742862129 | 0.000181936 | 0.027808927 |
| NFE2L3 | 9.915584237 | 1.050244833 | 0.281973217 | 3.724626208 | 0.000195605 | 0.029651107 |
| ZNF483 | 32.42193688 | -0.72327601 | 0.194389258 | -3.7207612 | 0.000198623 | 0.029861857 |
| NECTIN4 | 98.4240505 | 0.599153381 | 0.161477622 | 3.710442194 | 0.000206898 | 0.030852964 |
| CD82 | 101.0589656 | 0.697135156 | 0.188252956 | 3.70318306 | 0.000212911 | 0.031493677 |
| OXSM | 7.945832696 | 0.65772738 | 0.177840372 | 3.698414336 | 0.000216951 | 0.031834455 |
| STX6 | 60.95605891 | 0.417090281 | 0.113320053 | 3.680639653 | 0.00023265 | 0.032144136 |
| FANCD2 | 23.42311418 | 0.523594397 | 0.142297882 | 3.679565643 | 0.000233632 | 0.032144136 |
| FRK | 30.33004691 | 0.814783112 | 0.221728728 | 3.674684464 | 0.000238144 | 0.032144136 |
| PABPC1 | 2742.592388 | 0.499265284 | 0.13573025 | 3.678364136 | 0.000234735 | 0.032144136 |
| LYVE1 | 44.50160275 | -0.900142454 | 0.244315251 | -3.684348196 | 0.000229289 | 0.032144136 |
| DNAH9 | 2.31824064 | -2.02427017 | 0.550882556 | -3.674594786 | 0.000238227 | 0.032144136 |
| EZH1 | 71.81472286 | -0.399673555 | 0.108674341 | -3.67771778 | 0.00023533 | 0.032144136 |
| LINC00663 | 5.43074539 | -0.798934968 | 0.216727754 | -3.686352815 | 0.000227491 | 0.032144136 |
| LIPE | 25.59385607 | -0.842336054 | 0.229239798 | -3.674475648 | 0.000238338 | 0.032144136 |
| HSPA12B | 19.77724726 | -0.730926261 | 0.19817059 | -3.688369003 | 0.000225696 | 0.032144136 |
| BMX | 6.405175943 | -1.074039625 | 0.291078047 | -3.689868186 | 0.00022437 | 0.032144136 |
| GALNT14 | 8.101465716 | 1.323781519 | 0.360886907 | 3.668133957 | 0.000244327 | 0.032711306 |

| | | | | | | |
|---|---|---|---|---|---|---|
| HOXA11 | 2.227087195 | 2.143307741 | 0.584800162 | 3.665025898 | 0.000247314 | 0.032871197 |
| SDHC | 85.2050184 | 0.336198997 | 0.092115241 | 3.649765175 | 0.00026248 | 0.033012212 |
| TP53BP2 | 49.73508761 | 0.613781294 | 0.168033936 | 3.652722232 | 0.000259475 | 0.033012212 |
| PPP1CB | 266.0451604 | 0.360339631 | 0.098633726 | 3.653310546 | 0.000258881 | 0.033012212 |
| BTNL9 | 58.92044143 | -0.776520912 | 0.212267938 | -3.658211026 | 0.000253982 | 0.033012212 |
| WDR86 | 12.7251577 | -0.821495779 | 0.224558054 | -3.65827796 | 0.000253916 | 0.033012212 |
| TXN | 325.2339618 | 0.368753439 | 0.101042771 | 3.649478692 | 0.000262773 | 0.033012212 |
| CES3 | 5.062300305 | 2.021267537 | 0.552492238 | 3.658454178 | 0.000253741 | 0.033012212 |
| LAMP2 | 249.6878274 | 0.312452166 | 0.085487725 | 3.654936052 | 0.000257246 | 0.033012212 |
| FAR2 | 14.00590133 | 1.065155019 | 0.292107762 | 3.646445451 | 0.000265893 | 0.033176921 |
| TOMM40L | 21.33010085 | 0.510996622 | 0.140463849 | 3.637922685 | 0.000274846 | 0.034062321 |
| CHRD | 27.50143394 | -0.776649871 | 0.213674359 | -3.634735925 | 0.000278266 | 0.034254672 |
| FH | 40.64407656 | 0.554423475 | 0.152625939 | 3.632563885 | 0.000280619 | 0.0343141 |
| TOR1AIP2 | 185.4526905 | 0.269523301 | 0.074374189 | 3.623882241 | 0.000290214 | 0.034580867 |
| KCMF1 | 89.63642514 | 0.263080364 | 0.072525231 | 3.627432299 | 0.000286254 | 0.034580867 |
| LEP | 24.61347621 | -1.272736561 | 0.35115692 | -3.624409735 | 0.000289622 | 0.034580867 |
| FHL1 | 96.40887157 | -0.778075952 | 0.214714613 | -3.62376803 | 0.000290342 | 0.034580867 |
| HMGA1 | 144.4964303 | 0.554955522 | 0.153230366 | 3.621707212 | 0.000292665 | 0.034632684 |
| GALNT15 | 19.30609827 | -0.790016894 | 0.218474593 | -3.616058431 | 0.000299123 | 0.035169948 |
| CD55 | 83.29630672 | 0.587597159 | 0.162766355 | 3.610065225 | 0.00030612 | 0.035763397 |
| BROX | 80.8264897 | 0.345669735 | 0.0958407 | 3.60671128 | 0.000310102 | 0.035782797 |
| CCN5 | 96.81004305 | -0.878451808 | 0.243565164 | -3.606639773 | 0.000310188 | 0.035782797 |
| ITIH5 | 83.61721826 | -0.678135387 | 0.18811081 | -3.604978288 | 0.000312179 | 0.035787423 |
| E2F2 | 4.818084678 | 1.079475849 | 0.29980603 | 3.60058084 | 0.000317507 | 0.035948858 |
| ZFHX4 | 101.6000453 | -0.70607567 | 0.196023684 | -3.601991629 | 0.000315789 | 0.035948858 |
| YWHAQ | 58.25048669 | 0.364209215 | 0.101204748 | 3.598736452 | 0.000319767 | 0.03598262 |
| SLC19A3 | 10.87570867 | -1.001437008 | 0.279048459 | -3.588756632 | 0.000332259 | 0.036935099 |
| PPP1R12C | 106.1878147 | -0.316673786 | 0.088206735 | -3.590131598 | 0.000330511 | 0.036935099 |
| RGN | 5.394727053 | -1.116317641 | 0.31126344 | -3.586407841 | 0.000335264 | 0.037044705 |
| LAD1 | 59.25202892 | 0.954958782 | 0.266805927 | 3.579226272 | 0.000344613 | 0.03784965 |
| HAP1 | 2.48560172 | -1.203312636 | 0.336549004 | -3.575445543 | 0.000349632 | 0.038172317 |
| S100A8 | 203.8197456 | 1.801689677 | 0.504934372 | 3.568166035 | 0.000359489 | 0.03874207 |
| SLC39A7 | 244.7182085 | 0.313872467 | 0.087921894 | 3.56990112 | 0.000357116 | 0.03874207 |
| NDUFA8 | 56.60410068 | 0.397496173 | 0.111439279 | 3.566930583 | 0.000361187 | 0.03874207 |
| LEPR | 80.96927327 | -0.745762895 | 0.209637344 | -3.557395258 | 0.00037455 | 0.039392684 |

| | | | | | | |
|---|---|---|---|---|---|---|
| ABHD10 | 48.00636146 | 0.390089614 | 0.10955955 | 3.560525883 | 0.000370113 | 0.039392684 |
| ADH1B | 113.1912145 | -0.896970578 | 0.252206723 | -3.556489562 | 0.000375843 | 0.039392684 |
| LHFPL6 | 89.38131055 | -0.566129751 | 0.159114907 | -3.557993165 | 0.000373699 | 0.039392684 |
| COPA | 295.8031846 | 0.285429723 | 0.080436692 | 3.548501498 | 0.00038743 | 0.040376348 |
| TGFBR3 | 153.6944185 | -0.701379526 | 0.198505936 | -3.53329245 | 0.000410418 | 0.041644788 |
| PSMA5 | 92.69157593 | 0.331917593 | 0.093949188 | 3.532947954 | 0.000410953 | 0.041644788 |
| FGFR1 | 340.7191317 | -0.566328873 | 0.160258069 | -3.533855586 | 0.000409545 | 0.041644788 |
| PENK | 9.133139732 | -1.660409019 | 0.469384586 | -3.537417011 | 0.000404061 | 0.041644788 |
| DCX | 13.36432176 | -1.365698729 | 0.386465826 | -3.533814989 | 0.000409608 | 0.041644788 |
| LRRC8B | 38.96864858 | 0.460034927 | 0.130525127 | 3.524493225 | 0.000424294 | 0.042066997 |
| PRSS2 | 2.832963317 | -4.237192331 | 1.201959268 | -3.52523787 | 0.000423103 | 0.042066997 |
| MAP3K12 | 29.01179337 | -0.463053546 | 0.131279634 | -3.527230614 | 0.000419931 | 0.042066997 |
| ZBTB4 | 204.4409073 | -0.285165263 | 0.080904102 | -3.524731806 | 0.000423912 | 0.042066997 |
| STK38L | 48.21367985 | 0.419459546 | 0.119141465 | 3.520684825 | 0.000430434 | 0.042446337 |
| CALCOCO1 | 72.92253155 | -0.363788514 | 0.10347969 | -3.515554741 | 0.000438836 | 0.043043503 |
| BSPRY | 59.09164822 | 0.506898048 | 0.144298138 | 3.51285232 | 0.000443324 | 0.043252369 |
| ITPR3 | 46.27825849 | 0.399058975 | 0.113776695 | 3.50738765 | 0.000452529 | 0.043883628 |
| NANOS1 | 10.47989595 | 1.077005475 | 0.307173028 | 3.506185044 | 0.000454579 | 0.043883628 |
| PLPP3 | 231.472246 | -0.371112857 | 0.106042312 | -3.499667722 | 0.000465838 | 0.044006941 |
| F5 | 4.2490383 | 1.420215697 | 0.405573683 | 3.501745195 | 0.000462221 | 0.044006941 |
| FAM228B | 26.63420439 | -0.428880223 | 0.122471165 | -3.501887344 | 0.000461975 | 0.044006941 |
| FREM1 | 6.628525376 | -0.936046372 | 0.267555161 | -3.498517342 | 0.000467853 | 0.044006941 |
| PCED1A | 78.00210481 | -0.396067957 | 0.113134361 | -3.500863517 | 0.000463753 | 0.044006941 |
| GABBR2 | 2.02382744 | 1.861983635 | 0.53254775 | 3.496369356 | 0.000471635 | 0.044136409 |
| PCDH18 | 48.75485775 | -0.622795313 | 0.178361417 | -3.491760295 | 0.000479849 | 0.044677077 |
| SDC1 | 253.8374092 | 0.834224548 | 0.239104864 | 3.488948469 | 0.000484925 | 0.044921654 |
| CHIT1 | 36.2309437 | -1.580141036 | 0.453696483 | -3.482815263 | 0.000496171 | 0.045041374 |
| UGT2B4 | 4.910906688 | -4.98458964 | 1.432981009 | -3.478475714 | 0.000504274 | 0.045041374 |
| KIF2A | 64.28704016 | 0.319994783 | 0.091879033 | 3.482783544 | 0.000496229 | 0.045041374 |
| ERMARD | 22.00131155 | -0.45853853 | 0.131840763 | -3.477972362 | 0.000505222 | 0.045041374 |
| AUXG01000058.1 | 26.94930857 | -0.681052415 | 0.195704355 | -3.480006445 | 0.000501402 | 0.045041374 |
| ADAM8 | 14.88278887 | 0.742921991 | 0.213628638 | 3.477632945 | 0.000505862 | 0.045041374 |
| SGK2 | 5.606524642 | -1.135560517 | 0.326430658 | -3.478718954 | 0.000503817 | 0.045041374 |
| PLXNB2 | 187.970412 | 0.351447583 | 0.100801894 | 3.486517658 | 0.000489353 | 0.045041374 |
| SEPTIN3 | 7.453326326 | 1.13434564 | 0.326623401 | 3.472946629 | 0.000514778 | 0.045613782 |

| | | | | | | |
|---|---|---|---|---|---|---|
| IGFBP6 | 104.8102928 | -0.671901306 | 0.193691512 | -3.468924894 | 0.000522546 | 0.04607947 |
| PRELID3A | 8.006765849 | -0.786987663 | 0.22701951 | -3.46660806 | 0.00052707 | 0.046256051 |
| CYBRD1 | 187.6592022 | -0.677312153 | 0.195503881 | -3.464443513 | 0.00053133 | 0.046407855 |
| MDH1 | 154.0087381 | 0.322024491 | 0.093134779 | 3.457618037 | 0.000544973 | 0.047373948 |
| H4C3 | 216.9745028 | 0.423501689 | 0.12264568 | 3.453050177 | 0.000554286 | 0.047507999 |
| SOBP | 35.44990693 | -0.577708459 | 0.167276033 | -3.453623626 | 0.000553109 | 0.047507999 |
| CMA1 | 7.187089197 | -1.038653995 | 0.300648033 | -3.45471742 | 0.00055087 | 0.047507999 |
| CADM3-AS1 | 7.898819897 | -0.795502426 | 0.230516812 | -3.450951886 | 0.000558613 | 0.047656197 |
| LPAR1 | 36.42332188 | -0.609391849 | 0.176662088 | -3.449477226 | 0.000561673 | 0.047695408 |
| EXTL2 | 48.91833144 | -0.330556617 | 0.095875388 | -3.447773451 | 0.000565228 | 0.047776085 |
| CXCL12 | 261.6298318 | -0.560111448 | 0.162518207 | -3.446453536 | 0.000567996 | 0.047789848 |
| PDZK1IP1 | 66.20156332 | 1.23664228 | 0.362249474 | 3.41378627 | 0.000640668 | 0.048369838 |
| ILF2 | 134.0086858 | 0.344474967 | 0.101023524 | 3.409849054 | 0.000649988 | 0.048369838 |
| DAP3 | 117.1149903 | 0.344997978 | 0.100669825 | 3.427024697 | 0.000610234 | 0.048369838 |
| DHX9 | 141.9738121 | 0.212306315 | 0.061895808 | 3.430059677 | 0.000603449 | 0.048369838 |
| APOB | 2.181756144 | -1.781909586 | 0.519000946 | -3.433345547 | 0.000596182 | 0.048369838 |
| CHMP3 | 230.743923 | 0.219404356 | 0.064104624 | 3.422597982 | 0.000620257 | 0.048369838 |
| KCNE4 | 99.62472294 | -1.213782317 | 0.355500151 | -3.414294796 | 0.000639473 | 0.048369838 |
| MST1 | 18.41216627 | -0.763504358 | 0.223486176 | -3.416338196 | 0.000634694 | 0.048369838 |
| DAPP1 | 33.40679257 | 0.72041017 | 0.210151165 | 3.428056984 | 0.000607918 | 0.048369838 |
| ADAMTS19 | 7.448842414 | -1.442321972 | 0.42303758 | -3.409441717 | 0.00065096 | 0.048369838 |
| ADTRP | 5.546158139 | 1.110223368 | 0.325648252 | 3.409271698 | 0.000651366 | 0.048369838 |
| OGN | 110.9856955 | -0.853159472 | 0.249134215 | -3.424497397 | 0.000615938 | 0.048369838 |
| BBOX1 | 9.154507816 | 1.321956203 | 0.385671652 | 3.427672729 | 0.000608779 | 0.048369838 |
| SOX5 | 17.37955368 | -0.744321346 | 0.217790876 | -3.417596558 | 0.000631767 | 0.048369838 |
| RASSF8 | 64.6866264 | -0.517889316 | 0.150859327 | -3.432928718 | 0.000597099 | 0.048369838 |
| DCN | 1460.710012 | -0.681309207 | 0.199633726 | -3.412796133 | 0.000643 | 0.048369838 |
| GALNT16 | 30.27175391 | -0.691565168 | 0.202214215 | -3.419963175 | 0.000626296 | 0.048369838 |
| KIF26A | 6.986366628 | -0.751261903 | 0.220255593 | -3.410864139 | 0.000647574 | 0.048369838 |
| BUB1B | 8.889568339 | 0.826143138 | 0.241040301 | 3.427406683 | 0.000609376 | 0.048369838 |
| GPT2 | 29.28634777 | 0.737151272 | 0.214805946 | 3.431707952 | 0.000599793 | 0.048369838 |
| SLC7A5 | 44.89239187 | 1.079768039 | 0.314973258 | 3.428126078 | 0.000607763 | 0.048369838 |
| HIC1 | 44.15173104 | -0.575074572 | 0.167594655 | -3.431341968 | 0.000600603 | 0.048369838 |
| GID4 | 19.30008229 | -0.386435445 | 0.113092126 | -3.416996918 | 0.00063316 | 0.048369838 |
| KAT2A | 85.38920403 | -0.429479868 | 0.125515574 | -3.421725721 | 0.000622251 | 0.048369838 |

| | | | | | | |
|---|---|---|---|---|---|---|
| LINC00511 | 8.937613038 | 0.889898709 | 0.260484416 | 3.416322264 | 0.000634731 | 0.048369838 |
| TXNL1 | 126.5142395 | 0.23667627 | 0.069044566 | 3.427876859 | 0.000608321 | 0.048369838 |
| MIRLET7BHG | 25.11696972 | -0.69435439 | 0.201759847 | -3.441489469 | 0.000578521 | 0.048369838 |
| U62317.1 | 2.993954799 | 1.752896077 | 0.513943838 | 3.41067632 | 0.00064802 | 0.048369838 |
| PGK1 | 280.2145149 | 0.392477937 | 0.114863166 | 3.41691729 | 0.000633345 | 0.048369838 |
| HAND2 | 4.594425584 | -1.074162352 | 0.315303528 | -3.40675653 | 0.000657397 | 0.0486209 |
| AC020929.1 | 4.458362844 | -1.129074513 | 0.331530265 | -3.405645378 | 0.000660079 | 0.048623139 |
| UFC1 | 182.417181 | 0.330764848 | 0.097344221 | 3.397888889 | 0.00067908 | 0.048657532 |
| LINC02202 | 2.616332692 | -1.321183063 | 0.389191534 | -3.394686033 | 0.000687073 | 0.048657532 |
| SDK1 | 19.12373523 | -0.722237812 | 0.212713211 | -3.395359449 | 0.000685386 | 0.048657532 |
| ARPC5L | 109.352103 | 0.338958839 | 0.099726198 | 3.398894645 | 0.000676588 | 0.048657532 |
| C9orf16 | 152.6859015 | 0.307652413 | 0.090458567 | 3.401031231 | 0.000671322 | 0.048657532 |
| TMEM88 | 12.71802497 | -0.581471484 | 0.171270424 | -3.39504901 | 0.000686163 | 0.048657532 |
| CBX2 | 8.625129468 | 1.14445999 | 0.33652174 | 3.400850085 | 0.000671767 | 0.048657532 |
| SPC24 | 9.249096538 | 0.928853339 | 0.273266172 | 3.399079113 | 0.000676132 | 0.048657532 |
| COX6B1 | 329.7016215 | 0.325082172 | 0.095674855 | 3.39778065 | 0.000679349 | 0.048657532 |
| PLP1 | 8.187914698 | -1.027971508 | 0.302167007 | -3.401997854 | 0.000668952 | 0.048657532 |
| MT-CO2 | 725.8715982 | 0.840358703 | 0.24773601 | 3.392154026 | 0.000693454 | 0.048920545 |
| EMCN | 67.02094367 | -0.472951433 | 0.139500282 | -3.390325996 | 0.000698096 | 0.049059267 |
| CYCS | 132.4357969 | 0.325419847 | 0.096045492 | 3.388184479 | 0.000703569 | 0.049094763 |
| SRSF6 | 207.9114281 | -0.288320715 | 0.085099697 | -3.388034572 | 0.000703954 | 0.049094763 |
| LAMC3 | 9.988321458 | -1.182542401 | 0.349432417 | -3.384180585 | 0.00071391 | 0.049600542 |
| SHMT2 | 60.84716106 | 0.472165184 | 0.139626777 | 3.38162344 | 0.000720588 | 0.049836041 |
| ADCY4 | 30.70233036 | -0.529442286 | 0.156602359 | -3.380806578 | 0.000722734 | 0.049836041 |

**Table S2:** DE genes in Black cases vs controls (n=266, P.adj < 0.05). Log2FC>0 = up in cases vs controls.

### Table S3

| | baseMean | log2FoldChang | lfcSE | stat | pvalue | padj |
|---|---|---|---|---|---|---|
| GLRA3 | 63.97937577 | -3.600043344 | 0.533280314 | -6.750752366 | 1.47E-11 | 2.72E-07 |
| RN7SL752P | 11.52318523 | -1.095309592 | 0.187612649 | -5.838143634 | 5.28E-09 | 2.37E-05 |
| RN7SL8P | 11.3826316 | -1.141666587 | 0.196656872 | -5.805373456 | 6.42E-09 | 2.37E-05 |
| SDCBPP2 | 12.73807833 | -1.914326676 | 0.329526263 | -5.809329608 | 6.27E-09 | 2.37E-05 |
| TOGARAM1 | 72.70723101 | -0.419188939 | 0.071745992 | -5.842680961 | 5.14E-09 | 2.37E-05 |
| RPL19 | 1690.519316 | 1.074282917 | 0.188703675 | 5.692962353 | 1.25E-08 | 3.84E-05 |
| SNORA13 | 51.66099512 | -0.834020532 | 0.147305273 | -5.661851161 | 1.50E-08 | 3.95E-05 |
| RN7SL838P | 18.68426667 | -1.064487238 | 0.192453039 | -5.531153183 | 3.18E-08 | 7.34E-05 |
| SNORA75 | 86.525957 | -0.992743836 | 0.180246377 | -5.507704796 | 3.64E-08 | 7.46E-05 |
| S100A9 | 529.1532596 | 1.947710349 | 0.35653308 | 5.462916232 | 4.68E-08 | 8.65E-05 |
| RPS7 | 64.21385194 | 0.9302486 | 0.171792953 | 5.414940386 | 6.13E-08 | 9.44E-05 |
| LTF | 289.3285827 | 1.673150047 | 0.308506466 | 5.4233873 | 5.85E-08 | 9.44E-05 |
| PTMA | 433.3086352 | 0.680229157 | 0.12745635 | 5.336957755 | 9.45E-08 | 0.000134282 |
| PKD1L3 | 6.706831527 | -0.762428086 | 0.146248774 | -5.213227209 | 1.86E-07 | 0.000228503 |
| RPS19 | 429.2109337 | 0.681320508 | 0.130562754 | 5.218337456 | 1.81E-07 | 0.000228503 |
| RPS27 | 372.0443796 | 0.881362388 | 0.171393147 | 5.142343208 | 2.71E-07 | 0.000275154 |
| SCN7A | 17.99808201 | -0.936032225 | 0.182306525 | -5.13438686 | 2.83E-07 | 0.000275154 |
| SNORA20 | 20.25989871 | -0.765850301 | 0.149112054 | -5.136072375 | 2.81E-07 | 0.000275154 |
| RPL23A | 222.3380893 | 1.022636082 | 0.198889939 | 5.141718523 | 2.72E-07 | 0.000275154 |
| MT-CO2 | 628.9122017 | 0.963176087 | 0.188607453 | 5.106776388 | 3.28E-07 | 0.000302616 |
| FRS2 | 145.4040368 | -0.82659196 | 0.162728019 | -5.0795921 | 3.78E-07 | 0.000332658 |
| YBX1 | 87.2562182 | 0.680922133 | 0.135040035 | 5.042372323 | 4.60E-07 | 0.000339679 |
| RPS2 | 286.0018282 | 0.563463584 | 0.111346514 | 5.060451052 | 4.18E-07 | 0.000339679 |
| RPL23 | 1731.74663 | 0.916055481 | 0.181560832 | 5.045446587 | 4.52E-07 | 0.000339679 |
| TMEM47 | 79.85215254 | -0.901660896 | 0.178796333 | -5.042949589 | 4.58E-07 | 0.000339679 |
| SNORA35B | 16.1919445 | -0.987064385 | 0.196908759 | -5.012800797 | 5.36E-07 | 0.000381054 |
| RN7SL277P | 2.804033025 | -1.262315929 | 0.253653482 | -4.976536973 | 6.47E-07 | 0.000407266 |
| RPS27A | 382.2518833 | 0.67244691 | 0.135158628 | 4.975242196 | 6.52E-07 | 0.000407266 |
| TFPI2 | 64.89411478 | -1.850665051 | 0.373132682 | -4.959804214 | 7.06E-07 | 0.000407266 |
| CISD3 | 201.4447578 | 0.992056712 | 0.199929642 | 4.962029151 | 6.98E-07 | 0.000407266 |
| AC009283.1 | 25.2767834 | 1.400789426 | 0.281886454 | 4.969339272 | 6.72E-07 | 0.000407266 |

| | | | | | | |
|---|---|---|---|---|---|---|
| TOMM40 | 42.82562775 | 0.420543003 | 0.084302262 | 4.988513833 | 6.08E-07 | 0.000407266 |
| RN7SKP104 | 18.25845386 | -0.866076439 | 0.175150785 | -4.944747692 | 7.62E-07 | 0.000423374 |
| SRCIN1 | 28.25935098 | 1.356518966 | 0.274573596 | 4.940456721 | 7.79E-07 | 0.000423374 |
| SLC25A36 | 227.0521175 | -0.324969627 | 0.066474395 | -4.888643646 | 1.02E-06 | 0.000535776 |
| GRB14 | 32.87111075 | -1.925859208 | 0.395147821 | -4.873769018 | 1.09E-06 | 0.000561709 |
| PIN4 | 39.10972033 | 0.436854782 | 0.090024036 | 4.852646046 | 1.22E-06 | 0.000608104 |
| VDAC1 | 71.16061725 | 0.490696664 | 0.101368189 | 4.840736204 | 1.29E-06 | 0.000628719 |
| SET | 197.7547621 | 0.363314961 | 0.075292485 | 4.825381471 | 1.40E-06 | 0.000661737 |
| ATP5MG | 173.8539005 | 0.376045042 | 0.07801948 | 4.819886532 | 1.44E-06 | 0.000663221 |
| RPL26 | 525.4623526 | 0.583195442 | 0.121150807 | 4.813797434 | 1.48E-06 | 0.000667086 |
| ALB | 11.52792642 | -2.183025928 | 0.454474025 | -4.803411872 | 1.56E-06 | 0.000685924 |
| NDUFS5 | 130.1282796 | 0.406831791 | 0.084868291 | 4.79368425 | 1.64E-06 | 0.000693586 |
| MTCO1P40 | 72.194575 | 0.765567588 | 0.159764003 | 4.791865322 | 1.65E-06 | 0.000693586 |
| RPL9 | 279.5658294 | 0.509962914 | 0.107312921 | 4.752111016 | 2.01E-06 | 0.000826195 |
| RN7SL151P | 16.61857783 | -0.833198351 | 0.175512061 | -4.747242694 | 2.06E-06 | 0.000827927 |
| RPS15 | 118.1599841 | 0.817520687 | 0.172634084 | 4.735569411 | 2.18E-06 | 0.00085838 |
| RNU2-64P | 24.90382131 | 0.996783908 | 0.210756108 | 4.729561194 | 2.25E-06 | 0.000865756 |
| KRR1 | 180.6096453 | -0.284809308 | 0.060312862 | -4.722198542 | 2.33E-06 | 0.000879384 |
| LINC02611 | 17.96447543 | 1.454912851 | 0.308692096 | 4.713152266 | 2.44E-06 | 0.000900969 |
| MRPS21 | 125.95129 | 0.442903328 | 0.094136972 | 4.704881824 | 2.54E-06 | 0.000907082 |
| IRAK1 | 136.9951856 | 0.306074116 | 0.065069855 | 4.703777467 | 2.55E-06 | 0.000907082 |
| GDAP1 | 46.37885257 | -1.172049075 | 0.249750627 | -4.692877404 | 2.69E-06 | 0.000923731 |
| PSMB6 | 86.84834931 | 0.363418926 | 0.077449208 | 4.692351762 | 2.70E-06 | 0.000923731 |
| AC112128.1 | 3.500274008 | -1.042214148 | 0.223065968 | -4.672223905 | 2.98E-06 | 0.001000536 |
| ATF2 | 97.65329226 | -0.243982444 | 0.052407043 | -4.655527793 | 3.23E-06 | 0.001005776 |
| RNU1-89P | 3.158592106 | -1.05845403 | 0.227111859 | -4.660496521 | 3.15E-06 | 0.001005776 |
| RPS10 | 166.3787942 | 0.418084297 | 0.089847821 | 4.653249131 | 3.27E-06 | 0.001005776 |
| EIF3K | 207.7518305 | 0.3058401 | 0.065694862 | 4.655464537 | 3.23E-06 | 0.001005776 |
| PSMA7 | 375.1244594 | 0.451182318 | 0.096914427 | 4.655471137 | 3.23E-06 | 0.001005776 |
| RN7SL792P | 10.98807619 | -0.821341194 | 0.176702277 | -4.648164187 | 3.35E-06 | 0.001013985 |
| SEM1 | 46.0244404 | 0.550764335 | 0.118670209 | 4.641133922 | 3.47E-06 | 0.001027709 |
| ATP5MD | 117.3489001 | 0.450250163 | 0.097198783 | 4.632261339 | 3.62E-06 | 0.001027709 |
| PGAP2 | 38.27253201 | -0.470158302 | 0.101471371 | -4.633408373 | 3.60E-06 | 0.001027709 |
| YWHAE | 283.1230545 | 0.430140687 | 0.092756553 | 4.637307787 | 3.53E-06 | 0.001027709 |
| AL049796.1 | 48.41284172 | -1.056369726 | 0.228576781 | -4.621509322 | 3.81E-06 | 0.001050137 |

| | | | | | | |
|---|---|---|---|---|---|---|
| UBE3AP2 | 9.282123992 | -1.389105531 | 0.300567475 | -4.621609605 | 3.81E-06 | 0.001050137 |
| BMP2K | 131.4268368 | -0.646601612 | 0.140115649 | -4.614770842 | 3.94E-06 | 0.001067803 |
| FO393411.1 | 10.04874504 | 1.002533873 | 0.217377954 | 4.611939051 | 3.99E-06 | 0.001067803 |
| AHNAK | 914.9393572 | -0.327940819 | 0.07139165 | -4.593545846 | 4.36E-06 | 0.001149767 |
| NHLRC2 | 74.14538096 | -0.275454255 | 0.060063244 | -4.586070251 | 4.52E-06 | 0.001174906 |
| SHOC2 | 149.5944909 | -0.25199684 | 0.055485582 | -4.541663445 | 5.58E-06 | 0.001431658 |
| SNORA79B | 22.96054395 | -0.650709097 | 0.143412092 | -4.537337739 | 5.70E-06 | 0.001441311 |
| TUBA1C | 63.6370561 | 0.581902274 | 0.128391976 | 4.532232402 | 5.84E-06 | 0.001456645 |
| PCGF2 | 145.7671074 | 0.939405971 | 0.207945946 | 4.517548869 | 6.26E-06 | 0.001540551 |
| RPL3 | 101.0702744 | 0.794927144 | 0.176498574 | 4.503872895 | 6.67E-06 | 0.001621533 |
| AL139287.1 | 13.50562642 | -0.641678648 | 0.142825123 | -4.492757555 | 7.03E-06 | 0.001653485 |
| RN7SKP55 | 14.36953712 | -0.769414537 | 0.171334512 | -4.490715432 | 7.10E-06 | 0.001653485 |
| RPL24 | 536.6438059 | 0.308881329 | 0.068739713 | 4.493491692 | 7.01E-06 | 0.001653485 |
| ANXA2 | 278.1435947 | 0.452086879 | 0.10071701 | 4.48868446 | 7.17E-06 | 0.001653485 |
| ABHD12 | 57.83294018 | 0.4313449 | 0.096150077 | 4.486162802 | 7.25E-06 | 0.001653485 |
| ACOX2 | 14.73090749 | 0.911394253 | 0.203715277 | 4.47386305 | 7.68E-06 | 0.001709361 |
| SNORA38B | 13.63095322 | -0.963405018 | 0.215239831 | -4.475960662 | 7.61E-06 | 0.001709361 |
| RN7SL538P | 4.564666293 | -1.23348814 | 0.276867763 | -4.455152626 | 8.38E-06 | 0.001760795 |
| HAX1 | 46.62915152 | 0.458236959 | 0.102728544 | 4.460658544 | 8.17E-06 | 0.001760795 |
| RPS18 | 130.913108 | 0.966362961 | 0.216895595 | 4.455429163 | 8.37E-06 | 0.001760795 |
| NDUFB8 | 59.95956613 | 0.456333108 | 0.102431915 | 4.454989506 | 8.39E-06 | 0.001760795 |
| UBALD2 | 95.23582922 | 0.494573451 | 0.110998064 | 4.4556944 | 8.36E-06 | 0.001760795 |
| RPL7A | 379.1105429 | 0.472528417 | 0.106326115 | 4.444142585 | 8.82E-06 | 0.001790946 |
| TPM1-AS | 12.70093754 | -0.738844403 | 0.166149053 | -4.446877006 | 8.71E-06 | 0.001790946 |
| ST13 | 176.7960512 | 0.347779681 | 0.078238878 | 4.445100554 | 8.79E-06 | 0.001790946 |
| RN7SL687P | 14.15712716 | -0.978885735 | 0.2205517 | -4.438350436 | 9.07E-06 | 0.001819818 |
| NAA10 | 57.51392728 | 0.316622048 | 0.071499255 | 4.428326538 | 9.50E-06 | 0.001885964 |
| GLYR1 | 145.3602068 | -0.264625983 | 0.059872393 | -4.419833089 | 9.88E-06 | 0.001940761 |
| PRLR | 317.8228786 | -0.644051195 | 0.146233476 | -4.404266465 | 1.06E-05 | 0.00206352 |
| EIF5A | 66.82925428 | 0.641656736 | 0.146112259 | 4.391532516 | 1.13E-05 | 0.002165384 |
| H1-4 | 39.97542838 | -0.583236973 | 0.133211155 | -4.37828928 | 1.20E-05 | 0.002277485 |
| PUM1 | 290.4605693 | -0.164825664 | 0.037672296 | -4.375248728 | 1.21E-05 | 0.002285899 |
| SRP9 | 39.3005666 | 0.691695357 | 0.158900119 | 4.353019766 | 1.34E-05 | 0.002373593 |
| HSPE1 | 81.57866085 | 0.516401323 | 0.118339919 | 4.363711984 | 1.28E-05 | 0.002373593 |
| SAP30 | 15.78744545 | 0.549702145 | 0.126260171 | 4.353725633 | 1.34E-05 | 0.002373593 |

| | | | | | | |
|---|---|---|---|---|---|---|
| TRAF6 | 24.79401872 | -0.364005041 | 0.083682146 | -4.3498531 | 1.36E-05 | 0.002373593 |
| PPP1R14B-AS1 | 6.644117505 | 1.022523649 | 0.23467706 | 4.357152128 | 1.32E-05 | 0.002373593 |
| RN7SKP118 | 4.19558005 | -0.911872731 | 0.209535845 | -4.351869875 | 1.35E-05 | 0.002373593 |
| CYP4F3 | 3.913730574 | -1.714694439 | 0.393089647 | -4.362095137 | 1.29E-05 | 0.002373593 |
| GRPR | 13.1439896 | -1.838554058 | 0.421687695 | -4.359989821 | 1.30E-05 | 0.002373593 |
| UBN2 | 172.2294121 | -0.316412794 | 0.072857569 | -4.342895331 | 1.41E-05 | 0.002427157 |
| SERPINA1 | 273.121142 | -1.292489966 | 0.298356274 | -4.332035488 | 1.48E-05 | 0.002526442 |
| RN7SL4P | 6.087373825 | -0.805353008 | 0.186156645 | -4.326211449 | 1.52E-05 | 0.002546973 |
| SMIM26 | 20.64385721 | 0.637840489 | 0.14740574 | 4.327107537 | 1.51E-05 | 0.002546973 |
| SPOPL | 138.886728 | -0.422679626 | 0.097809833 | -4.321443068 | 1.55E-05 | 0.002558692 |
| ATF4 | 260.9596996 | 0.364532318 | 0.084358538 | 4.321226105 | 1.55E-05 | 0.002558692 |
| RPL23AP42 | 8.329009739 | 1.090358204 | 0.252455667 | 4.319008631 | 1.57E-05 | 0.002561661 |
| PSMB4 | 148.3354661 | 0.335734509 | 0.077778138 | 4.31656654 | 1.58E-05 | 0.002567433 |
| LRRC37B | 5.99740319 | 1.003068828 | 0.232668752 | 4.311145443 | 1.62E-05 | 0.002601188 |
| PSMB3 | 219.0206667 | 0.855877018 | 0.198586889 | 4.309836483 | 1.63E-05 | 0.002601188 |
| PLAC9P1 | 21.20805769 | -2.109389451 | 0.489740538 | -4.30715713 | 1.65E-05 | 0.002610382 |
| EPGN | 2.704840764 | 1.746921836 | 0.406967488 | 4.292534138 | 1.77E-05 | 0.002706917 |
| MDM1 | 22.99677504 | -0.426774146 | 0.099380472 | -4.294346126 | 1.75E-05 | 0.002706917 |
| MAPK4 | 2.030095115 | -2.213961395 | 0.515499619 | -4.294787649 | 1.75E-05 | 0.002706917 |
| PTOV1 | 101.4363404 | 0.404625321 | 0.094281825 | 4.291657701 | 1.77E-05 | 0.002706917 |
| RPL21 | 69.82498055 | 0.789330365 | 0.184044352 | 4.288805141 | 1.80E-05 | 0.002719437 |
| SCARNA3 | 135.2304261 | -0.485868811 | 0.113682248 | -4.273919807 | 1.92E-05 | 0.002883959 |
| RPL18A | 29.5148327 | 0.942582722 | 0.220732328 | 4.270252264 | 1.95E-05 | 0.002908153 |
| S100P | 88.16422309 | 1.543687686 | 0.361686966 | 4.268021334 | 1.97E-05 | 0.002908991 |
| HINT1 | 202.9082683 | 0.35761794 | 0.083852341 | 4.264853403 | 2.00E-05 | 0.002908991 |
| NONO | 264.6021583 | 0.205337887 | 0.048129756 | 4.266339671 | 1.99E-05 | 0.002908991 |
| ITGB6 | 65.33000799 | 0.852650614 | 0.200438837 | 4.253919178 | 2.10E-05 | 0.00293671 |
| BRK1 | 145.473739 | 0.35973217 | 0.084727117 | 4.245773771 | 2.18E-05 | 0.00293671 |
| GYG1 | 35.85244914 | 0.399279633 | 0.093932823 | 4.250693442 | 2.13E-05 | 0.00293671 |
| MDM2 | 182.4870423 | -0.429354165 | 0.101009272 | -4.250641122 | 2.13E-05 | 0.00293671 |
| COX5A | 38.21079858 | 0.485477084 | 0.114103074 | 4.254723967 | 2.09E-05 | 0.00293671 |
| RPL17 | 6.93067623 | -0.651300854 | 0.153006555 | -4.256685968 | 2.07E-05 | 0.00293671 |
| ZNF236 | 53.86025938 | -0.360110626 | 0.084570849 | -4.258094014 | 2.06E-05 | 0.00293671 |
| C19orf48 | 28.40356105 | 0.495145276 | 0.116610455 | 4.246148218 | 2.17E-05 | 0.00293671 |
| CHD6 | 200.0807793 | -0.349713281 | 0.082294199 | -4.249549626 | 2.14E-05 | 0.00293671 |

| | | | | | | |
|---|---|---|---|---|---|---|
| AC091807.2 | 1.410480326 | -1.257951497 | 0.29610457 | -4.248335299 | 2.15E-05 | 0.00293671 |
| SPIN1 | 141.6927882 | -0.270644767 | 0.063829508 | -4.240119894 | 2.23E-05 | 0.0029752 |
| PRPF39 | 57.15102978 | -0.374992392 | 0.088449921 | -4.239601185 | 2.24E-05 | 0.0029752 |
| SP3 | 158.2586547 | -0.209997773 | 0.049609414 | -4.233022641 | 2.31E-05 | 0.002982281 |
| RPL34 | 1254.263765 | 0.342221526 | 0.080843275 | 4.233147702 | 2.30E-05 | 0.002982281 |
| LIFR | 113.0621355 | -0.482493502 | 0.113905871 | -4.235896698 | 2.28E-05 | 0.002982281 |
| C5orf46 | 5.045926432 | 1.140733782 | 0.26950541 | 4.232693441 | 2.31E-05 | 0.002982281 |
| RPS25 | 297.0356564 | 0.441798256 | 0.104664782 | 4.221078457 | 2.43E-05 | 0.003118392 |
| ANKIB1 | 161.3218013 | -0.242190451 | 0.057443087 | -4.216180991 | 2.48E-05 | 0.003134106 |
| SLC30A8 | 10.01287006 | -2.365458185 | 0.561160827 | -4.215294563 | 2.49E-05 | 0.003134106 |
| RPL13A | 503.7713841 | 0.512409408 | 0.121493385 | 4.217590998 | 2.47E-05 | 0.003134106 |
| MAPT-AS1 | 4.030432657 | -2.06411617 | 0.490685594 | -4.206596232 | 2.59E-05 | 0.003235141 |
| CFAP70 | 53.09493332 | -0.625018913 | 0.148765355 | -4.201374134 | 2.65E-05 | 0.00328847 |
| CA3-AS1 | 10.67661156 | -0.730120369 | 0.174015671 | -4.195716204 | 2.72E-05 | 0.003349177 |
| SPEF2 | 48.88264036 | -0.533098021 | 0.127153739 | -4.192546944 | 2.76E-05 | 0.003373833 |
| CPLANE1 | 182.6897201 | -0.342603248 | 0.081791821 | -4.188722598 | 2.81E-05 | 0.003408613 |
| RPL10 | 112.7547738 | 1.016781397 | 0.243027418 | 4.183813515 | 2.87E-05 | 0.003460334 |
| S100A11 | 543.2076174 | 0.449230652 | 0.107643877 | 4.173304289 | 3.00E-05 | 0.003590429 |
| PFN1 | 621.5489777 | 0.365878563 | 0.087688871 | 4.172462938 | 3.01E-05 | 0.003590429 |
| LEPR | 91.65777614 | -0.693744636 | 0.166382288 | -4.169582247 | 3.05E-05 | 0.003612803 |
| SLC25A5 | 76.90730732 | 0.587370319 | 0.141085128 | 4.163233417 | 3.14E-05 | 0.003691122 |
| HPCAL1 | 48.18337419 | 0.391725049 | 0.094345115 | 4.152043808 | 3.30E-05 | 0.003851829 |
| PDAP1 | 30.18071398 | 0.463742959 | 0.111753759 | 4.149685555 | 3.33E-05 | 0.00386725 |
| MED1 | 384.2144304 | 0.826685387 | 0.199297002 | 4.148007139 | 3.35E-05 | 0.003871356 |
| POLRMT | 24.3545474 | 0.359415559 | 0.086679741 | 4.146477095 | 3.38E-05 | 0.003873098 |
| ZYG11B | 148.0410247 | -0.23618083 | 0.057061357 | -4.139067912 | 3.49E-05 | 0.00396231 |
| ACTG1 | 1669.756289 | 0.245371732 | 0.059293008 | 4.138291152 | 3.50E-05 | 0.00396231 |
| SCGB1B2P | 5.961605129 | -1.540620154 | 0.37239838 | -4.137021626 | 3.52E-05 | 0.00396231 |
| ARF4 | 72.03259491 | 0.389450099 | 0.094201708 | 4.134214827 | 3.56E-05 | 0.003986732 |
| NBR1 | 165.5400963 | -0.303389187 | 0.07346141 | -4.129912398 | 3.63E-05 | 0.004037607 |
| RPS8 | 1302.818878 | 0.475618135 | 0.115204163 | 4.128480461 | 3.65E-05 | 0.004038501 |
| AGL | 115.6001767 | -0.478745161 | 0.116037026 | -4.125796542 | 3.69E-05 | 0.004038759 |
| MAP4K5 | 113.519063 | -0.271620684 | 0.065835835 | -4.125727043 | 3.70E-05 | 0.004038759 |
| NPM1 | 122.9954533 | 0.344991542 | 0.083684523 | 4.122525045 | 3.75E-05 | 0.004045325 |
| ACTB | 205.2198469 | 0.353471947 | 0.085733108 | 4.122933999 | 3.74E-05 | 0.004045325 |

| | | | | | | |
|---|---|---|---|---|---|---|
| MYBPC1 | 25.28044899 | -1.753582039 | 0.42551623 | -4.121069693 | 3.77E-05 | 0.004045325 |
| RNU1-98P | 7.059753154 | -0.801413675 | 0.194519464 | -4.119966505 | 3.79E-05 | 0.004045325 |
| TOMM22 | 52.27596665 | 0.396194936 | 0.096327515 | 4.112998602 | 3.91E-05 | 0.004145471 |
| STC1 | 144.4066718 | -1.156027127 | 0.281354551 | -4.108791283 | 3.98E-05 | 0.004197585 |
| LINC01238 | 10.89801878 | -1.043217603 | 0.254088139 | -4.105731212 | 4.03E-05 | 0.004197907 |
| MYOZ3 | 4.105980941 | -1.180251172 | 0.28760511 | -4.103721149 | 4.07E-05 | 0.004197907 |
| RPS13 | 158.2282525 | 0.736560697 | 0.179493515 | 4.10355047 | 4.07E-05 | 0.004197907 |
| UBTF | 165.7905472 | -0.204703503 | 0.049849129 | -4.106460975 | 4.02E-05 | 0.004197907 |
| RPS29 | 1277.612646 | 0.419854473 | 0.102482942 | 4.096823 | 4.19E-05 | 0.004291088 |
| SNRPD2 | 228.6945186 | 0.259249673 | 0.063294957 | 4.095897763 | 4.21E-05 | 0.004291088 |
| LTN1 | 118.1439656 | -0.284354847 | 0.06947272 | -4.09304323 | 4.26E-05 | 0.00432041 |
| UFC1 | 168.6172271 | 0.389041245 | 0.095205322 | 4.086339262 | 4.38E-05 | 0.004398762 |
| ANKRD30B | 108.798 | -1.276354233 | 0.312307753 | -4.08684774 | 4.37E-05 | 0.004398762 |
| MRPL45 | 50.04261082 | 0.792764801 | 0.19407996 | 4.084732913 | 4.41E-05 | 0.00440536 |
| XAB2 | 23.10314669 | -0.480125984 | 0.117739491 | -4.077867 | 4.55E-05 | 0.004513064 |
| RNVU1-1 | 3.626955286 | -0.964741576 | 0.23674961 | -4.074944738 | 4.60E-05 | 0.004521494 |
| LINC02533 | 27.69066507 | -0.624015804 | 0.153116177 | -4.075440087 | 4.59E-05 | 0.004521494 |
| HES6 | 8.693118628 | 0.898119672 | 0.220590688 | 4.071430576 | 4.67E-05 | 0.004565975 |
| HOXA7 | 25.02878329 | -0.898842471 | 0.22092293 | -4.068579352 | 4.73E-05 | 0.004597872 |
| SNRPF | 113.7099771 | 0.400447305 | 0.098705091 | 4.057007592 | 4.97E-05 | 0.004806336 |
| KLHL9 | 91.46355542 | -0.253681502 | 0.062609226 | -4.051822996 | 5.08E-05 | 0.004857642 |
| RPL36 | 531.12276 | 0.482174562 | 0.118980136 | 4.052563551 | 5.07E-05 | 0.004857642 |
| ATP5PO | 133.5992022 | 0.302173142 | 0.074594421 | 4.050881245 | 5.10E-05 | 0.004857642 |
| ADIPOR1 | 81.70680689 | 0.384465675 | 0.095091174 | 4.043126792 | 5.27E-05 | 0.004995446 |
| ZNF638 | 373.4263824 | -0.249830514 | 0.061861729 | -4.038531086 | 5.38E-05 | 0.005042596 |
| LMLN | 61.4053248 | -0.337460805 | 0.083560055 | -4.038542169 | 5.38E-05 | 0.005042596 |
| NIPBL | 455.218356 | -0.217096561 | 0.053952797 | -4.02382402 | 5.73E-05 | 0.005314312 |
| SERF2 | 427.4876578 | 0.4180958 | 0.103894283 | 4.024242614 | 5.72E-05 | 0.005314312 |
| TAC1 | 11.60627334 | -1.241626908 | 0.308860761 | -4.020021523 | 5.82E-05 | 0.005357052 |
| SLC24A1 | 32.61989924 | -0.381058222 | 0.094800446 | -4.01958259 | 5.83E-05 | 0.005357052 |
| CHST1 | 37.91158676 | 1.180673228 | 0.294010001 | 4.015758728 | 5.93E-05 | 0.005417712 |
| YWHAQ | 55.93278983 | 0.361940199 | 0.090213826 | 4.012025839 | 6.02E-05 | 0.005450157 |
| PGK1 | 248.5440294 | 0.334039302 | 0.083237414 | 4.013090832 | 5.99E-05 | 0.005450157 |
| SAA2 | 25.93375213 | 1.177521639 | 0.294054271 | 4.004436437 | 6.22E-05 | 0.005600657 |
| AC105914.2 | 3.26446627 | -3.764691276 | 0.940518419 | -4.002783148 | 6.26E-05 | 0.005612575 |

| | | | | | | |
|---|---|---|---|---|---|---|
| CNBP | 312.5235761 | -0.247942118 | 0.062013315 | -3.998207789 | 6.38E-05 | 0.0056727 |
| STEAP4 | 225.7639452 | -0.707015117 | 0.17684329 | -3.997975365 | 6.39E-05 | 0.0056727 |
| AP3S1 | 19.01315963 | 0.442810598 | 0.110906652 | 3.992642371 | 6.53E-05 | 0.00576415 |
| HOXC4 | 33.79185955 | -0.517737906 | 0.12969649 | -3.99191917 | 6.55E-05 | 0.00576415 |
| ASS1 | 52.046875 | 0.675858492 | 0.16935935 | 3.990677178 | 6.59E-05 | 0.005766959 |
| FAM120B | 51.07212687 | -0.326173167 | 0.081811814 | -3.986871213 | 6.70E-05 | 0.00582043 |
| TPSG1 | 3.059828989 | -2.092657165 | 0.524969009 | -3.98624896 | 6.71E-05 | 0.00582043 |
| CALY | 2.169408912 | 1.823056739 | 0.457580634 | 3.984121277 | 6.77E-05 | 0.005845376 |
| RPS21 | 1418.150291 | 0.374322452 | 0.094268305 | 3.970819807 | 7.16E-05 | 0.006152818 |
| SCNM1 | 42.24428082 | 0.304719203 | 0.076799138 | 3.967742462 | 7.26E-05 | 0.006188083 |
| PON3 | 27.17061445 | -1.063799137 | 0.268145147 | -3.967251133 | 7.27E-05 | 0.006188083 |
| SEC11A | 110.4232546 | 0.248006946 | 0.062556111 | 3.964551879 | 7.35E-05 | 0.00622981 |
| TIMM8B | 67.13359531 | 0.355532832 | 0.089820722 | 3.958249551 | 7.55E-05 | 0.006367255 |
| PKHD1 | 1.993540141 | -1.289318763 | 0.325928084 | -3.95583819 | 7.63E-05 | 0.006402595 |
| AC090740.1 | 7.351742718 | -1.431911621 | 0.362512654 | -3.949963136 | 7.82E-05 | 0.006532112 |
| SERPINA6 | 15.22547318 | -2.724786841 | 0.691411689 | -3.94090364 | 8.12E-05 | 0.006722984 |
| TMPRSS3 | 17.79263844 | -1.167445033 | 0.296160621 | -3.941932016 | 8.08E-05 | 0.006722984 |
| EEF1B2 | 285.0529478 | 0.293612423 | 0.074804717 | 3.925052246 | 8.67E-05 | 0.006801907 |
| MEGF10 | 9.259901774 | -1.613973994 | 0.411051276 | -3.926454164 | 8.62E-05 | 0.006801907 |
| INSYN2B | 4.281011477 | -1.091454576 | 0.277942094 | -3.926913548 | 8.60E-05 | 0.006801907 |
| PPT2-EGFL8 | 5.669866361 | -0.753495844 | 0.191805709 | -3.928432823 | 8.55E-05 | 0.006801907 |
| HMGA1 | 144.6762244 | 0.418287612 | 0.10658411 | 3.924483774 | 8.69E-05 | 0.006801907 |
| PHF3 | 338.1215729 | -0.206140391 | 0.05247307 | -3.928498803 | 8.55E-05 | 0.006801907 |
| LCN2 | 29.14170097 | 1.577982266 | 0.401465957 | 3.930550621 | 8.48E-05 | 0.006801907 |
| TUBB4B | 137.0678946 | 0.441246793 | 0.112239581 | 3.931294007 | 8.45E-05 | 0.006801907 |
| GAPDH | 1276.382898 | 0.337391798 | 0.085825021 | 3.931158913 | 8.45E-05 | 0.006801907 |
| NABP2 | 47.69766163 | 0.352335611 | 0.08973297 | 3.926490032 | 8.62E-05 | 0.006801907 |
| RPLP0 | 569.8662546 | 0.436139064 | 0.111091637 | 3.925939671 | 8.64E-05 | 0.006801907 |
| RPLP1 | 2423.281833 | 0.390180813 | 0.099406467 | 3.925104908 | 8.67E-05 | 0.006801907 |
| ABHD17A | 11.25389565 | 0.438328905 | 0.11160742 | 3.927417221 | 8.59E-05 | 0.006801907 |
| SNORA66 | 18.56775491 | -0.788658541 | 0.201365497 | -3.916552494 | 8.98E-05 | 0.006928286 |
| PPFIA1 | 223.5504626 | -0.450739264 | 0.115102043 | -3.915997092 | 9.00E-05 | 0.006928286 |
| IDH2 | 162.91855 | 0.464554972 | 0.118595554 | 3.917136478 | 8.96E-05 | 0.006928286 |
| COX4I1 | 245.241108 | 0.335961676 | 0.085782058 | 3.916456216 | 8.99E-05 | 0.006928286 |
| LINC01488 | 6.861246713 | -2.020170205 | 0.517455431 | -3.904046769 | 9.46E-05 | 0.007249465 |

| | | | | | | |
|---|---|---|---|---|---|---|
| CAPN13 | 24.08514517 | 0.797672621 | 0.204741758 | 3.895993809 | 9.78E-05 | 0.007372289 |
| RPL32 | 1304.09893 | 0.28944295 | 0.074268803 | 3.897234627 | 9.73E-05 | 0.007372289 |
| RPL35A | 817.1587448 | 0.309767679 | 0.079507692 | 3.896071806 | 9.78E-05 | 0.007372289 |
| RNU1-82P | 2.911235022 | -0.864720135 | 0.221787627 | -3.898865539 | 9.66E-05 | 0.007372289 |
| NUTF2 | 73.8103991 | 0.367079481 | 0.094282835 | 3.893386123 | 9.89E-05 | 0.007421731 |
| C1QTNF7 | 5.291500574 | -0.780446083 | 0.200817048 | -3.886353726 | 0.000101761 | 0.007609018 |
| SF3B4 | 7.726057223 | 0.73368487 | 0.18901458 | 3.881631082 | 0.000103758 | 0.007618566 |
| ITPRID2 | 144.1172567 | -0.529898939 | 0.13646445 | -3.883054803 | 0.000103152 | 0.007618566 |
| SEC13 | 105.7530725 | 0.292107675 | 0.075264254 | 3.881094429 | 0.000103987 | 0.007618566 |
| TIMM10B | 54.0324988 | -0.271908941 | 0.069999 | -3.884468919 | 0.000102554 | 0.007618566 |
| LINC01252 | 6.141153256 | -0.772406689 | 0.199062828 | -3.880215598 | 0.000104364 | 0.007618566 |
| ABCC13 | 3.278682432 | -1.801540319 | 0.46409445 | -3.881839828 | 0.000103669 | 0.007618566 |
| GRIA2 | 20.23424543 | -1.887734344 | 0.487254463 | -3.874226895 | 0.000106964 | 0.007769553 |
| RPL4 | 151.8164112 | 0.3751193 | 0.096841911 | 3.87352227 | 0.000107274 | 0.007769553 |
| RPL10A | 155.663745 | 0.539692235 | 0.139409385 | 3.871276209 | 0.000108267 | 0.007810874 |
| ATP5MF | 171.649868 | 0.251359914 | 0.065013211 | 3.86628981 | 0.000110504 | 0.007941211 |
| C1orf167 | 4.058718873 | -1.025125614 | 0.265448925 | -3.861856345 | 0.000112529 | 0.0080554 |
| CHCHD2 | 148.2363564 | 0.544892219 | 0.141143941 | 3.86054275 | 0.000113135 | 0.008067562 |
| TMSB4X | 181.807486 | 0.678590964 | 0.175945615 | 3.856822268 | 0.000114871 | 0.00815979 |
| GLYATL2 | 29.05282246 | 1.690502704 | 0.439224238 | 3.848837471 | 0.00011868 | 0.008384485 |
| PHB2 | 168.2852673 | 0.285260055 | 0.074133127 | 3.847943088 | 0.000119114 | 0.008384485 |
| NDUFA3 | 144.2521778 | 0.347932007 | 0.090433878 | 3.84736358 | 0.000119396 | 0.008384485 |
| PLEKHH1 | 33.16546946 | -0.557636027 | 0.145002836 | -3.84569049 | 0.000120213 | 0.008409931 |
| BIN2 | 14.11710961 | 0.77635426 | 0.202223747 | 3.839085522 | 0.000123493 | 0.008606792 |
| PRR11 | 39.13131471 | 0.709188118 | 0.184839949 | 3.836768626 | 0.000124664 | 0.0086557 |
| FAU | 494.8735704 | 0.313234714 | 0.081712561 | 3.833372882 | 0.000126398 | 0.008743248 |
| RPL29 | 156.4372388 | 0.426097376 | 0.111269641 | 3.829412713 | 0.000128449 | 0.008851987 |
| RARRES1 | 94.09311778 | 0.798674411 | 0.208731501 | 3.826324291 | 0.000130071 | 0.008864148 |
| RPL7 | 274.1420751 | 0.660618058 | 0.172632979 | 3.826719912 | 0.000129862 | 0.008864148 |
| LDHA | 70.83149445 | 0.476530116 | 0.124502843 | 3.827463742 | 0.00012947 | 0.008864148 |
| ATXN2L | 130.7988804 | 0.244960658 | 0.064034856 | 3.825426858 | 0.000130546 | 0.008864148 |
| PMF1 | 52.22956752 | 0.330670595 | 0.086504755 | 3.82257134 | 0.000132067 | 0.008897107 |
| GSTO1 | 88.44218148 | 0.303677442 | 0.079454574 | 3.822025912 | 0.00013236 | 0.008897107 |
| TRAPPC8 | 86.72664071 | -0.23203944 | 0.060728779 | -3.820913996 | 0.000132958 | 0.008897107 |
| MT-TY | 518.9411927 | -0.639378069 | 0.167329637 | -3.821068874 | 0.000132875 | 0.008897107 |

| | | | | | | |
|---|---|---|---|---|---|---|
| DDIT4 | 69.82060523 | 0.709633309 | 0.186019557 | 3.814831726 | 0.000136276 | 0.009053515 |
| GMFG | 39.78313741 | 0.561562362 | 0.147193599 | 3.815127592 | 0.000136113 | 0.009053515 |
| SDAD1 | 85.08075302 | 0.2570975 | 0.067467935 | 3.81066204 | 0.000138595 | 0.009076998 |
| ACHE | 5.229369725 | 1.159585075 | 0.304168297 | 3.81231406 | 0.000137672 | 0.009076998 |
| NOP10 | 48.44125891 | 0.412490605 | 0.108215481 | 3.811752275 | 0.000137985 | 0.009076998 |
| SNORD124 | 18.66320274 | -0.849106697 | 0.222813008 | -3.810848861 | 0.00013849 | 0.009076998 |
| CAP1 | 342.532488 | 0.317664513 | 0.083446461 | 3.806806308 | 0.000140773 | 0.009177697 |
| PFKFB3 | 65.68524797 | 0.514078218 | 0.135092324 | 3.805384358 | 0.000141584 | 0.009177697 |
| SP1 | 139.6033858 | -0.196617589 | 0.051669193 | -3.805315672 | 0.000141623 | 0.009177697 |
| DEAF1 | 49.21087868 | -0.281549014 | 0.074022363 | -3.80356693 | 0.000142627 | 0.009210441 |
| PSMB7 | 129.2337906 | 0.254027826 | 0.066803334 | 3.802621988 | 0.000143173 | 0.009213439 |
| EVC | 12.09110958 | -0.559955196 | 0.147414652 | -3.798504344 | 0.000145572 | 0.009251404 |
| CCDC158 | 9.548330664 | -0.898127431 | 0.236462988 | -3.798173396 | 0.000145766 | 0.009251404 |
| RPL37 | 975.5412724 | 0.391958832 | 0.103139834 | 3.800266269 | 0.000144541 | 0.009251404 |
| KPNA2 | 14.36145169 | 0.586218828 | 0.154288071 | 3.799508449 | 0.000144983 | 0.009251404 |
| SURF4 | 121.1264538 | 0.277080101 | 0.072995399 | 3.795857052 | 0.000147134 | 0.009294858 |
| SKA2 | 33.02267907 | 0.551278593 | 0.145252487 | 3.795312594 | 0.000147458 | 0.009294858 |
| UQCRH | 209.0941741 | 0.321409079 | 0.08474346 | 3.792730192 | 0.000149 | 0.009360141 |
| SMIM4 | 56.50919666 | 0.376550892 | 0.09935015 | 3.790139152 | 0.000150563 | 0.009426259 |
| TRIM8 | 177.563309 | 0.265678751 | 0.070252183 | 3.781786399 | 0.000155707 | 0.009715377 |
| MTRNR2L5 | 9.945154961 | -0.580075613 | 0.153459231 | -3.779998169 | 0.00015683 | 0.009752472 |
| KMT5A | 13.1889105 | 0.483828205 | 0.128126485 | 3.776176372 | 0.000159254 | 0.009870022 |
| CP | 60.77218675 | 1.300202045 | 0.344573199 | 3.773369633 | 0.000161057 | 0.009882289 |
| RN7SL183P | 1.993160397 | -0.99993302 | 0.264994152 | -3.773415417 | 0.000161028 | 0.009882289 |
| CAPN15 | 11.64017238 | 0.566819995 | 0.150153397 | 3.77493955 | 0.000160046 | 0.009882289 |
| OXCT1 | 22.48793788 | 0.492947783 | 0.130838771 | 3.767597173 | 0.000164826 | 0.010080062 |
| AC114490.3 | 8.183384202 | -0.59501883 | 0.158075327 | -3.764147395 | 0.000167118 | 0.010086627 |
| RN7SKP174 | 2.606052374 | -0.994901922 | 0.264148649 | -3.766447132 | 0.000165587 | 0.010086627 |
| CIAPIN1 | 16.59657453 | 0.422518579 | 0.112227239 | 3.764848726 | 0.00016665 | 0.010086627 |
| AC142472.1 | 14.74408313 | -0.620091555 | 0.164681956 | -3.765388584 | 0.00016629 | 0.010086627 |
| MRPL51 | 115.7350199 | 0.283632794 | 0.075378593 | 3.762776419 | 0.000168037 | 0.010109065 |
| TAGLN2 | 595.2911234 | 0.346843391 | 0.092303396 | 3.757644964 | 0.00017152 | 0.010245471 |
| RPS14 | 577.2741031 | 0.424068645 | 0.112874559 | 3.756990489 | 0.000171969 | 0.010245471 |
| AC015961.2 | 5.475669543 | -0.767388947 | 0.20422201 | -3.757621168 | 0.000171536 | 0.010245471 |
| AC004223.2 | 4.364597999 | -1.01300009 | 0.269767352 | -3.755087797 | 0.000173281 | 0.010290424 |

| | | | | | | |
|---|---|---|---|---|---|---|
| NCR3LG1 | 6.003056991 | -0.830130941 | 0.221150937 | -3.753684934 | 0.000174254 | 0.010315047 |
| SH3D21 | 46.09389803 | -0.516760594 | 0.137718049 | -3.752308422 | 0.000175214 | 0.010338729 |
| ENO1 | 534.7875281 | 0.322424817 | 0.085982629 | 3.74988321 | 0.000176917 | 0.01037295 |
| RPL12 | 763.3258525 | 0.422400954 | 0.112631933 | 3.750277037 | 0.000176639 | 0.01037295 |
| SYNPO2L | 3.227738206 | -1.368288524 | 0.36528527 | -3.745808106 | 0.000179814 | 0.010509446 |
| UQCRB | 225.3373121 | 0.285945199 | 0.076424472 | 3.741539756 | 0.000182896 | 0.010655869 |
| MRPL52 | 82.93673035 | 0.272405383 | 0.072845649 | 3.739487362 | 0.000184396 | 0.01070946 |
| RPL6 | 66.55877097 | 0.397345185 | 0.106318096 | 3.737324108 | 0.000185989 | 0.010768128 |
| MT-CO1 | 639.1680272 | 0.475322035 | 0.127276192 | 3.734571458 | 0.000188035 | 0.010852563 |
| GLYATL1 | 6.342687631 | 1.520983353 | 0.407738488 | 3.730291344 | 0.000191258 | 0.011004215 |
| AXDND1 | 1.554791223 | -1.287343324 | 0.345405405 | -3.727050313 | 0.000193734 | 0.011007629 |
| ZNF148 | 204.2361996 | -0.170264833 | 0.045692522 | -3.726317228 | 0.000194298 | 0.011007629 |
| RPS12 | 1075.670882 | 0.405318692 | 0.10869266 | 3.729034626 | 0.000192215 | 0.011007629 |
| CHPF2 | 47.37190101 | 0.310132408 | 0.0832114 | 3.727042294 | 0.00019374 | 0.011007629 |
| GLTP | 43.20255067 | 0.426800618 | 0.114528834 | 3.726577868 | 0.000194097 | 0.011007629 |
| FABP5 | 4.392197764 | 0.894528535 | 0.240835009 | 3.714279489 | 0.000203783 | 0.011439748 |
| UBOX5 | 27.73316109 | -0.286719421 | 0.077192061 | -3.714364125 | 0.000203715 | 0.011439748 |
| MT-ATP6 | 3512.373259 | 0.519359002 | 0.139821533 | 3.714442211 | 0.000203652 | 0.011439748 |
| S100A8 | 143.683736 | 1.35437753 | 0.364828228 | 3.712370441 | 0.000205327 | 0.011491477 |
| EXOC4 | 119.9866252 | -0.244093329 | 0.065767691 | -3.711447429 | 0.000206077 | 0.011498625 |
| C1orf43 | 126.1701956 | 0.320888677 | 0.086537386 | 3.708093021 | 0.000208826 | 0.011549669 |
| PKM | 305.56672 | 0.288382191 | 0.07775697 | 3.708763225 | 0.000208274 | 0.011549669 |
| COX7B | 42.30180698 | 0.388164256 | 0.104681742 | 3.708041617 | 0.000208868 | 0.011549669 |
| NDUFB5 | 51.79280146 | 0.31006678 | 0.083693926 | 3.704770397 | 0.000211582 | 0.011652057 |
| FAR1 | 88.49882607 | -0.341608313 | 0.09221959 | -3.70429225 | 0.000211982 | 0.011652057 |
| TRPM7 | 141.771731 | -0.316635257 | 0.085520085 | -3.702466595 | 0.000213513 | 0.011701426 |
| IGIP | 35.77465112 | -0.353479354 | 0.095531824 | -3.700121465 | 0.000215496 | 0.011775151 |
| AC090994.1 | 4.925311542 | -1.524271351 | 0.412105923 | -3.698736823 | 0.000216675 | 0.01180464 |
| DNAJC8 | 85.87316968 | 0.221144944 | 0.059817896 | 3.696969609 | 0.000218188 | 0.011852125 |
| SLC30A2 | 7.53052349 | -1.047285442 | 0.283558666 | -3.69336426 | 0.000221307 | 0.011986254 |
| EIF4E2 | 99.52695508 | 0.201422727 | 0.054647101 | 3.685881301 | 0.000227913 | 0.012307945 |
| PSMD4 | 125.4829569 | 0.23111236 | 0.062846749 | 3.677395599 | 0.000235627 | 0.012687474 |
| TPRKB | 38.24777236 | 0.301645754 | 0.08210302 | 3.673990975 | 0.000238791 | 0.012820448 |
| NPY1R | 170.7093629 | -1.44989384 | 0.395308626 | -3.667751591 | 0.000244693 | 0.01307443 |
| ATG2B | 45.05483699 | -0.272855949 | 0.074398436 | -3.66749576 | 0.000244938 | 0.01307443 |

| | | | | | | |
|---|---|---|---|---|---|---|
| BOLA1 | 23.31223142 | 0.400361891 | 0.109256004 | 3.664438364 | 0.000247882 | 0.013193462 |
| CASD1 | 71.76430907 | -0.414378256 | 0.113146118 | -3.662328523 | 0.000249933 | 0.013264407 |
| ANP32B | 270.719707 | 0.209258049 | 0.057153753 | 3.661317714 | 0.000250921 | 0.013278701 |
| EIF1AX | 234.0061844 | -0.297588565 | 0.081328425 | -3.659096622 | 0.000253106 | 0.013356037 |
| NAT1 | 86.36627672 | -1.193418073 | 0.326410048 | -3.656192818 | 0.000255989 | 0.01346968 |
| TMSB10 | 1588.032199 | 0.380822567 | 0.104374689 | 3.648610298 | 0.000263663 | 0.013640298 |
| MTMR12 | 86.04507771 | -0.270689188 | 0.07418529 | -3.648825645 | 0.000263442 | 0.013640298 |
| SLC2A6 | 12.39268987 | 0.636078596 | 0.174323892 | 3.648832008 | 0.000263435 | 0.013640298 |
| ATP5F1B | 215.6879498 | 0.312824034 | 0.085724887 | 3.649162393 | 0.000263097 | 0.013640298 |
| CDK12 | 447.1656719 | 0.683404784 | 0.187271053 | 3.649281477 | 0.000262975 | 0.013640298 |
| RPS11 | 1750.014946 | 0.284763677 | 0.07802715 | 3.649546065 | 0.000262704 | 0.013640298 |
| RPS16 | 775.7233636 | 0.316956167 | 0.086902122 | 3.647277649 | 0.000265033 | 0.013672915 |
| FBXW5 | 117.402097 | 0.19386439 | 0.053177998 | 3.645575173 | 0.000266794 | 0.01372542 |
| FH | 36.08099361 | 0.433836259 | 0.119158435 | 3.640835488 | 0.000271755 | 0.01376014 |
| THRB | 42.39690112 | -0.43038753 | 0.118157595 | -3.642487233 | 0.000270016 | 0.01376014 |
| CLN8 | 24.86740758 | 0.408298509 | 0.112154297 | 3.640507067 | 0.000272102 | 0.01376014 |
| MTRNR2L8 | 18.24463463 | -0.854843484 | 0.234621522 | -3.643499865 | 0.000268956 | 0.01376014 |
| RPS6KA4 | 27.44875848 | 0.350821156 | 0.096329203 | 3.641898249 | 0.000270635 | 0.01376014 |
| SCN8A | 13.06103407 | -0.919486308 | 0.252609177 | -3.639956071 | 0.000272685 | 0.01376014 |
| ARPC3 | 130.3215245 | 0.271875488 | 0.074652365 | 3.641887131 | 0.000270647 | 0.01376014 |
| SEC14L2 | 78.84864899 | -0.772220903 | 0.212634441 | -3.631683079 | 0.000281579 | 0.014131736 |
| SHANK3 | 57.83737697 | -0.334842012 | 0.092194979 | -3.63188989 | 0.000281353 | 0.014131736 |
| KCNS3 | 17.56196754 | 0.588246021 | 0.162044305 | 3.630155484 | 0.000283251 | 0.014146762 |
| ZNF451 | 122.0667071 | -0.18562595 | 0.051136483 | -3.63001009 | 0.00028341 | 0.014146762 |
| DHX16 | 89.85956075 | -0.589508049 | 0.162438314 | -3.629119479 | 0.00028439 | 0.01415739 |
| SSB | 157.9511254 | 0.179749447 | 0.049552144 | 3.627480722 | 0.0002862 | 0.014209221 |
| NACA | 167.6750193 | 0.277734779 | 0.076695454 | 3.621267797 | 0.000293163 | 0.014515884 |
| LINC00893 | 6.767707338 | -0.713464588 | 0.197110414 | -3.61961895 | 0.000295037 | 0.014569629 |
| PDCD4 | 458.3867938 | -0.434300814 | 0.120032406 | -3.618196358 | 0.000296663 | 0.014610865 |
| RPS28 | 237.8725866 | 0.590567246 | 0.163681641 | 3.608023745 | 0.000308538 | 0.015155301 |
| SNORD6 | 4.886602883 | -0.688157462 | 0.190816367 | -3.606385925 | 0.000310491 | 0.015210777 |
| KIAA2026 | 216.6541021 | -0.18282929 | 0.05072318 | -3.604452425 | 0.000312812 | 0.015283915 |
| GZMB | 14.70313888 | 1.062506433 | 0.295669408 | 3.593562284 | 0.000326188 | 0.015853577 |
| SUMO2 | 21.68305062 | 0.584370005 | 0.162614529 | 3.593590377 | 0.000326152 | 0.015853577 |
| NUP54 | 39.45600398 | -0.274497115 | 0.076527455 | -3.586910271 | 0.000334619 | 0.016192098 |

| | | | | | | |
|---|---|---|---|---|---|---|
| SNORD14E | 22.88455658 | -0.496528822 | 0.138436626 | -3.586686835 | 0.000334906 | 0.016192098 |
| MBTPS1 | 172.2626864 | -0.285175626 | 0.079542221 | -3.585210751 | 0.000336806 | 0.016241441 |
| ZFYVE16 | 151.4807366 | -0.229443718 | 0.064013132 | -3.584322651 | 0.000337954 | 0.01625436 |
| ZMYND11 | 204.6178334 | -0.22899941 | 0.063934173 | -3.581799852 | 0.000341235 | 0.016369539 |
| DTD1 | 16.43432434 | 0.480330087 | 0.134387441 | 3.574218566 | 0.000351275 | 0.016807528 |
| TMEM87B | 119.5945301 | -0.214132806 | 0.059929899 | -3.573054682 | 0.000352841 | 0.016822232 |
| HSP90AA1 | 1416.539689 | 0.331270281 | 0.092724303 | 3.572637057 | 0.000353404 | 0.016822232 |
| SNX13 | 87.90062152 | -0.233859569 | 0.065477934 | -3.571578299 | 0.000354836 | 0.016846977 |
| METTL14 | 74.92388132 | -0.216257882 | 0.060594822 | -3.568916872 | 0.00035846 | 0.016949984 |
| TBK1 | 51.26018817 | -0.269983343 | 0.075654444 | -3.568638242 | 0.000358841 | 0.016949984 |
| SURF2 | 24.8681925 | 0.308339222 | 0.086551083 | 3.562511441 | 0.000367324 | 0.01730639 |
| JTB | 158.418141 | 0.257126451 | 0.072330134 | 3.554900792 | 0.000378122 | 0.017730129 |
| B4GALT3 | 64.73800129 | 0.387218911 | 0.108968065 | 3.553508189 | 0.000380129 | 0.017730129 |
| SAPCD2 | 30.13249261 | 0.569315428 | 0.160213099 | 3.553488634 | 0.000380158 | 0.017730129 |
| PLAAT5 | 15.59990054 | -0.967671536 | 0.272214575 | -3.554811629 | 0.00037825 | 0.017730129 |
| MYB | 131.3272909 | -0.656604188 | 0.18493691 | -3.550422608 | 0.000384613 | 0.017847792 |
| MAST1 | 19.41973046 | -1.40272476 | 0.395071148 | -3.550562391 | 0.000384409 | 0.017847792 |
| FBXL20 | 135.8439897 | 0.654342386 | 0.184496334 | 3.546641667 | 0.000390175 | 0.018060502 |
| UQCR11 | 209.8786858 | 0.251381989 | 0.070894668 | 3.545851865 | 0.000391346 | 0.018069429 |
| AC055845.1 | 3.431447301 | -0.875815312 | 0.247070779 | -3.544795203 | 0.000392918 | 0.018096776 |
| OST4 | 208.5014832 | 0.286035501 | 0.080767576 | 3.541464478 | 0.000397912 | 0.018211125 |
| ABCB9 | 38.30753173 | -0.451177219 | 0.127409136 | -3.541168488 | 0.000398359 | 0.018211125 |
| AC093249.3 | 2.946002212 | -1.803046535 | 0.509024486 | -3.54216071 | 0.000396864 | 0.018211125 |
| PSMB9 | 36.58372415 | 0.635073454 | 0.179639745 | 3.535261382 | 0.000407372 | 0.01848588 |
| H2AX | 43.39160352 | 0.394213555 | 0.11150413 | 3.535416631 | 0.000407133 | 0.01848588 |
| HNRNPA1 | 412.124858 | 0.24363687 | 0.068911493 | 3.535504158 | 0.000406998 | 0.01848588 |
| WNT11 | 8.586860129 | -1.00356267 | 0.284013231 | -3.533506759 | 0.000410086 | 0.018563409 |
| RUFY2 | 78.29324528 | -0.23775759 | 0.067363726 | -3.529460216 | 0.000416408 | 0.018803533 |
| AL596247.1 | 3.398372278 | -0.88865127 | 0.251845548 | -3.528556598 | 0.000417833 | 0.018821829 |
| CHIT1 | 25.45435109 | 1.22297446 | 0.347097822 | 3.523428793 | 0.000426002 | 0.018913035 |
| CFAP69 | 36.75905342 | -0.506744949 | 0.143752377 | -3.525123973 | 0.000423285 | 0.018913035 |
| NOMO1 | 17.02383253 | 0.510543568 | 0.144805172 | 3.525727437 | 0.000422321 | 0.018913035 |
| NDUFB7 | 153.3179959 | 0.247921863 | 0.070355262 | 3.523856746 | 0.000425314 | 0.018913035 |
| RPL23AP2 | 16.01822656 | 0.585318641 | 0.165981754 | 3.526403515 | 0.000421245 | 0.018913035 |
| NRIP1 | 375.7797581 | -0.436250071 | 0.123769561 | -3.524695945 | 0.000423969 | 0.018913035 |

| | | | | | | |
|---|---|---|---|---|---|---|
| NCF4 | 12.1923059 | 0.562360186 | 0.159652381 | 3.522403993 | 0.000427652 | 0.018940775 |
| GPATCH8 | 149.4362425 | -0.232743717 | 0.066087278 | -3.521762773 | 0.000428688 | 0.018941222 |
| AC015912.3 | 2.447785499 | 0.952989587 | 0.271034624 | 3.516117508 | 0.000437907 | 0.019302403 |
| PRC1 | 8.441181555 | 0.612952333 | 0.174503018 | 3.512560075 | 0.000443812 | 0.019516089 |
| TRIP12 | 326.4709675 | -0.158549212 | 0.045166959 | -3.510291909 | 0.000447615 | 0.019636583 |
| MIR6797 | 2.911818373 | 1.037191657 | 0.295581477 | 3.508987326 | 0.000449816 | 0.019686393 |
| PCDH19 | 11.1996762 | -0.753778404 | 0.214869092 | -3.50808204 | 0.00045135 | 0.019706809 |
| EIF2S2 | 101.8196457 | 0.27741217 | 0.079204752 | 3.502468763 | 0.000460968 | 0.02007928 |
| MMP1 | 2.870320127 | 1.547046829 | 0.441984057 | 3.500232205 | 0.000464853 | 0.020200872 |
| TASOR2 | 200.6959313 | -0.201937647 | 0.057723872 | -3.498338536 | 0.000468166 | 0.020297104 |
| LINC00882 | 7.231218969 | -0.607882717 | 0.173910644 | -3.495373842 | 0.000473398 | 0.020442485 |
| ZNF92 | 36.54418979 | -0.615215744 | 0.176018083 | -3.495184889 | 0.000473733 | 0.020442485 |
| PPP1R14B | 79.11583258 | 0.384012591 | 0.109988266 | 3.491395983 | 0.000480504 | 0.020655601 |
| RAB5C | 231.1801484 | -0.241722081 | 0.069238117 | -3.491170632 | 0.000480909 | 0.020655601 |
| RBM27 | 82.22059725 | -0.195764841 | 0.056100377 | -3.489545918 | 0.000483842 | 0.020733356 |
| ARPC4 | 106.385593 | 0.251163575 | 0.071989593 | 3.488887268 | 0.000485036 | 0.020736397 |
| SPTLC3 | 16.18670661 | -0.47165561 | 0.135248174 | -3.487334397 | 0.000487861 | 0.020809016 |
| RPL37A | 2229.268952 | 0.277020329 | 0.079528483 | 3.483284482 | 0.000495302 | 0.021077714 |
| SLC16A9 | 12.98154905 | -0.629836693 | 0.180853068 | -3.482587818 | 0.000496592 | 0.021084054 |
| MRPL9 | 59.28213942 | 0.260458328 | 0.074821625 | 3.481056841 | 0.000499439 | 0.02110789 |
| CSKMT | 51.73187185 | 0.50985458 | 0.146457921 | 3.481235948 | 0.000499106 | 0.02110789 |
| BCL2L12 | 18.79313439 | 0.419441092 | 0.120548964 | 3.4794251 | 0.000502491 | 0.021188363 |
| TMA7 | 69.41176557 | 0.346478653 | 0.099620274 | 3.477993369 | 0.000505182 | 0.021253335 |
| TMEM106B | 175.1076201 | -0.227657106 | 0.065562487 | -3.472368376 | 0.000515888 | 0.021303969 |
| MKLN1 | 172.1630942 | -0.193502935 | 0.055683315 | -3.475061336 | 0.000510737 | 0.021303969 |
| RPL30 | 515.0432607 | 0.330174488 | 0.095050921 | 3.473659017 | 0.000513413 | 0.021303969 |
| AKR1C2 | 7.824634217 | 1.11409532 | 0.320875867 | 3.472044594 | 0.000516511 | 0.021303969 |
| BLOC1S1 | 95.95753951 | 0.204840708 | 0.058998291 | 3.471976974 | 0.000516641 | 0.021303969 |
| TBC1D15 | 86.27345323 | -0.233327206 | 0.067184521 | -3.472931005 | 0.000514808 | 0.021303969 |
| AC013652.1 | 11.71170394 | -1.009450423 | 0.290747775 | -3.471911086 | 0.000516767 | 0.021303969 |
| CYP4F8 | 13.24560049 | -1.607973934 | 0.462590029 | -3.476023768 | 0.000508907 | 0.021303969 |
| SNORA69 | 5.720926453 | -0.691754242 | 0.198966035 | -3.476745371 | 0.00050754 | 0.021303969 |
| PTDSS1 | 50.26646972 | 0.310455028 | 0.089588422 | 3.465347655 | 0.000529546 | 0.021780594 |
| TRIM3 | 32.4838242 | -0.431893925 | 0.124653029 | -3.4647688 | 0.000530687 | 0.021780594 |
| PNMA2 | 9.701856217 | -0.719041887 | 0.207793923 | -3.460360522 | 0.000539453 | 0.022076387 |

| | | | | | | |
|---|---|---|---|---|---|---|
| NFAM1 | 8.907913148 | 0.55889958 | 0.161534225 | 3.459945272 | 0.000540285 | 0.022076387 |
| HIGD2A | 46.15375185 | 0.329367441 | 0.09527758 | 3.456924933 | 0.000546377 | 0.022276021 |
| TBCA | 155.3454252 | 0.27261613 | 0.078884852 | 3.455874279 | 0.000548511 | 0.022313769 |
| PLP2 | 106.177501 | 0.319370359 | 0.092436944 | 3.455007751 | 0.000550277 | 0.022336409 |
| STUM | 10.63397101 | -0.980696546 | 0.283992257 | -3.453250997 | 0.000553873 | 0.022387534 |
| NCOA1 | 158.7201456 | -0.192690894 | 0.055838562 | -3.450857021 | 0.00055881 | 0.022387534 |
| PCM1 | 440.8138524 | -0.295659205 | 0.085649774 | -3.451955453 | 0.00055654 | 0.022387534 |
| NUP62 | 84.88966784 | 0.230957921 | 0.066921383 | 3.451182723 | 0.000558136 | 0.022387534 |
| MIS18A | 30.55118541 | 0.37600203 | 0.108946715 | 3.451247078 | 0.000558002 | 0.022387534 |
| CSTB | 162.7171841 | 0.440293571 | 0.127548316 | 3.451974799 | 0.0005565 | 0.022387534 |
| NKX6-1 | 2.136153322 | -1.194928098 | 0.346709527 | -3.446481871 | 0.000567937 | 0.022703945 |
| TEAD1 | 173.9682864 | -0.250056031 | 0.072597328 | -3.444424734 | 0.000572276 | 0.022828001 |
| CEP126 | 68.53137642 | -0.450744113 | 0.130940948 | -3.442346505 | 0.000576691 | 0.022954541 |
| GPX1 | 172.8288023 | 0.382644906 | 0.111233175 | 3.440025036 | 0.00058166 | 0.02310255 |
| ETFBKMT | 14.05132818 | -0.412279799 | 0.119977044 | -3.43632235 | 0.000589669 | 0.023370376 |
| ICA1L | 25.05582714 | -0.40619679 | 0.118425897 | -3.42996592 | 0.000603657 | 0.023813823 |
| LINC00472 | 172.2657703 | -0.601073864 | 0.175296011 | -3.428907821 | 0.000606015 | 0.023813823 |
| SEC61B | 118.0350842 | 0.349257017 | 0.101809622 | 3.430491256 | 0.000602489 | 0.023813823 |
| ATRN | 65.72805821 | -0.271089203 | 0.079058059 | -3.428988856 | 0.000605834 | 0.023813823 |
| PLP1 | 7.071570975 | -0.778574091 | 0.227125468 | -3.427947112 | 0.000608164 | 0.023847515 |
| POLN | 15.79054489 | -0.620610683 | 0.181114425 | -3.426622061 | 0.000611139 | 0.023913404 |
| KCNJ3 | 50.93126072 | -1.648604919 | 0.481218131 | -3.425899427 | 0.000612767 | 0.023926423 |
| GIMAP4 | 48.08526372 | 0.449252948 | 0.131201574 | 3.424142983 | 0.000616742 | 0.023936048 |
| VPS37A | 45.44177747 | -0.31237415 | 0.091228838 | -3.424072429 | 0.000616902 | 0.023936048 |
| MORF4L1 | 64.10443942 | 0.306228459 | 0.089431258 | 3.424177032 | 0.000616664 | 0.023936048 |
| SQOR | 56.80223412 | 0.321055219 | 0.093798177 | 3.422830055 | 0.000619728 | 0.023945101 |
| PIP4K2B | 96.49226529 | 0.615134455 | 0.179694783 | 3.423218225 | 0.000618844 | 0.023945101 |
| SNRPE | 120.8050403 | 0.305979977 | 0.089437055 | 3.421176797 | 0.000623508 | 0.023990533 |
| RN7SL832P | 9.209340393 | -0.499850372 | 0.146153008 | -3.420048469 | 0.0006261 | 0.023990533 |
| RN7SL123P | 1.41941533 | -1.383428695 | 0.404465335 | -3.420388785 | 0.000625317 | 0.023990533 |
| PLBD1-AS1 | 6.692489646 | -0.643687779 | 0.188172983 | -3.420723683 | 0.000624548 | 0.023990533 |
| AL590762.5 | 4.052104604 | -0.690391522 | 0.201945629 | -3.418699995 | 0.000629211 | 0.024059814 |
| CNNM4 | 43.73515562 | -0.452459947 | 0.132531742 | -3.413974183 | 0.000640226 | 0.024380086 |
| CALM3 | 174.1041968 | 0.242287969 | 0.070966691 | 3.414108282 | 0.000639911 | 0.024380086 |
| PRDX4 | 55.20585967 | 0.32345482 | 0.094779433 | 3.412711067 | 0.000643201 | 0.02444296 |

| | | | | | | |
|---|---|---|---|---|---|---|
| PGD | 52.31512091 | 0.400729982 | 0.11748289 | 3.410964627 | 0.000647335 | 0.024471014 |
| ADA | 14.48141444 | 0.509363589 | 0.149341921 | 3.410720749 | 0.000647914 | 0.024471014 |
| MORF4L2-AS1 | 1.655079693 | -1.043219462 | 0.305793235 | -3.411519098 | 0.00064602 | 0.024471014 |
| CYS1 | 20.44176439 | -0.654835565 | 0.192074044 | -3.409287125 | 0.000651329 | 0.024539013 |
| AL354861.2 | 5.143941296 | -0.538935811 | 0.15809898 | -3.408850652 | 0.000652372 | 0.024539013 |
| DNAH14 | 70.86143475 | -0.412780113 | 0.12112306 | -3.40793992 | 0.000654553 | 0.024571017 |
| RAB1B | 61.67523984 | 0.391766133 | 0.114989161 | 3.406983153 | 0.000656852 | 0.024607294 |
| HOXA5 | 21.88751561 | -0.618990346 | 0.181844587 | -3.40395255 | 0.000664183 | 0.024831567 |
| CDC34 | 51.4162594 | 0.285239348 | 0.083862301 | 3.40128215 | 0.000670706 | 0.024974319 |
| SUN2 | 85.02822438 | 0.277136859 | 0.081473498 | 3.401558389 | 0.000670028 | 0.024974319 |
| CLTC | 553.4881821 | 0.417058189 | 0.122749369 | 3.397640184 | 0.000679698 | 0.025258217 |
| ARID1A | 310.2752836 | -0.15111305 | 0.044535375 | -3.39310156 | 0.00069106 | 0.025616768 |
| FDPS | 140.6845747 | 0.310327447 | 0.091484452 | 3.392133199 | 0.000693507 | 0.025616768 |
| TSHZ1 | 88.24893704 | -0.230173133 | 0.067851751 | -3.392294667 | 0.000693099 | 0.025616768 |
| KIAA1109 | 290.3992179 | -0.223051626 | 0.06586442 | -3.386526831 | 0.000707834 | 0.026041788 |
| PDCD5 | 53.75195866 | 0.336676314 | 0.099408923 | 3.386781618 | 0.000707177 | 0.026041788 |
| AIF1L | 43.18436479 | 0.485862259 | 0.143498363 | 3.385838342 | 0.000709612 | 0.026055306 |
| SH3BP2 | 104.2971581 | 0.230695687 | 0.068149814 | 3.385125677 | 0.000711457 | 0.026071219 |
| TYROBP | 112.396634 | 0.502508135 | 0.148551143 | 3.382728162 | 0.000717696 | 0.026247789 |
| ST14 | 88.74418336 | 0.397608672 | 0.117578307 | 3.381649903 | 0.000720519 | 0.026273839 |
| FSIP1 | 30.32672015 | -0.984572255 | 0.291175554 | -3.3813699 | 0.000721254 | 0.026273839 |
| LIMK1 | 49.61115511 | 0.337505142 | 0.099896196 | 3.378558498 | 0.000728669 | 0.026491718 |
| CDC42SE1 | 94.93058792 | 0.277755906 | 0.082257473 | 3.376664725 | 0.000733704 | 0.026549177 |
| H2AZ1 | 196.7275228 | 0.307573466 | 0.091125631 | 3.375268441 | 0.000737437 | 0.026549177 |
| EMB | 37.78210587 | 0.470541986 | 0.139389333 | 3.375738833 | 0.000736178 | 0.026549177 |
| OARD1 | 57.01014179 | -0.262273291 | 0.077687726 | -3.375993923 | 0.000735495 | 0.026549177 |
| PSD3 | 89.90170734 | -0.614454141 | 0.182029385 | -3.375576653 | 0.000736612 | 0.026549177 |
| TBX3 | 239.9867503 | -0.598309924 | 0.177347949 | -3.373650086 | 0.000741786 | 0.026653784 |
| IRF8 | 19.87398329 | 0.804079416 | 0.238548524 | 3.370716381 | 0.00074973 | 0.02688692 |
| DNAH6 | 6.251002762 | -0.642669283 | 0.190698226 | -3.37008528 | 0.000751449 | 0.026888925 |
| AL353795.3 | 23.61040968 | -0.426895871 | 0.126689324 | -3.369627818 | 0.000752698 | 0.026888925 |
| AC017048.3 | 1.899471327 | -1.053392229 | 0.312731432 | -3.368360588 | 0.000756166 | 0.02691803 |
| CNPY2 | 99.96354475 | 0.207103112 | 0.061496267 | 3.367734706 | 0.000757885 | 0.02691803 |
| FXR2 | 44.36685418 | 0.27917547 | 0.082890676 | 3.367996046 | 0.000757167 | 0.02691803 |
| RIC8B | 35.66399938 | -0.333164983 | 0.098959598 | -3.366676799 | 0.000760798 | 0.026953256 |

| | | | | | | |
|---|---|---|---|---|---|---|
| CDK2AP1 | 99.90812799 | 0.330746555 | 0.098251801 | 3.366315438 | 0.000761795 | 0.026953256 |
| UBP1 | 132.8455084 | -0.214920916 | 0.063855565 | -3.365735115 | 0.0007634 | 0.026958376 |
| SEC62 | 523.5891403 | 0.177523084 | 0.052772417 | 3.36393699 | 0.000768391 | 0.027031258 |
| RPL22L1 | 45.32933306 | 0.342127785 | 0.10169617 | 3.364215031 | 0.000767617 | 0.027031258 |
| SP6 | 13.05478521 | 0.868003832 | 0.258250693 | 3.361090042 | 0.000776355 | 0.027259511 |
| CHM | 50.89180774 | -0.238253561 | 0.070922831 | -3.35933519 | 0.000781302 | 0.027381165 |
| HHIPL2 | 3.552945565 | 1.187121892 | 0.353487297 | 3.358315569 | 0.00078419 | 0.02738964 |
| TPT1 | 1792.490753 | 0.300759443 | 0.089559642 | 3.358202829 | 0.00078451 | 0.02738964 |
| COX7C | 441.2457525 | 0.225666068 | 0.067273961 | 3.354434084 | 0.000795275 | 0.027660892 |
| HERC1 | 209.1348155 | -0.2369009 | 0.070619064 | -3.354630991 | 0.000794709 | 0.027660892 |
| DUBR | 21.34248226 | -0.497909158 | 0.148479104 | -3.353395483 | 0.000798266 | 0.027712719 |
| EXTL2 | 48.4148132 | -0.258816417 | 0.077470048 | -3.340857826 | 0.0008352 | 0.028865818 |
| AL391058.1 | 3.07591528 | -0.937739402 | 0.280645739 | -3.341363403 | 0.00083368 | 0.028865818 |
| CCT8 | 58.27876243 | 0.333453396 | 0.099820337 | 3.340535665 | 0.000836169 | 0.028865818 |
| KIF20A | 4.672165848 | 0.776482462 | 0.232671734 | 3.33724449 | 0.000846135 | 0.029155343 |
| SLC10A7 | 30.3409435 | -0.331692355 | 0.099419564 | -3.336288574 | 0.00084905 | 0.02919052 |
| NDUFAB1 | 99.93818027 | 0.290721654 | 0.087156724 | 3.335619328 | 0.000851096 | 0.02919052 |
| HSD17B10 | 72.49596941 | 0.248317166 | 0.074449934 | 3.335357756 | 0.000851897 | 0.02919052 |
| LIN7C | 61.86066648 | -0.259644277 | 0.077858761 | -3.334811332 | 0.000853573 | 0.029193774 |
| OTULINL | 21.6151152 | 0.44764401 | 0.134310171 | 3.332912223 | 0.000859421 | 0.029339441 |
| COMMD6 | 80.48232925 | 0.311663774 | 0.093545028 | 3.331697901 | 0.000863179 | 0.029413383 |
| C15orf48 | 60.27654815 | -0.842054725 | 0.252794823 | -3.330980895 | 0.000865405 | 0.02943494 |
| GK5 | 114.7097485 | -0.249208258 | 0.074853574 | -3.329276669 | 0.000870719 | 0.029561218 |
| LMTK2 | 24.67585625 | -0.312675374 | 0.093990692 | -3.326663177 | 0.000878925 | 0.029785089 |
| C1orf162 | 20.32866738 | 0.527115555 | 0.158526747 | 3.325089081 | 0.000883903 | 0.029844246 |
| UBE2D2 | 130.7357973 | 0.169993681 | 0.051120328 | 3.325363627 | 0.000883033 | 0.029844246 |
| COX5B | 151.7180877 | 0.230487141 | 0.069347002 | 3.323678533 | 0.000888385 | 0.029940857 |
| EXOSC5 | 25.34272103 | 0.31500515 | 0.094860334 | 3.320725717 | 0.000897837 | 0.030204292 |
| FAM174C | 80.38340522 | 0.244850339 | 0.073780489 | 3.318632651 | 0.000904593 | 0.030376248 |
| KDM4A | 93.96609402 | -0.223200669 | 0.067390568 | -3.312046142 | 0.000926163 | 0.031041088 |
| SNX5 | 73.62117365 | 0.26265259 | 0.079313712 | 3.311565988 | 0.000927754 | 0.031041088 |
| SNRPB2 | 104.9167237 | 0.272089519 | 0.082180127 | 3.31089193 | 0.000929991 | 0.031059687 |
| AC004151.1 | 48.7900012 | 0.496620895 | 0.150053 | 3.309636556 | 0.000934172 | 0.031142996 |
| TWF2 | 42.75108586 | 0.286728288 | 0.086681636 | 3.30783199 | 0.000940212 | 0.031287883 |
| RNU6-681P | 1.789484564 | 1.12597673 | 0.340667941 | 3.305203089 | 0.000949076 | 0.031459933 |

| | | | | | | |
|---|---|---|---|---|---|---|
| DNPH1 | 44.3873226 | 0.33711874 | 0.102043998 | 3.303660656 | 0.000954313 | 0.031459933 |
| EFHC1 | 53.81102734 | -0.331355542 | 0.100311008 | -3.303281963 | 0.000955602 | 0.031459933 |
| NUP98 | 50.89804462 | -0.212886411 | 0.06442328 | -3.30449508 | 0.000951476 | 0.031459933 |
| TP53BP1 | 155.535437 | -0.207836057 | 0.062872354 | -3.305682762 | 0.000947453 | 0.031459933 |
| PALB2 | 23.65608149 | -0.282986077 | 0.085648503 | -3.304039985 | 0.000953022 | 0.031459933 |
| VNN2 | 6.069580893 | 0.845479868 | 0.256001591 | 3.302635213 | 0.000957809 | 0.031476466 |
| COL2A1 | 13.31186317 | 1.680713799 | 0.509349773 | 3.299724252 | 0.000967799 | 0.031748267 |
| TTC28 | 49.31175136 | -0.289069731 | 0.087638725 | -3.298424668 | 0.00097229 | 0.031839038 |
| ARMCX6 | 6.097894375 | -0.544397252 | 0.165098761 | -3.29740362 | 0.000975832 | 0.031898467 |
| NDUFB2 | 132.3660602 | 0.247001213 | 0.074957543 | 3.295214892 | 0.000983464 | 0.032034576 |
| NAT14 | 23.50127419 | 0.4149841 | 0.125918968 | 3.295644065 | 0.000981963 | 0.032034576 |
| CYTOR | 25.77367386 | 0.43520985 | 0.132220468 | 3.291546743 | 0.000996381 | 0.032131324 |
| CCDC149 | 50.84652823 | -0.299198307 | 0.090897321 | -3.291607556 | 0.000996165 | 0.032131324 |
| ZDHHC21 | 72.44302576 | -0.291828845 | 0.088639046 | -3.292328355 | 0.000993615 | 0.032131324 |
| PLD2 | 26.0579144 | 0.315586732 | 0.09584865 | 3.292552713 | 0.000992823 | 0.032131324 |
| JADE3 | 17.0185968 | -0.400067757 | 0.121549132 | -3.29140775 | 0.000996873 | 0.032131324 |
| MT-TG | 6852.986011 | -0.376543971 | 0.114396084 | -3.291580953 | 0.000996259 | 0.032131324 |
| NIBAN2 | 113.3888145 | 0.238396963 | 0.072461865 | 3.289964468 | 0.001002 | 0.03220321 |
| RNASE4 | 17.42876496 | -0.486930419 | 0.148012221 | -3.289798755 | 0.001002591 | 0.03220321 |
| AL158206.1 | 23.36328944 | -0.428576727 | 0.130329775 | -3.288402268 | 0.001007578 | 0.032307206 |
| LINC02515 | 2.298106749 | 1.263074241 | 0.384167223 | 3.287824062 | 0.001009649 | 0.032317522 |
| MFSD2A | 12.79746177 | 0.760632487 | 0.231497166 | 3.285709715 | 0.001017258 | 0.032336894 |
| MYSM1 | 175.9051568 | -0.270800556 | 0.082407584 | -3.286112051 | 0.001015806 | 0.032336894 |
| VSTM2A | 2.464401568 | -2.31487597 | 0.704354873 | -3.286519423 | 0.001014338 | 0.032336894 |
| PEX1 | 83.6818361 | -0.241556476 | 0.073511546 | -3.285966469 | 0.001016331 | 0.032336894 |
| ATP5MC2 | 388.7028249 | 0.24770487 | 0.075402293 | 3.285110556 | 0.001019424 | 0.03235006 |
| ACP1 | 59.85011774 | 0.226451662 | 0.068978094 | 3.282950429 | 0.001027267 | 0.032532934 |
| RBM20 | 24.00038023 | -1.363091304 | 0.415253158 | -3.282554938 | 0.001028709 | 0.032532934 |
| DYNLT1 | 33.14801901 | 0.397741553 | 0.121262149 | 3.280014064 | 0.001038019 | 0.032769337 |
| AC106037.1 | 1.898501287 | -1.003167067 | 0.305885718 | -3.279548564 | 0.001039733 | 0.032769337 |
| RPLP2 | 1317.07743 | 0.387293279 | 0.118202439 | 3.276525284 | 0.001050929 | 0.033009538 |
| AL138957.1 | 1.838526488 | -0.973519831 | 0.297077386 | -3.276990699 | 0.001049198 | 0.033009538 |
| H2BC20P | 21.44462842 | 0.574646315 | 0.175536416 | 3.273658703 | 0.001061648 | 0.03328959 |
| H2BC21 | 48.07725435 | 0.710641124 | 0.217117306 | 3.273074528 | 0.001063844 | 0.03330193 |
| DHODH | 12.83375136 | 0.373474039 | 0.114129001 | 3.272385068 | 0.001066442 | 0.03332677 |

| | | | | | | |
|---|---|---|---|---|---|---|
| ORAI2 | 83.27045945 | 0.399670269 | 0.12215462 | 3.271839145 | 0.001068503 | 0.033334782 |
| SPX | 2.910904339 | 1.035408115 | 0.316569612 | 3.270712275 | 0.00107277 | 0.033411448 |
| MMP24OS | 122.4853968 | 0.35250697 | 0.107801055 | 3.269976984 | 0.001075562 | 0.033442022 |
| C20orf96 | 39.26792207 | -0.394819727 | 0.120793491 | -3.268551346 | 0.001080996 | 0.033554467 |
| AC011815.2 | 3.011134599 | -0.91054926 | 0.278660097 | -3.267598301 | 0.001084642 | 0.033611161 |
| FAM193A | 96.56769778 | -0.197828832 | 0.060562128 | -3.266543611 | 0.00108869 | 0.033680105 |
| OLA1 | 54.14719653 | 0.371262869 | 0.113726802 | 3.264515171 | 0.001096516 | 0.033705963 |
| NEMF | 270.0654946 | -0.215794143 | 0.066093543 | -3.264980715 | 0.001094715 | 0.033705963 |
| SRRM2 | 1165.266362 | -0.226633757 | 0.069422029 | -3.264579873 | 0.001096265 | 0.033705963 |
| MT2A | 221.6202293 | 0.515463204 | 0.157902731 | 3.264435007 | 0.001096826 | 0.033705963 |
| MIR335 | 129.5012266 | -0.50961874 | 0.156170059 | -3.263229471 | 0.001101503 | 0.033761237 |
| DHPS | 52.38280479 | -0.282685959 | 0.08663298 | -3.263029385 | 0.001102281 | 0.033761237 |
| ISY1 | 42.35079031 | 0.242805043 | 0.074438776 | 3.261808653 | 0.001107038 | 0.033850815 |
| PSMG1 | 33.6333756 | 0.304232702 | 0.093380702 | 3.257982584 | 0.001122073 | 0.034253826 |
| AC023590.1 | 1.743320755 | 1.068258289 | 0.327960568 | 3.25727662 | 0.001124868 | 0.034282473 |
| EMX1 | 2.509201669 | -1.757210232 | 0.53981816 | -3.255189178 | 0.001133169 | 0.034421859 |
| ZCCHC14 | 47.00805103 | -0.314043094 | 0.096474549 | -3.255191117 | 0.001133161 | 0.034421859 |
| AKIRIN1 | 63.97820077 | 0.220343379 | 0.067717813 | 3.253846628 | 0.001138537 | 0.034489497 |
| MIR320E | 3.990222546 | -0.893061469 | 0.274475769 | -3.253698754 | 0.00113913 | 0.034489497 |
| SEMA7A | 3.884308538 | 0.724750571 | 0.22291938 | 3.251177935 | 0.001149279 | 0.034739828 |
| AL162741.1 | 1.418104157 | -1.132580236 | 0.348765508 | -3.247397495 | 0.001164656 | 0.03474965 |
| PTGER3 | 42.84407937 | -0.560844735 | 0.17269248 | -3.247650019 | 0.001163623 | 0.03474965 |
| HIPK1 | 159.5886107 | -0.204412984 | 0.062921372 | -3.248705146 | 0.001159316 | 0.03474965 |
| NDUFB3 | 38.44330811 | 0.314952111 | 0.096963696 | 3.24814465 | 0.001161602 | 0.03474965 |
| UBTD2 | 61.76932095 | -0.228019311 | 0.070168461 | -3.249598269 | 0.001155681 | 0.03474965 |
| CORO2A | 53.75965372 | 0.407038213 | 0.125314288 | 3.248138889 | 0.001161626 | 0.03474965 |
| IL32 | 35.14230167 | 0.583638031 | 0.179703225 | 3.247788305 | 0.001163058 | 0.03474965 |
| STAC2 | 50.90770126 | 0.878662271 | 0.270557583 | 3.247598016 | 0.001163836 | 0.03474965 |
| FGD5-AS1 | 186.1987646 | -0.230854255 | 0.071139073 | -3.245111928 | 0.001174045 | 0.034915225 |
| SNORD90 | 20.20355919 | -0.455165015 | 0.140245459 | -3.245488434 | 0.001172493 | 0.034915225 |
| TAX1BP3 | 112.3210869 | 0.290020866 | 0.089396443 | 3.244210345 | 0.001177767 | 0.034915225 |
| AC091153.1 | 1.46264036 | 0.984113371 | 0.303334365 | 3.244318756 | 0.001177319 | 0.034915225 |
| CAB39L | 53.46662355 | -0.402861744 | 0.124301926 | -3.240993591 | 0.001191139 | 0.035198623 |
| GRB7 | 282.8385595 | 0.959894348 | 0.296163228 | 3.24109902 | 0.001190698 | 0.035198623 |
| LINC01881 | 10.23321674 | 0.438520773 | 0.13534479 | 3.240026982 | 0.001195184 | 0.035208303 |

| | | | | | | |
|---|---|---|---|---|---|---|
| CEP290 | 191.3899442 | -0.234636123 | 0.07241846 | -3.240004321 | 0.001195279 | 0.035208303 |
| AC092620.1 | 17.63951691 | -0.524243088 | 0.161844043 | -3.239186798 | 0.00119871 | 0.035253156 |
| SELENOS | 115.0121929 | 0.217327352 | 0.067107906 | 3.238476156 | 0.001201701 | 0.035284912 |
| SDC1 | 293.2932132 | 0.603166414 | 0.186353267 | 3.236682794 | 0.001209277 | 0.035322046 |
| LARP1B | 85.9095386 | -0.229154545 | 0.070803962 | -3.236465001 | 0.001210201 | 0.035322046 |
| SNORA38 | 45.88293453 | -0.729704625 | 0.225476673 | -3.236275471 | 0.001211005 | 0.035322046 |
| RDX | 156.4074066 | 0.231174472 | 0.071440181 | 3.235916647 | 0.001212528 | 0.035322046 |
| GDPD1 | 27.18859145 | 0.588051351 | 0.181704223 | 3.236310857 | 0.001210854 | 0.035322046 |
| HNRNPL | 69.47725981 | 0.268389225 | 0.082980943 | 3.234347739 | 0.001219209 | 0.035460745 |
| POLR2F | 34.25706718 | 0.317659984 | 0.098258265 | 3.232908546 | 0.001225368 | 0.035583834 |
| SENP6 | 267.6285306 | -0.189392383 | 0.05861855 | -3.230929196 | 0.001233885 | 0.035718844 |
| COLCA1 | 4.132748668 | 0.796987561 | 0.246661263 | 3.231101436 | 0.001233142 | 0.035718844 |
| LAPTM5 | 204.3759554 | 0.481179299 | 0.149086507 | 3.227517433 | 0.001248694 | 0.036090148 |
| MT-CYB | 3124.863848 | 0.39424218 | 0.122166972 | 3.227076616 | 0.00125062 | 0.036090148 |
| LINC02210 | 26.84168494 | -0.389980891 | 0.120958945 | -3.224076479 | 0.001263796 | 0.036413504 |
| POTEE | 2.400356707 | -1.435031065 | 0.445406699 | -3.221844366 | 0.001273683 | 0.036584217 |
| TTC37 | 167.4109226 | -0.167966974 | 0.052126675 | -3.222284464 | 0.001271728 | 0.036584217 |
| ZFYVE9 | 48.97807159 | -0.30671058 | 0.095245029 | -3.220226655 | 0.001280893 | 0.036705606 |
| SIT1 | 7.589052842 | 0.838645843 | 0.260484551 | 3.219560773 | 0.001283871 | 0.036705606 |
| RALGAPA1 | 78.89250019 | -0.244261335 | 0.075862701 | -3.219781693 | 0.001282883 | 0.036705606 |
| POLDIP3 | 31.0975072 | 0.270889609 | 0.08416696 | 3.218479175 | 0.001288723 | 0.036787373 |
| ABCC4 | 26.43085957 | 0.599511148 | 0.186400787 | 3.216247946 | 0.001298786 | 0.037017396 |
| CCDC171 | 32.05170182 | -0.351747104 | 0.109398799 | -3.21527391 | 0.001303201 | 0.037035515 |
| MLLT6 | 369.6034078 | 0.565725886 | 0.175952298 | 3.215223059 | 0.001303432 | 0.037035515 |
| MYT1L | 2.689196684 | -1.064958206 | 0.331396075 | -3.213551053 | 0.001311045 | 0.037096922 |
| CMYA5 | 147.4281565 | -0.482453916 | 0.15013022 | -3.213569634 | 0.00131096 | 0.037096922 |
| RPL21P120 | 1.816699237 | 0.633790753 | 0.197232139 | 3.213425338 | 0.001311619 | 0.037096922 |
| TDG | 31.82917569 | 0.239839567 | 0.074653194 | 3.21271675 | 0.001314859 | 0.037131697 |
| CDSN | 5.319013494 | -1.378614683 | 0.4292274 | -3.211851531 | 0.001318825 | 0.037186842 |
| ZNF26 | 59.7202135 | -0.299688708 | 0.0934216 | -3.207916662 | 0.001337002 | 0.037641915 |
| TET1 | 27.04532159 | -0.413251341 | 0.128877003 | -3.206556109 | 0.001343341 | 0.037762809 |
| GTF2H1 | 52.72010773 | -0.21579416 | 0.067307323 | -3.206102254 | 0.001345462 | 0.037764942 |
| PAQR3 | 27.71595749 | -0.378672608 | 0.118138008 | -3.205341048 | 0.001349025 | 0.03780751 |
| SPP1 | 214.1540844 | 0.736256452 | 0.230009793 | 3.200978716 | 0.001369616 | 0.038326435 |
| MAPT | 184.7284445 | -0.829528848 | 0.259310583 | -3.198977999 | 0.001379157 | 0.038535022 |

| | | | | | | |
|---|---|---|---|---|---|---|
| PQBP1 | 51.17157772 | 0.238874546 | 0.074691956 | 3.198129483 | 0.001383222 | 0.03859021 |
| CORO6 | 13.23294262 | -0.965338812 | 0.302003854 | -3.196445346 | 0.001391322 | 0.038757651 |
| AC103702.1 | 6.586437095 | -0.898806147 | 0.281246975 | -3.195789565 | 0.001394488 | 0.038787341 |
| MED14 | 59.02198782 | -0.203523139 | 0.063697665 | -3.195142832 | 0.001397617 | 0.03881591 |
| CEP192 | 50.07997222 | -0.27364323 | 0.085695159 | -3.193216892 | 0.001406972 | 0.039017073 |
| GADD45GIP1 | 77.08432266 | 0.228123952 | 0.071454466 | 3.192577967 | 0.001410089 | 0.03904487 |
| VKORC1 | 141.0344558 | 0.205748213 | 0.064469789 | 3.191389577 | 0.001415902 | 0.039108331 |
| BCL2 | 152.9496732 | -0.516120107 | 0.161730064 | -3.191244058 | 0.001416616 | 0.039108331 |
| LATS1 | 88.53332441 | -0.21131282 | 0.066248936 | -3.189678705 | 0.00142431 | 0.039210251 |
| ESCO1 | 92.52912172 | -0.210330192 | 0.06594189 | -3.189629399 | 0.001424553 | 0.039210251 |
| RNU6-439P | 6.509125794 | -0.603415836 | 0.189253251 | -3.188404073 | 0.001430605 | 0.039259789 |
| ADGRE5 | 25.80233069 | 0.453510422 | 0.142221505 | 3.188761226 | 0.001428838 | 0.039259789 |
| CCNL2 | 312.9224018 | -0.223214937 | 0.070065188 | -3.18581802 | 0.001443454 | 0.039436605 |
| AC093838.1 | 5.028182201 | -1.216321397 | 0.381710509 | -3.186502252 | 0.001440044 | 0.039436605 |
| NSUN6 | 35.18241014 | -0.281861957 | 0.088471628 | -3.185902223 | 0.001443034 | 0.039436605 |
| AC124068.2 | 3.642168973 | -0.84809751 | 0.266378503 | -3.183806125 | 0.001453523 | 0.03965306 |
| COX6B2 | 3.590720263 | -1.317383266 | 0.414053977 | -3.181670359 | 0.001464284 | 0.039887695 |
| NBPF1 | 62.80975272 | -0.357259488 | 0.112429642 | -3.177627205 | 0.001484855 | 0.039905719 |
| MANEAL | 14.1354857 | 0.627664546 | 0.197453469 | 3.17879726 | 0.001478875 | 0.039905719 |
| HDGF | 488.7198256 | 0.206526358 | 0.065000667 | 3.177295996 | 0.001486552 | 0.039905719 |
| CD28 | 7.206311881 | 0.749242365 | 0.235575112 | 3.180481833 | 0.001470304 | 0.039905719 |
| ASB6 | 20.28764759 | 0.336614686 | 0.105909631 | 3.178319869 | 0.001481312 | 0.039905719 |
| C12orf57 | 219.7138024 | 0.247541822 | 0.077899735 | 3.177697887 | 0.001484493 | 0.039905719 |
| ELOB | 186.339202 | 0.235453773 | 0.074093333 | 3.177799716 | 0.001483972 | 0.039905719 |
| AC018628.2 | 13.2376744 | 0.541786372 | 0.17035024 | 3.180426232 | 0.001470586 | 0.039905719 |
| MYL12A | 226.5323027 | 0.264795898 | 0.083321721 | 3.177993623 | 0.00148298 | 0.039905719 |
| PITPNB | 63.57624504 | 0.265332542 | 0.08346167 | 3.179094572 | 0.001477359 | 0.039905719 |
| AC004540.1 | 11.9602338 | -0.564503458 | 0.177753451 | -3.175766521 | 0.001494412 | 0.040000421 |
| SOCS1 | 5.633942904 | 0.781411034 | 0.24605221 | 3.175793595 | 0.001494272 | 0.040000421 |
| FCER1G | 76.22913455 | 0.418982592 | 0.131965267 | 3.174945964 | 0.001498644 | 0.040055654 |
| ENSA | 276.522049 | 0.23886628 | 0.075277732 | 3.173133328 | 0.001508033 | 0.040248343 |
| PFDN2 | 79.03140784 | 0.278204608 | 0.08770338 | 3.172108152 | 0.001513366 | 0.040274299 |
| SPIDR | 58.87759825 | -0.274598449 | 0.086560906 | -3.172314883 | 0.001512289 | 0.040274299 |
| CAP2 | 42.07847217 | -0.535727904 | 0.168938265 | -3.171146003 | 0.001518388 | 0.040334043 |
| SCART1 | 3.662985875 | -0.630117503 | 0.198734029 | -3.170657318 | 0.001520945 | 0.040334043 |

| | | | | | | |
|---|---|---|---|---|---|---|
| NDUFA6 | 110.8126417 | 0.249842191 | 0.078804014 | 3.170424666 | 0.001522163 | 0.040334043 |
| SOS2 | 119.9443457 | -0.228537731 | 0.072166465 | -3.16681343 | 0.001541192 | 0.040721413 |
| CD37 | 67.29973712 | 0.610441947 | 0.192743034 | 3.167128447 | 0.001539523 | 0.040721413 |
| RGS5 | 552.1947595 | -0.46197786 | 0.145968077 | -3.164923927 | 0.001551235 | 0.040872275 |
| RUNDC3B | 7.195351583 | -0.525424949 | 0.166037702 | -3.164491809 | 0.00155354 | 0.040872275 |
| XPO6 | 62.21982299 | 0.233180466 | 0.073681137 | 3.16472403 | 0.001552301 | 0.040872275 |
| DDX39B | 58.1237289 | -0.329391122 | 0.104124936 | -3.163422067 | 0.001559261 | 0.040906236 |
| STIM1 | 61.04701594 | -0.238711851 | 0.075452278 | -3.1637461 | 0.001557526 | 0.040906236 |
| PER3 | 62.91691394 | -0.395613423 | 0.125118275 | -3.161915591 | 0.00156735 | 0.04096625 |
| HADHA | 204.0155004 | 0.160676717 | 0.050843739 | 3.160206551 | 0.001576573 | 0.04096625 |
| ZNF862 | 20.9131519 | -0.31077123 | 0.09835453 | -3.159704288 | 0.001579293 | 0.04096625 |
| TUSC3 | 54.2842942 | -0.442163494 | 0.139931296 | -3.159861362 | 0.001578442 | 0.04096625 |
| KLRD1 | 12.45659363 | 0.542134873 | 0.171493638 | 3.16125356 | 0.001570917 | 0.04096625 |
| DTD2 | 26.57290797 | -0.318839028 | 0.100832291 | -3.162072621 | 0.001566505 | 0.04096625 |
| KLHDC1 | 18.58142008 | -0.386472598 | 0.122260855 | -3.161049375 | 0.001572018 | 0.04096625 |
| MAP2K2 | 109.243034 | 0.230604376 | 0.07295579 | 3.160878329 | 0.001572942 | 0.04096625 |
| AIF1 | 59.09930024 | 0.436793458 | 0.138370982 | 3.156683936 | 0.001595742 | 0.041334868 |
| GPC4 | 28.28320443 | 0.499501356 | 0.158278406 | 3.155840204 | 0.001600365 | 0.041396559 |
| TTC1 | 94.62970467 | 0.192111055 | 0.060929234 | 3.153019394 | 0.001615911 | 0.041740217 |
| AL157387.1 | 90.77038448 | -0.942603872 | 0.299095934 | -3.151510149 | 0.001624285 | 0.041897937 |
| MTR | 125.4368967 | -0.173481191 | 0.055097119 | -3.148643614 | 0.001640301 | 0.042110305 |
| RPL35 | 1126.517685 | 0.202864311 | 0.064426185 | 3.148786638 | 0.001639498 | 0.042110305 |
| AC092338.2 | 2.537472831 | -0.834056156 | 0.264942362 | -3.148066433 | 0.001643543 | 0.042110305 |
| SNHG16 | 100.9811116 | 0.363458473 | 0.115435352 | 3.148588941 | 0.001640608 | 0.042110305 |
| ZNF460 | 26.61573822 | -0.404907719 | 0.12862381 | -3.147999733 | 0.001643918 | 0.042110305 |
| KYNU | 31.24941783 | 0.771351653 | 0.245089074 | 3.147229867 | 0.001648253 | 0.042162867 |
| AHSA2P | 157.5323009 | -0.309298043 | 0.098307717 | -3.146223431 | 0.001653936 | 0.042249716 |
| AC098591.3 | 1.602089113 | 1.085960444 | 0.345441595 | 3.143687556 | 0.001668335 | 0.042558664 |
| TXNDC12 | 2.366450255 | -0.859698557 | 0.273718981 | -3.140807243 | 0.001684829 | 0.042920143 |
| OCIAD2 | 32.19820897 | 0.348614252 | 0.111033909 | 3.1397098 | 0.001691153 | 0.042962723 |
| CNOT6 | 70.4441937 | -0.258767737 | 0.082417193 | -3.139730034 | 0.001691036 | 0.042962723 |
| NSA2 | 96.12068562 | 0.310201614 | 0.098910851 | 3.136173753 | 0.001711678 | 0.043424415 |
| TCEAL4 | 297.6479768 | -0.404692302 | 0.129146646 | -3.133587397 | 0.001726835 | 0.043748852 |
| RANBP6 | 36.15912915 | -0.268595744 | 0.08585368 | -3.128529195 | 0.001756836 | 0.044301231 |
| DCLRE1A | 11.72407563 | -0.338975304 | 0.108357826 | -3.128295553 | 0.001758233 | 0.044301231 |

| | | | | | | |
|---|---|---|---|---|---|---|
| RPS15A | 191.8800132 | 0.214110864 | 0.068429218 | 3.128939209 | 0.001754386 | 0.044301231 |
| IGBP1 | 63.85549743 | 0.236228369 | 0.075499255 | 3.128883471 | 0.001754719 | 0.044301231 |
| YWHAH | 176.1109769 | 0.22532787 | 0.072046756 | 3.127522751 | 0.001762862 | 0.044357348 |
| CDC14A | 20.28528889 | -0.590392409 | 0.188825501 | -3.126656125 | 0.001768066 | 0.04442777 |
| AURKB | 4.103659271 | 0.804863371 | 0.257457764 | 3.126195766 | 0.001770836 | 0.044436922 |
| SNHG5 | 10092.84791 | -0.394920238 | 0.126362566 | -3.125294545 | 0.001776271 | 0.044452506 |
| LILRA4 | 5.955617458 | 1.019492738 | 0.326170809 | 3.125640646 | 0.001774182 | 0.044452506 |
| CHI3L1 | 62.9109805 | 0.634301566 | 0.20305545 | 3.123784984 | 0.001785409 | 0.044611327 |
| QTRT1 | 69.81096445 | -0.279365632 | 0.089441393 | -3.123449021 | 0.001787448 | 0.044611327 |
| TRMT10B | 36.6121861 | -0.29479091 | 0.09449877 | -3.119521146 | 0.001811453 | 0.044969673 |
| CCNB2 | 14.10448095 | 0.574054887 | 0.184018786 | 3.119545019 | 0.001811306 | 0.044969673 |
| ZNF44 | 64.59207113 | -0.3056507 | 0.097980481 | -3.119506012 | 0.001811546 | 0.044969673 |
| AF127936.1 | 1.699375532 | 0.822336906 | 0.263599755 | 3.119642142 | 0.001810709 | 0.044969673 |
| FOCAD | 36.70567742 | -0.239502329 | 0.076787129 | -3.119042629 | 0.001814397 | 0.04498 |
| SLF2 | 123.4274235 | -0.217007569 | 0.069615387 | -3.117235679 | 0.001825556 | 0.045075117 |
| MACROD1 | 35.36811873 | -0.358346738 | 0.114934224 | -3.117841877 | 0.001821805 | 0.045075117 |
| RSL24D1 | 68.64206908 | 0.254515808 | 0.081640755 | 3.117509233 | 0.001823862 | 0.045075117 |
| SFSWAP | 142.3143208 | -0.187883323 | 0.060295566 | -3.116038793 | 0.001832982 | 0.045195082 |
| EZH1 | 71.31075399 | -0.255177965 | 0.081901614 | -3.11566467 | 0.001835308 | 0.045195082 |
| TMEM94 | 121.1555766 | -0.278017015 | 0.089246509 | -3.115158447 | 0.001838461 | 0.045212438 |
| EEF1D | 424.096978 | 0.292744141 | 0.094001451 | 3.114251311 | 0.001844123 | 0.045291377 |
| SNORD1B | 6.631066891 | -0.652151078 | 0.209439852 | -3.113786944 | 0.001847028 | 0.045302473 |
| CALM2 | 422.2869425 | 0.227553376 | 0.073114189 | 3.112301164 | 0.00185635 | 0.045470731 |
| TES | 124.3963066 | 0.288051936 | 0.092582284 | 3.111307286 | 0.00186261 | 0.045563636 |
| DUSP23 | 86.22999939 | 0.251945546 | 0.081031677 | 3.109222919 | 0.001875801 | 0.045825628 |
| RPL14 | 128.5222725 | 0.276710504 | 0.089012628 | 3.108665706 | 0.001879342 | 0.045851483 |
| RERE | 330.6790073 | -0.185969437 | 0.059835948 | -3.107988455 | 0.001883654 | 0.045896056 |
| HSPD1 | 183.6846469 | 0.26128259 | 0.084080336 | 3.107535019 | 0.001886546 | 0.045905961 |
| IKBKE | 15.99201161 | 0.414650566 | 0.13348075 | 3.106444692 | 0.001893517 | 0.046014961 |
| SRD5A3 | 53.41344741 | 0.433974958 | 0.139763005 | 3.105077464 | 0.001902292 | 0.046106864 |
| SRP14 | 189.2932457 | 0.295608346 | 0.095198628 | 3.105174423 | 0.001901668 | 0.046106864 |
| SCN5A | 3.078343957 | -0.652936418 | 0.210405501 | -3.103228834 | 0.001914216 | 0.046335058 |
| EPG5 | 60.11862892 | -0.204067462 | 0.065814306 | -3.100655074 | 0.001930931 | 0.046678482 |
| TTYH3 | 78.74933431 | 0.413319175 | 0.13332404 | 3.100109905 | 0.001934488 | 0.046703356 |
| STMN3 | 18.82810297 | 0.539157062 | 0.173960913 | 3.099300019 | 0.001939785 | 0.046770086 |

| | | | | | | |
|---|---|---|---|---|---|---|
| FBXO34 | 43.1941825 | -0.251714204 | 0.081270146 | -3.097252988 | 0.001953231 | 0.047032886 |
| ARMCX5 | 18.29759956 | -0.344240215 | 0.111160374 | -3.096788921 | 0.001956291 | 0.047045237 |
| KRT81 | 187.5520212 | 1.116826145 | 0.360829807 | 3.095160446 | 0.001967065 | 0.047242803 |
| SCGB2A1 | 10.48567705 | -1.140720116 | 0.368617112 | -3.09459349 | 0.001970828 | 0.04727172 |
| SIX2 | 2.294343397 | 1.062282081 | 0.343565621 | 3.091933584 | 0.001988573 | 0.047285957 |
| TLR2 | 19.09515592 | 0.493976703 | 0.1596556 | 3.094014262 | 0.00197468 | 0.047285957 |
| MAP3K1 | 212.5392395 | -0.305238571 | 0.098712943 | -3.092183875 | 0.001986897 | 0.047285957 |
| GABRA1 | 2.7796016 | -0.790731185 | 0.255749539 | -3.091818611 | 0.001989344 | 0.047285957 |
| HLA-B | 1084.407628 | 0.414195194 | 0.133932714 | 3.092561804 | 0.001984369 | 0.047285957 |
| FBXO30 | 25.2750689 | -0.316914532 | 0.102490888 | -3.092123982 | 0.001987298 | 0.047285957 |
| ECH1 | 115.8009025 | 0.276208259 | 0.08931063 | 3.09266947 | 0.001983649 | 0.047285957 |
| TXNL4A | 122.8401567 | 0.228663826 | 0.073978365 | 3.090955379 | 0.001995136 | 0.047362685 |
| ELOVL1 | 100.8195085 | 0.235471071 | 0.07625351 | 3.088003061 | 0.002015064 | 0.047617476 |
| SREK1IP1 | 79.78392206 | 0.172510359 | 0.055867681 | 3.087838211 | 0.002016182 | 0.047617476 |
| AUXG01000058.1 | 25.73265002 | -0.43073264 | 0.139487603 | -3.087963586 | 0.002015332 | 0.047617476 |
| PDZD11 | 35.69208415 | 0.235334855 | 0.076206358 | 3.088126274 | 0.002014229 | 0.047617476 |
| IPP | 77.17149554 | -0.210934273 | 0.068382093 | -3.084641947 | 0.002037973 | 0.047911892 |
| MAP4K3 | 57.10021561 | -0.247338231 | 0.080167055 | -3.085285235 | 0.00203357 | 0.047911892 |
| SOCS5 | 32.06716397 | -0.279961264 | 0.090775373 | -3.0841103 | 0.002041619 | 0.047911892 |
| YBX1P2 | 4.823227165 | 0.581033876 | 0.18838809 | 3.084238907 | 0.002040736 | 0.047911892 |
| NPAS3 | 9.584332918 | -0.552979078 | 0.179243123 | -3.085078355 | 0.002034985 | 0.047911892 |
| AC024293.1 | 10.5842823 | 0.548482307 | 0.177999888 | 3.081363213 | 0.002060551 | 0.048294824 |
| CEBPB | 133.2534378 | 0.387152962 | 0.125667777 | 3.08076558 | 0.002064691 | 0.048330525 |
| AL590867.2 | 4.956441336 | 0.562184312 | 0.182663017 | 3.077712843 | 0.002085958 | 0.048643385 |
| ISG20 | 35.43599263 | 0.471527247 | 0.153204621 | 3.077761268 | 0.002085619 | 0.048643385 |
| DNTTIP1 | 32.02317656 | 0.232933817 | 0.075669846 | 3.078291135 | 0.002081914 | 0.048643385 |
| ZNF331 | 50.42930855 | -0.365029757 | 0.118637085 | -3.076860459 | 0.002091932 | 0.048659815 |
| RPL13AP7 | 5.442793726 | 0.468226733 | 0.152175585 | 3.07688473 | 0.002091762 | 0.048659815 |
| ERBB4 | 136.5563606 | -0.759496893 | 0.246930988 | -3.075745577 | 0.002099769 | 0.04874077 |
| USB1 | 21.84290597 | 0.375446033 | 0.122071865 | 3.075614788 | 0.002100691 | 0.04874077 |
| ADI1 | 103.2498453 | 0.295265411 | 0.096039302 | 3.07442271 | 0.002109104 | 0.048874573 |
| KMT5B | 153.7856803 | -0.198579218 | 0.064613239 | -3.07335185 | 0.002116688 | 0.048988851 |
| AFP | 2.955492604 | -1.585501319 | 0.516005246 | -3.072645734 | 0.002121702 | 0.049043448 |
| ARFIP2 | 108.455533 | -0.315808887 | 0.102795214 | -3.072213902 | 0.002124774 | 0.049053065 |
| RNU6-137P | 2.68469051 | -0.773102238 | 0.251701134 | -3.071508761 | 0.002129799 | 0.049107691 |

| | | | | | | |
|---|---|---|---|---|---|---|
| SETD2 | 249.641381 | -0.161465591 | 0.052599083 | -3.06974154 | 0.002142441 | 0.049216384 |
| SFT2D1 | 33.42671881 | 0.262387289 | 0.085475627 | 3.069732237 | 0.002142508 | 0.049216384 |
| RBIS | 108.0511863 | 0.2118362 | 0.06899441 | 3.070338612 | 0.002138162 | 0.049216384 |
| AC079140.2 | 6.092219881 | 0.700859589 | 0.22843612 | 3.068076925 | 0.002154412 | 0.049311882 |
| DOK2 | 19.34254921 | 0.567614056 | 0.185008664 | 3.06804041 | 0.002154675 | 0.049311882 |
| KLK12 | 1.409057424 | -2.457817196 | 0.801006332 | -3.068411693 | 0.002151999 | 0.049311882 |
| PIK3C2A | 210.2104924 | -0.189613188 | 0.06184327 | -3.066027832 | 0.002169231 | 0.049583578 |
| ZNF876P | 3.524722804 | -0.680119276 | 0.221854122 | -3.065614786 | 0.00217223 | 0.049590744 |
| KDM2A | 322.1672533 | -0.160930546 | 0.052503529 | -3.065137696 | 0.002175698 | 0.049608601 |
| CBY1 | 19.18073542 | 0.359230509 | 0.117222493 | 3.064518576 | 0.002180206 | 0.0496501 |
| C16orf54 | 8.729934344 | 0.6804076 | 0.222080719 | 3.063785105 | 0.002185558 | 0.049710688 |

**Table S3:** DE genes in White cases vs controls (n=812, P.adj < 0.05). Log2FC>0 = up in cases vs controls.
